# A Large-Scale Deep Normative Modeling of Primary Sulcal Patterns Reveals Deviations in a Spectrum of Disorders

**DOI:** 10.64898/2026.09.24.754179

**Authors:** Hyeokjin Kwon, Sarah U. Morton, Jane W. Newburger, Henry A. Feldman, Jong-Min Lee, P. Ellen Grant, Kiho Im

**Affiliations:** Fetal-Neonatal Neuroimaging & Developmental Science Center, Boston Children’s Hospital and Harvard Medical School, Boston, MA, USA; Division of Newborn Medicine, Boston Children’s Hospital, Boston, MA, USA; Department of Pediatrics, Harvard Medical School, Boston, MA, USA; Department of Cardiology, Boston Children’s Hospital, Boston, MA, USA; Biostatistics and Research Design Center, Boston Children’s Hospital, Boston, MA, USA; Department of Electronic Engineering, Department of Artificial Intelligence, and Department of Biomedical Engineering, Hanyang University, Seoul, South Korea; Department of Radiology, Harvard Medical School, Boston, MA, USA

**Keywords:** Primary sulcal patterns, Unsupervised anomaly detection, Normative modeling, Graph Neural Networks, Neurodevelopmental disorders

## Abstract

Primary sulcal patterns, the spatial arrangement of the earliest cortical folds, emerge prenatally and remain essentially stable after birth. Alterations are associated with cognitive outcomes and hold promise as clinical biomarkers. However, detecting abnormalities is challenging due to the high topological variability and complexity of normal individual sulcal patterns. Here, we introduce an unsupervised generative model that quantifies individual sulcal pattern deviations against a normative baseline learned from 10,349 typically developing subjects spanning early childhood through adulthood. We combine a generative reconstruction error with a discriminator trained on diffusion-generated pseudo-atypical graphs to leverage their complementary strengths in sensitivity and robustness. The discriminator captures subtle anomalies and is robust to demographic and data quality variations, whereas the generative model enables severity stratification and demonstrates cross-site generalizability. Across two clinical cohorts spanning subtle to overt sulcal abnormalities (congenital heart disease and polymicrogyria), our model captured global and localized condition-related deviations and linked them to neurodevelopmental outcomes. By enabling population-scale, individualized quantification of sulcal deviations, this framework provides a foundation for future clinical risk stratification.

## Introduction

The primary sulcal pattern, defined by the spatial arrangement and topology of the earliest cortical sulcal folds, emerges prenatally during the second trimester as a direct manifestation of emerging cortico-cortical connectivity^1–4^. Originating from neural stem cell proliferation and migration within the germinal zones, it reflects early functional specialization and remains remarkably stable after birth^5–7^, making it a candidate early biomarker of later neurodevelopmental outcomes^8,9^. Disruption of these patterns has been linked to various neuropsychiatric disorders and brain malformations^10–14^. Translating this into a clinically deployable marker, however, requires a generalizable, large-scale normative baseline to quantify individual deviation, which does not yet exist for sulcal patterns. Such a baseline would provide a foundation for the development of a novel biomarker for individual-level risk stratification, guiding targeted surveillance and early intervention.

Quantifying sulcal deviations is challenging due to the high multivariate complexity and profound inter-individual variability of cortical folding patterns. We previously developed a graphbased framework in which sulcal pits (putative primary folds) and their catchment basins serve as graph nodes, applying spectral graph-matching (SGM) to measure similarity between an individual and a predefined reference set of typical subjects^12,15–18^. However, this approach faces a critical bottleneck in scalability^19^. SGM evaluates an individual through pairwise graph comparisons against every reference pattern, requiring high-dimensional graph-alignment matrix construction for each pair. Consequently, the computational burden scales steeply with the number of reference subjects, rendering large-scale references intractable. This has confined prior studies to small, cohort-specific references, making the results potentially biased and less generalizable. Furthermore, SGM relies on manual feature weighting and treats all sulcal regions equally, limiting its sensitivity to localized sulcal alterations.

We previously showed that graph neural networks (GNNs) can model sulcal patterns to classify each subject from affected and unaffected groups^19^. However, such supervised frameworks depend on disease-specific labels and large patient cohorts, limiting generalizability. An unsupervised normative model capable of detecting sulcal pattern deviations across diverse clinical conditions without disease-specific training has not yet been explored.

We therefore introduce COMPASS (COMPosite Anomaly Scoring for Sulcal patterns), an unsupervised framework that quantifies sulcal deviations thorough two complementary ways. First, a generative graph autoencoder trained exclusively on typically developing (TD) patterns, learns to reconstruct them. It fails to reconstruct an unseen atypical graph, making the reconstruction error a direct indicator of sulcal deviations. However, such reconstruction error tends to overgeneralize and miss subtle deviations^20,21^. To recover sensitivity, we incorporate a discriminative component. Here, a diffusion probabilistic model synthesizes pseudo-atypical (PA) graphs, enabling a GNN to learn the boundary between TD and condition-associated graphs. COMPASS combines these two scores into a single composite index.

We train COMPASS on an unprecedented collection of 10,349 TD individuals spanning children to young adults. By absorbing the normative reference into the network at training, the model evaluates new subjects via a single forward pass, decoupling computational cost from reference scale to achieve sub-second inference per subject. We validate the clinical utility of COMPASS across two cohorts established in our previous studies: congenital heart disease (CHD) and polymicrogyria (PMG). This clinical cohorts serve to demonstrate model sensitivity and generalizability across diverse conditions. COMPASS efficiently differentiates patients from controls, localizes the regional alterations, correlates with neurodevelopmental outcomes, and outperforms traditional baselines including SGM. Leave-one-cohort-out (LOCO) and mixedeffects analyses show robustness across unseen data acquisition sites and to subject demographic and data quality variation. Together, these results establish COMPASS as a scalable foundation for individual-level neurodevelopmental risk stratification.

## Results

To model the normative distribution of typical sulcal patterns, we assembled a large-scale dataset of 10,349 T1-weighted brain magnetic resonance imaging (MRI) scans from 11 independent public cohorts^22–32^ (Methods, “Datasets”). For each MRI scan, three-dimensional cortical surfaces were reconstructed per hemisphere using FreeSurfer^33^, from which individual sulcal pattern graphs were extracted (Fig. 1a; Methods, “MRI processing”). We evaluated clinical utility of the model to two clinical cohorts (CHD^9,19,34,35^ and PMG^16^), allowing us to assess generalizability across diverse conditions.

**Figure 1.**
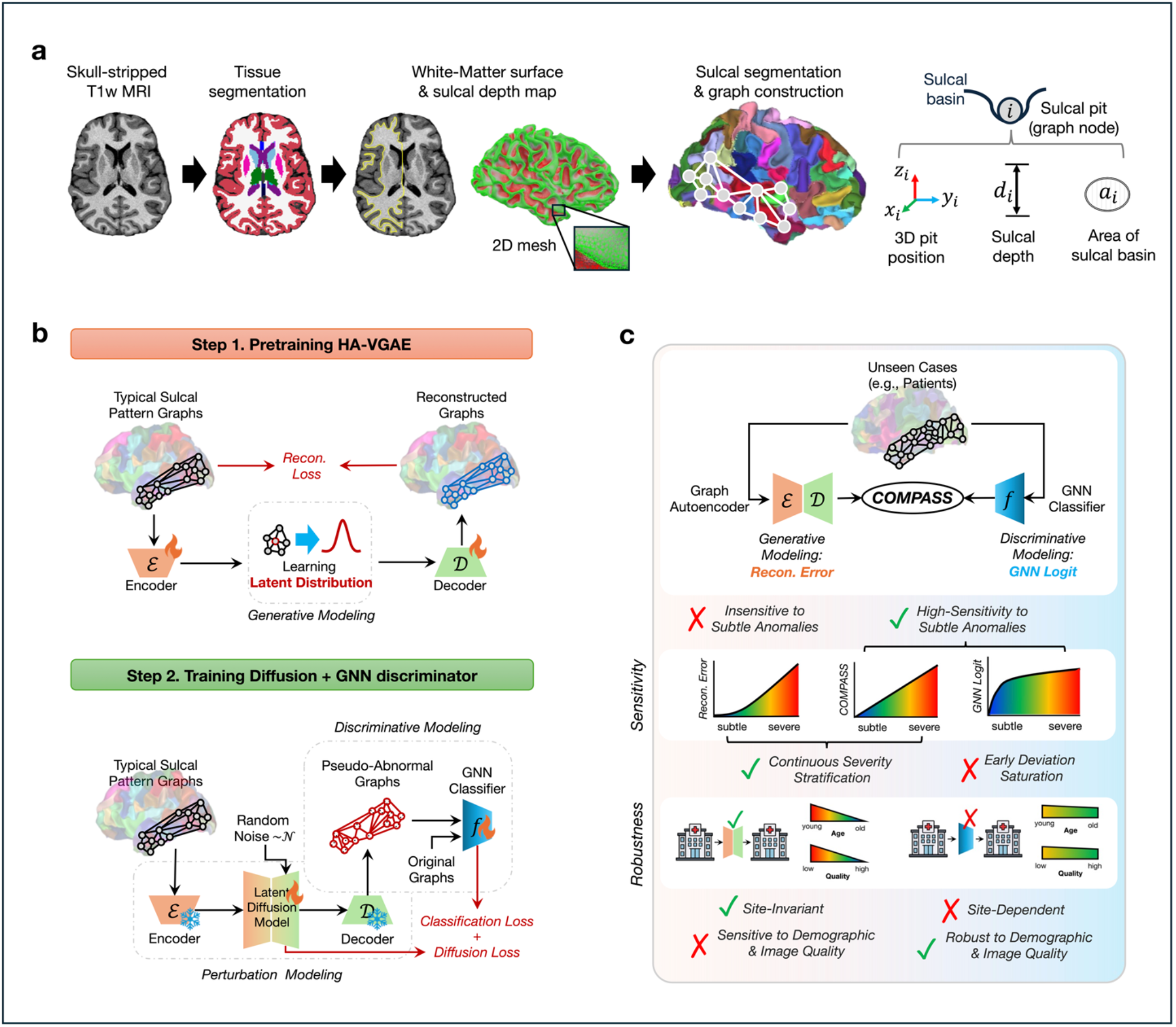
Overview of the COMPASS framework. COMPASS represents each sulcal pattern as a graph and scores its deviation from a large-scale normative baseline by combining a generative and a discriminative anomaly indices. (a) Sulcal pattern graph construction (Method, “MRI Processing”). From a skull-stripped T1-weighted MRI, tissue segmentation and white matter surface reconstruction yield a cortical surface and sulcal depth map, from which sulcal segmentation identifies sulcal pits and their catchment basins. Each pit becomes a graph node, linked to pits with adjacent basins. Each node carries three attributes: 3D pit position, sulcal depth, and basin area. (b) Two-stage normative model training. In step 1, a HA-VGAE with encoder ε and decoder *D* is pretrained on TD graphs to learn the normative distribution. In step 2, with the HA-VGAE frozen, a latent diffusion model perturbs encoded latents into PA graphs, and a GNN discriminator *f* is trained to separate these from the original graphs. This learns a normative decision boundary without real atypical data. (c) COMPASS inference and branch complementarity. For an unseen case, both frozen branches generate complementary metrics: the HA-VGAE-driven reconstruction error and GNN logits from the discriminative branch, combined into the composite COMPASS score. The two branches display complementary trade-offs across four functional axes. First, the GNN logit provides high sensitivity to subtle anomalies and robustness to demographic and data-quality variation, whereas the reconstruction error enables continuous severity stratification and robust cross-site generalizability. COMPASS combines both branches to achieve balanced reliability across the entire spectrum.

### Complementary Generative and Discriminative Approaches

Our first generative branch, a heterophily-aware variational graph autoencoder (HAVGAE), learns low-dimensional latent representations and reconstruct the input graphs (Fig. 1b). Trained exclusively on TD data, it yields high reconstruction errors on unseen atypical graphs, thereby quantifying deviation from the norm. However, these frameworks often suffer from overgeneralization by reconstructing subtle anomalies with unexpectedly high fidelity^36^, which blunts their detection sensitivity (Fig. 1c). To overcome this limitation, we synthesized PA graphs with controlled, low-intensity perturbations via a diffusion probabilistic model^37^ and trained a GNN discriminator to detect them. The resulting prediction score (“logit”) provides a sensitive discriminative deviation score^38^. However, because it is optimized for a binary (PA or not) decision boundary rather than to continuous deviation severity, the logit saturates as perturbation magnitude increases, reducing resolution for severe anomalies (Fig. 1c).

Evaluating the PA graphs across increasing perturbation magnitude *η* demonstrated a controlled spectrum of structural deviation. As *η* increased, sulcal pit positions shifted, inter-sulcal connectivity changed, and the morphological attributes, including sulcal area and depth, deviated smoothly from baseline values (Fig. 2a, first three columns). Critically, these perturbations degraded the graphs without violating the underlying graph topology (Fig. 2a, last column). We verified this via two Euler characteristics^39^ (Supplementary Note 1): the planar deficit, (3N − 6) – |E|, and the average node degree, both of which shifted measurably yet remained within valid planar topology up to extreme perturbations (*η* = 10). This fidelity is achieved by our latent semantic shifting (LSS) scheme, which applies anatomically constrained shifts within the learned normative manifold rather than adding random noise (Methods, “Latent Semantic Shifting”). A representative example illustrates the progressive transformation, where increasing *η* produces visible graph departures and a corresponding rise in predicted abnormality probability (Fig. 2b).

**Figure 2.**
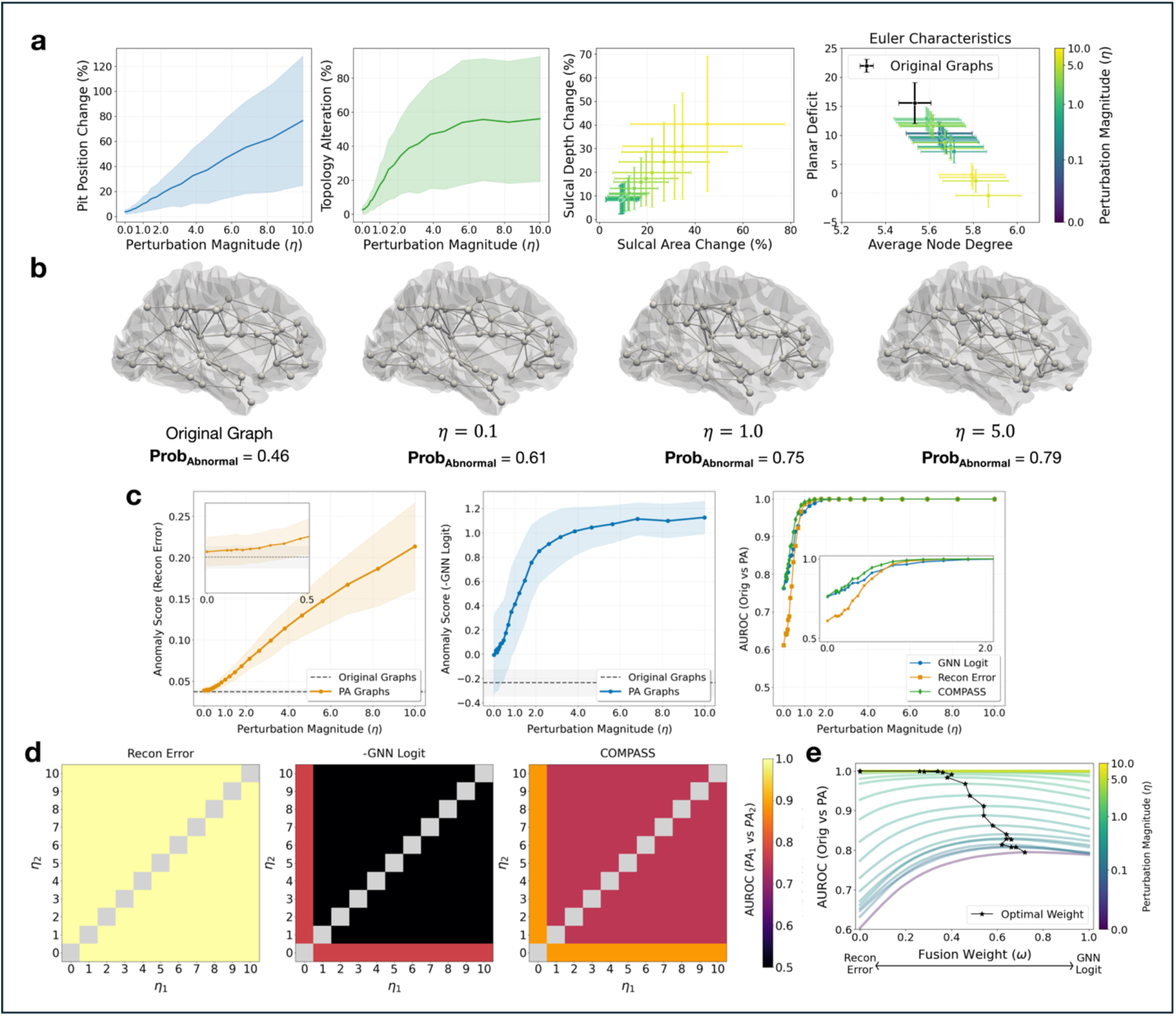
Complementarity of the generative and discriminative anomaly indices. (a) Synthesized PA graphs. As the perturbation magnitude *η* increases, the diffusion model produces a controllable spectrum of structural degradation: progressive change in sulcal pit position (as a percentage of mean edge length), graph topology (percentage of edges altered), and node attributes (sulcal depth and area). The rightmost panel shows two Euler characteristics, the planar deficit and average node degree (Supplementary Note 1). PA graphs stay within the range of the original graphs (black), departing from planarity only at the extreme *η* = 10. (b) Representative example. A single graph across increasing *η*, with the classifier’s abnormality probability (*Prob*_*abnormal*_) rising as the graph departs from its original form. (c) Behavior of the two indices. The reconstruction error rises approximately linearly with *η* but is insensitive and unresponsible at low intensities (left), whereas the GNN logit is sensitive early but saturates as *η* grows (middle). By AUROC against the original-graph baseline (right), each index leads in a different perturbation regime while their composite, COMPASS, tracks the stronger throughout. (d) Pairwise severity stratification. AUROC between PA graphs generated at each pair of perturbation levels (*η*1 *vs. η*_2_). The reconstruction error separates severity across all levels (left), the GNN logit only near the boundary (middle), and COMPASS combines both (right). (e) Fusion landscape. AUROC (original vs. PA) across fusion weights *ω* (Methods, “COMPASS”), colored by *η*. Optimal weights (black stars) shift from the GNN logit at low *η* toward the reconstruction error at high *η*, with equal weighting (*ω* = 0.5) near-optimal across most of the spectrum.

Empirical comparison of the generative reconstruction error and the diffusion-based GNN discriminator revealed clear trade-offs. Reconstruction error increased continuously with *η* (Fig. 2c, left), but consistently showed low sensitivity to subtle anomalies with low *η* (Fig. 2c, right). Conversely, the GNN logit provided precise detection at low perturbation magnitudes but saturated as *η* increased (Fig. 2c, middle). Pairwise comparisons between PA graphs across different perturbation levels (*η*_1_ *vs. η*_2_) further highlighted this saturation (Fig. 2d, middle). While the GNN logit separated original from PA graphs, it failed to stratify deviation severity, whereas reconstruction error provided continuous severity stratification.

These findings establish the primary axes of complementarity between the two branches: subtle anomaly sensitivity favoring the GNN logit, and severity stratification favoring reconstruction error. To combine these strengths, we introduced COMPASS by combining the two standardized discriminative and reconstruction indices (Methods, “COMPASS”). This composite metric matched or outperformed index alone in terms of area under receiver operating characteristic curve (AUROC) across all perturbation levels (Fig. 2c-d, right). Sweeping the fusion weight indicated that lower perturbations favored the GNN logit while severe ones favored reconstruction error, yet an equal weighting (*ω* = 0.5) achieved near-optimal performance across almost the entire *η* spectrum (Fig. 2e). We therefore adopt this parameter-free, equal-weight formulation as our primary metric, COMPASS_1. Empirically, a weight sweep across the clinical cohorts confirmed this choice, with performance peaking near *ω* = 0.5 for most conditions (Fig. 3a).

**Figure 3.**
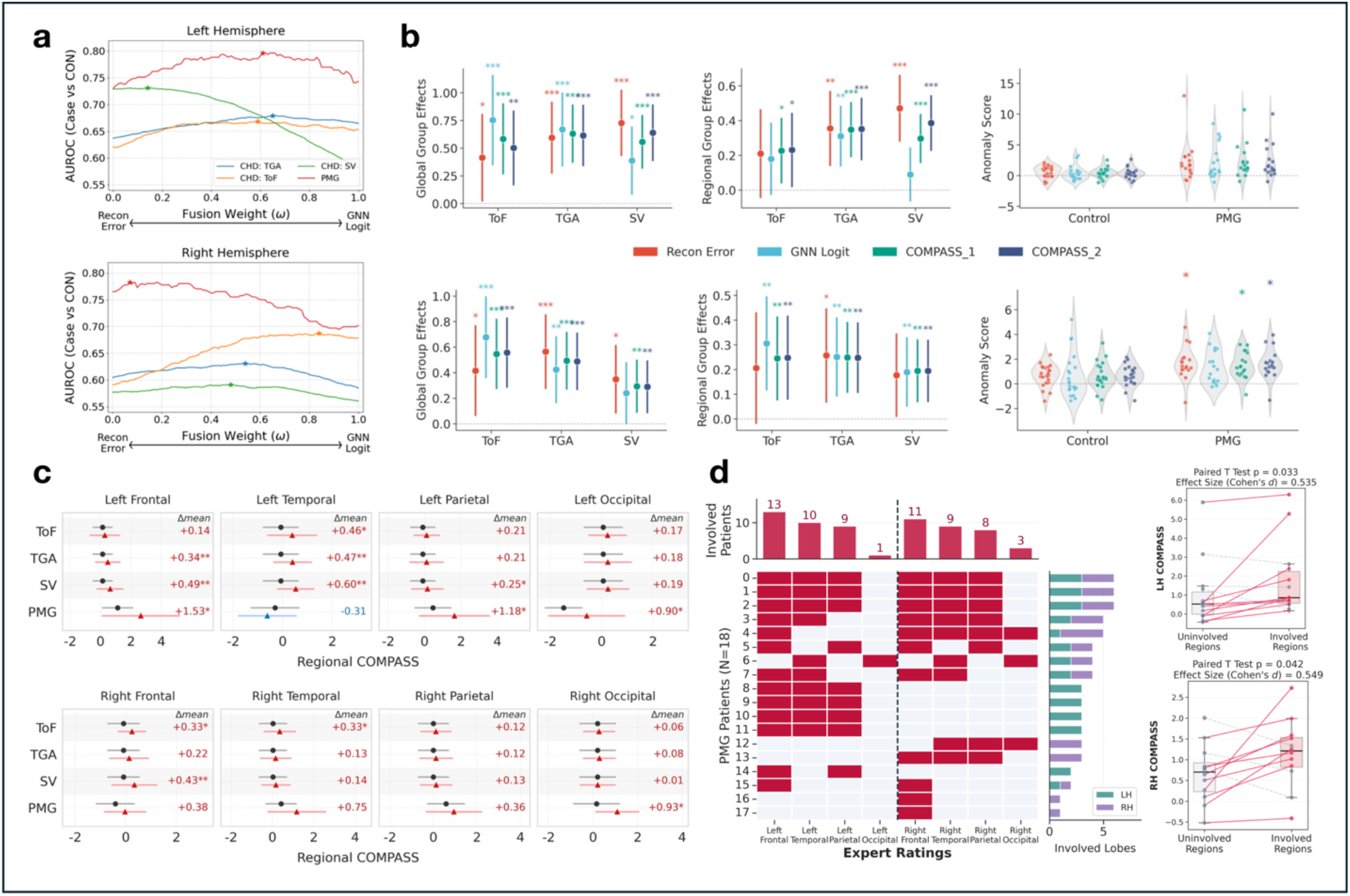
Clinical validation of COMPASS across three clinical cohorts. (a) Fusion landscape across cohorts. Case-control AUROC across fusion weights ω for each cohort, in the left (top) and right (bottom) hemispheres. Optimal weights (stars) cluster near ω = 0.5 for most cohorts, supporting equal weighting; LH of SV is a notable exception, leaning toward the reconstruction error. (b) Group effects across cohorts (top, LH; bottom, RH; colored by anomaly metric). For the CHD subtypes (ToF, TGA, SV), the first two sub-panels show global and regional group effects (standardized β with 95% CI) from ANCOVA and mixed models, respectively. The standalone reconstruction error and GNN logit each show subtype-specific blind spots that both COMPASS scores bridge. The right panel shows anomaly-score distributions for the PMG cohort versus controls. Asterisks denote significance (*q or p < 0.05, **< 0.01, ***< 0.001; see Methods). (c) Regional localization (top, LH; bottom, RH). Regional COMPASS group differences (mean difference with 95% CI; red triangles, patients; black circles, controls) across the four lobes per hemisphere, with the mean difference (Δmean) annotated. Each cohort shows a distinct, anatomically plausible spatial pattern of structural deviation. (d) PMG regional mapping. Heatmap (left) illustrates lobar involvement (manually rated by a pediatric neuroradiologist) sorted by the number of involved patients (columns) and lobes (rows), with top and right marginal bars showing lobe-wise and patient-wise totals, respectively. Paired intra-subject comparisons confirm significantly elevated COMPASS scores in involved versus uninvolved regions for both hemispheres.

### Clinical Validation in Congenital Heart Disease

We applied COMPASS to a CHD, the most common birth defect associated with subtle disrupted brain development and functional impairments, spanning three subtypes: tetralogy of Fallot (ToF)^9^, transposition of the great arteries (TGA)^35^, and single ventricle (SV)^34^. The standalone hemispheric (global) reconstruction error and GNN logit showed clear subtype- and hemisphere-dependent limitations in detecting patient-control differences (Fig. 3b, first column; Supplementary Table 1). Reconstruction error yielded relatively weak group separation in ToF (β = 0.414, 95% confidence interval [CI]: [0.017,0.812], *q* = 0.041 for left hemisphere [LH]; β = 0.416, 95% CI: [0.059, 0.774], *q* = 0.023 for right hemisphere [RH]), while the GNN logit showed more clearer group differences (β = 0.754, 95% CI: [0.345, 1.163], *q* < 0.001 for LH; β = 0.678, 95% CI: [0.357, 0.999], *q* < 0.001 for RH). The GNN logit performed poorly in SV, where its CIs approached zero (β = 0.387, 95% CI: [0.078, 0.696], *q* = 0.014 for LH; β = 0.240, 95% CI: [−0.002, 0.482], *q* = 0.052 for RH), whereas the reconstruction error yielded strong group separation (β = 0.728, 95% CI: [0.428, 1.028], *q* < 0.001 for LH; β = 0.348, 95% CI: [0.079, 0.618], *q* =0.017 for RH). In TGA, relative performance between the two metrics varied by hemisphere. The COMPASS_1 effectively eliminated these isolated blind spots, yielding consistently significant group separation across all CHD subtypes and both hemispheres (β = 0.558 to 0.631, all *q* < 0.001 for LH; β = 0.294 to 0.547, all *q* < 0.01 for RH). COMPASS_2, which uses cohort-optimized weights *ω* yielded comparable or marginally stronger separation as expected from in-sample tuning.

Lobe-by-subtype interactions were not significant (*p* = 0.245 for LH; *p* = 0.113 for RH), indicating no distinctive localized pattern across lobes. We therefore removed the interaction term. Across CHD subtypes, COMPASS maintained robust regional sensitivity while narrowing CIs (Supplementary Table 2). For example, in RH of the SV cohort, both standalone metrics produced attenuated regional effects with CIs reaching or crossing zero (β = 0.177, 95% CI: [0.008, 0.346], *q* = 0.060 for reconstruction error; β = 0.190, 95% CI: [0.048, 0.331], *q* = 0.009 for GNN logit. In contrast, COMPASS_1 (β = 0.195, 95% CI: [0.068, 0.322], *q* = 0.004) retained a robust and significant regional effect (Fig. 3b, second column). Adjusting for surface data quality proxy via the Euler number^40^ left both global and regional group separations unchanged (Supplementary Tables 3-4), indicating that the clinical signal is not a data-quality artifact.

Post-hoc regional comparisons revealed disease-specific spatial patterns that corroborate the established literature (Fig. 3c; Supplementary Table 5). ToF anomalies localized to the right frontal (RF) and temporal (RT) lobes, consistent with reported RH vulnerability^9^. SV deviations concentrated in the left temporal (LT) and parietal (LP) lobes, mirroring prior findings on the same dataset^34^. TGA, which lacks extensive isolated sulcal literature, showed novel left frontal (LF) and temporal disruptions.

### Clinical Validation in Polymicrogyria

In PMG cohort, the GNN logit failed to reach significance in RH (Fig. 3b, fourth column; Supplementary Table 6), consistent with logit saturation under severe perturbations far beyond the normative boundary, whereas both the reconstruction error and the COMPASS_1 captured these disruptions with robust effect sizes (Mean Diff = 1.732, 95% CI: [0.377, 3.086], *p* = 0.015 for LH; Mean Diff = 0.844, 95% CI: [0.166, 1.522], *p* = 0.016 for RH). Post-hoc regional comparisons showed widespread, highly significant deviations, most prominent in the LF, LT, LO, RT, and right occipital (RO) lobes (Fig. 3c; Δmean +0.90 to +1.53). To test whether COMPASS accurately captures regional alterations aligned with clinical evaluation, we conducted intrasubject paired comparisons between PMG-involved and uninvolved lobes, as manually rated by a pediatric neuroradiologist (Fig. 3d). These revealed that COMPASS scores were significantly higher in involved lobes than in uninvolved lobes (*p* = 0.033, Cohen’s *d* = 0.537 for LH, *p* = 0.042, Cohen’s *d* = 0.548 for RH). These results demonstrate that COMPASS effectively localizes local structural alterations in concordance with expert regional involvement ratings.

### Association With Neurodevelopmental Outcomes

To further assess the clinical significance, we correlated anomaly scores to cohort-specific neurodevelopmental outcomes (Fig. 4). In the CHD cohort, COMPASS showed stronger outcome associations than the standalone metrics (Fig. 4a, top; Supplementary Tables 7-10). Findings in RH aligned with prior reports^35^ on the same dataset: RT COMPASS_2 was associated with executive function (β = −0.429, 95% CI: [−0.723, −0.134], *q* = 0.035), and LF associations with processing speed and general memory corroborated earlier findings from an SV cohort within our dataset (β = −3.251, 95% CI: [−5.219, −1.284], *q* = 0.010 for processing speed; β = −2.529, 95% CI: [−4.692, −0.367], *q* = 0.088 for general memory)^34^. The LF-general memory association did not survive false discovery rate (FDR) correction, but its increased effect size relative to the reconstruction error (−2.465 to −2.529 for COMPASS_2) illustrates the added sensitivity. The previously reported RF-processing speed relationship (β = −3.411, 95% CI: [−5.574, −1.247], *q* = 0.017) was also replicated here (Fig. 4a)^35^. Fig. 4b confirms these patterns: higher anomaly scores, track a decline in the corresponding outcome (e.g., LF COMPASS_2 *vs*. processing speed and general memory). Adding the surface data quality proxy to the models, left the associations unchanged (Supplementary Tables 11-12), indicating the associations are not a data-quality artifact.

**Figure 4.**
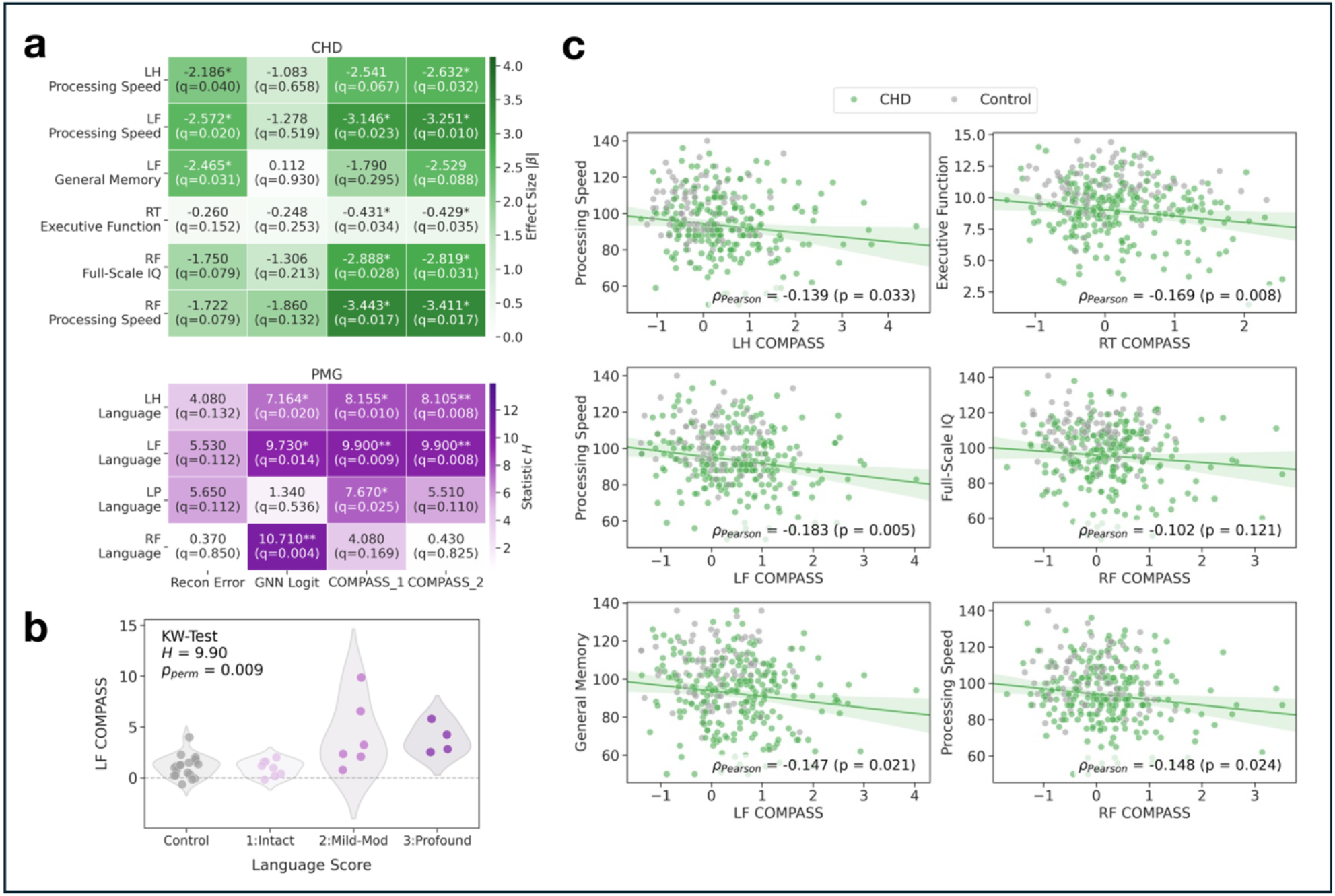
Associations between COMPASS scores and neurodevelopmental outcomes. (a) Association heatmaps for the CHD (green) and PMG (purple) cohorts. Rows are region-outcome pairs; columns are the four anomaly scores. Cells show the size of the association statistic (absolute standardized |β| for CHD, and Kruskal-Wallis *H* statistic for PMG; see Methods, “Statistical Analyses”) with its FDR *q*-value; asterisks denote *q < 0.05, **< 0.01, ***< 0.001. Across cohorts, the composite scores recover the broadest and strongest associations. (b) PMG distribution. LF COMPASS by ordinal language score (control, then intact, mild-to-moderate, and profound impairment), with the Kruskal-Wallis statistic and P value. Anomaly scores increase progressively with language-impairment severity. (c) CHD scatter plots. Global or regional COMPASS versus neurodevelopmental outcomes for representative region-outcome pairs (patients, green; controls, grey), with Pearson correlation coefficients *ρ* and P values. Higher anomaly scores track lower outcome scores.

In the PMG cohort, beyond a significant global effect in LH COMPASS_1 (*H* = 8.155, *q* = 0.010), the regional scores showed substantially stronger associations (Fig. 4a, bottom; Supplementary Table 13), with COMPASS_1 highest in LF and LP (*H* = 9.900, *q* = 0.009 for LF; *H* = 7.670, *q* = 0.025 for LP). In RF the GNN logit was strongest (*H* = 10.710, *q* = 0.004), suggesting that subtle topological folding differences there may bear on language function. In LF, where COMPASS was strongest (Fig. 4d), the scores stratified cleanly by severity: the COMPASS_1 distribution for the intact-language group (“1”) nearly matched healthy controls, while its distribution rose progressively from mild-to-moderate (“2”) to profound (“3”) impairment.

### Generalization and Robustness Across Cohorts

Although our prior works showed sulcal pit extraction is highly reproducible across sites and MRI sessions^18,41^, our multi-cohort design can still pose risks of over-fitting to specific sites and site-related batch effects. To evaluate cross-cohort generalizability, we used a LOCO validation design. For each target cohort, the model was retrained on the remaining ten cohorts. We then compared out-of-distribution (OoD) anomaly scores from the held-out cohort subjects against in-distribution (ID) scores obtained when all eleven cohorts were included in training (Fig. 5). Evaluating both scores on the exact same held-out validation subjects isolated site-level generalization while preventing data leakage (Supplementary Fig. 1).

**Figure 5.**
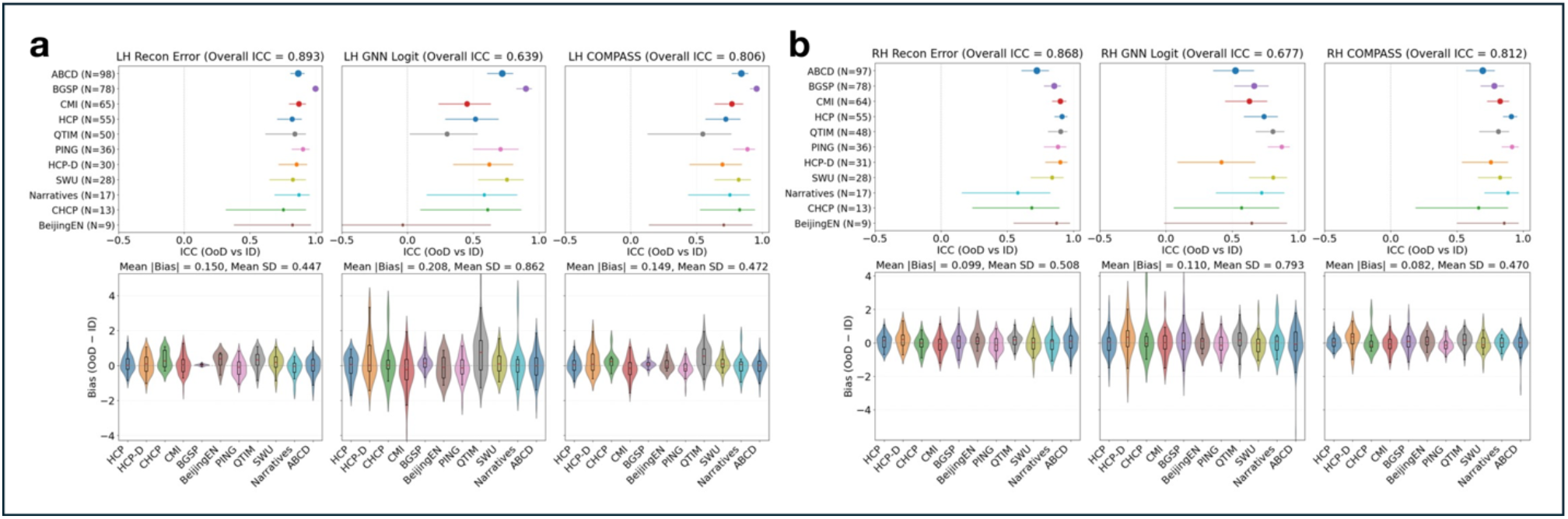
Generalization of COMPASS across cohorts. (a-b) Cross-cohort generalization under leave-one-cohortout (LOCO) validation, for the left (a) and right (b) hemispheres. Top: per-cohort intraclass correlation (ICC, with 95% CI) between out-of-distribution (OoD) and in-distribution (ID) scores, with the overall ICC in each title across anomaly scores. Bottom: per-cohort distributions of the OoD - ID bias, summarized by the mean absolute bias and SD. The reconstruction error transfers most stably to unseen cohorts and the GNN logit least, with COMPASS recovering most of the reconstruction error’s reproducibility

The reconstruction error showed high reproducibility between OoD and ID anomaly scores (overall intra-class correlation [ICC] = 0.893, mean absolute difference OoD - ID [|bias|] = 0.150 for LH; overall ICC = 0.868, |bias| = 0.099 for RH; Fig. 5a-b), indicating stable transfer across unseen sites. In contrast, the GNN logit was relatively less consistent (overall LH ICC = 0.639; overall RH ICC = 0.677) and exhibited long-tailed bias in several cohorts. COMPASS_1 recovered this lost stability (overall LH ICC = 0.806; overall RH ICC = 0.812), closely matching the reproducibility of the reconstruction error (see Supplementary Fig. 2 for visualization of correlations between OoD and ID scores, and Supplementary Tables 14-15 for full ICC and bias values). Surface data quality showed no significant effect on OoD bias (Supplementary Fig. 3, Supplementary Table 16).

### Demographic Trends in TD Subjects and Robustness to Batch Effects

We examined sulcal deviation variability across hemispheres, demographics, and image quality in TD subjects (Fig. 6a-b). In LH, the reconstruction error exhibited a modest negative age effect (β = −0.054, *q* = 0.001), which COMPASS attenuated to β = −0.032 (*q* = 0.018; Fig. 6a; Supplementary Table 17). This age-dependent trend may reflect heightened inter-individual variability and active maturation of cortex during preadolescent period. Although the normative training dataset includes substantial pediatric data (e.g., ABCD), active cortical remodeling in younger subjects introduces greater structural variance around the normative manifold, which gradually stabilizes into late adolescence^42,43^. Notably, combining the age-invariant GNN logit allows COMPASS to effectively attenuate this age drift of generative branch. Lower surface quality proxy correlated with higher anomaly scores (β = 0.219, *q* = 0.015 for the reconstruction error; β = 0.159, *q* = 0.033 for COMPASS; Supplementary Table 18). However, surface quality explained only a small fraction of the variance, leaving the age trend intact. Direct comparison between hemispheres revealed no significant differences in TD population across metrics (Fig. 6b; Supplementary Table 19). An age-by-sex interactions were not significant except for RH GNN logit (Supplementary Table 20), and sex main effects were not significant for any metric in either hemisphere.

**Figure 6.**
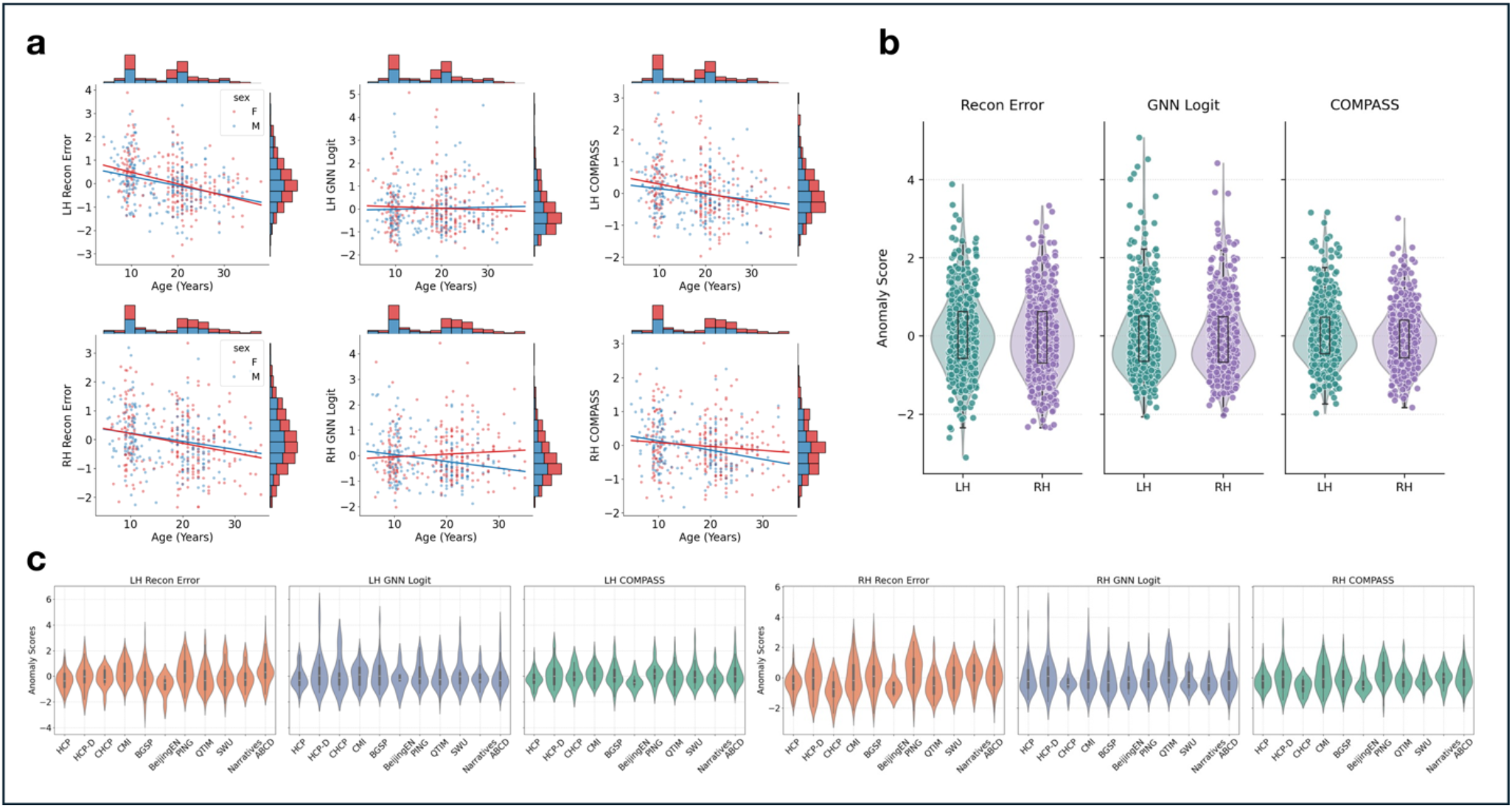
Cohort and demographic effects of COMPASS. (a) Age trajectories and sex differences: Scatter plots and marginal histograms showing the relationship between age (years) and anomaly scores (Recon Error, GNN Logit, and COMPASS) for the LH (top) and RH (bottom), stratified by sex (red: Female vs. blue: Male). Regression lines indicate the fitted age trends for each sex. (b) Anomaly score distribution for each hemisphere. Distribution comparison of anomaly scores between LH and RH across Recon Error, GNN Logit, and COMPASS, represented as combined violin and box plots. (c) anomaly-score distributions across the eleven normative training cohorts for each metric. (c) Anomaly score distributions across 11 independent normative training cohorts (HCP, HCP-D, CHCP, CMI, BGSP, BeijingEN, PING, QTIM, SWU, Narratives, and ABCD) for LH (left panels) and RH (right panels), demonstrating inter-site consistency and baseline stability around zero.

We next evaluated the potential batch effects across training cohorts (Fig. 6c). Unadjusted comparisons revealed significant cohort differences for the reconstruction error (*F* = 4.794 for LH; *F* = 5.272 for RH, both *q* < 0.001) and weaker differences for COMPASS (*F* = 2.005, *q* = 0.047 for LH; *F* = 2.133, *q* = 0.031 for RH), whereas the GNN logit showed no significant cohort differences (*F* = 0.880, *q* = 0.551 for LH; *F* = 1.699, *q* = 0.078 for RH). Post-hoc pairwise comparisons for the reconstruction error detected significant differences in 5 (LH) and 7 (RH) out of 55 comparisons, driven primarily by pediatric-adult contrasts such as ABCD *vs*. HCP. For COMPASS, only one pair (RH; CHCP *vs*. PING, *q* = 0.013) remained significantly different (Supplementary Tables 21-23). Adjusting for age and sex removed all cohort effects in LH across anomaly metrics, which remained unchanged after adding surface quality proxies. In RH, only the reconstruction error retained a residual cohort effect after adjustments (*F* = 3.222, *q* = 0.002 with age and sex; *F* = 3.482, *q* < 0.001 with age, sex, and quality), whereas COMPASS did not (Supplementary Table 24). These results indicate that observed cohort differences stem primarily from demographic differences rather than scanner- and site-related batch effects.

Together, these results demonstrate how the two branches complement each other across distinct dimensions of model robustness. LOCO validation measures cohort-level generalization, where the generative branch performs better. Conversely, demographic and quality analyses evaluate stability against age and image quality, where the discriminative branch excels. Because neither standalone branch satisfies both criteria, COMPASS provides a balanced solution by suppressing the generative branch’s demographic drift while preserving its cross-site reproducibility.

### Baseline Comparison

We compared COMPASS against the SGM, a one-class support vector machine (OCSVM)^44^ and our model’s two standalone branches. For a comparison purpose, patient-control group separation across clinical cohorts were performed (Fig. 7a; Supplementary Table 25). Clinical outcome associations were evaluated using Pearson correlation for CHD, and KW tests for ordinal PMG language scores (Fig. 7b; Supplementary Table 26).

**Figure 7.**
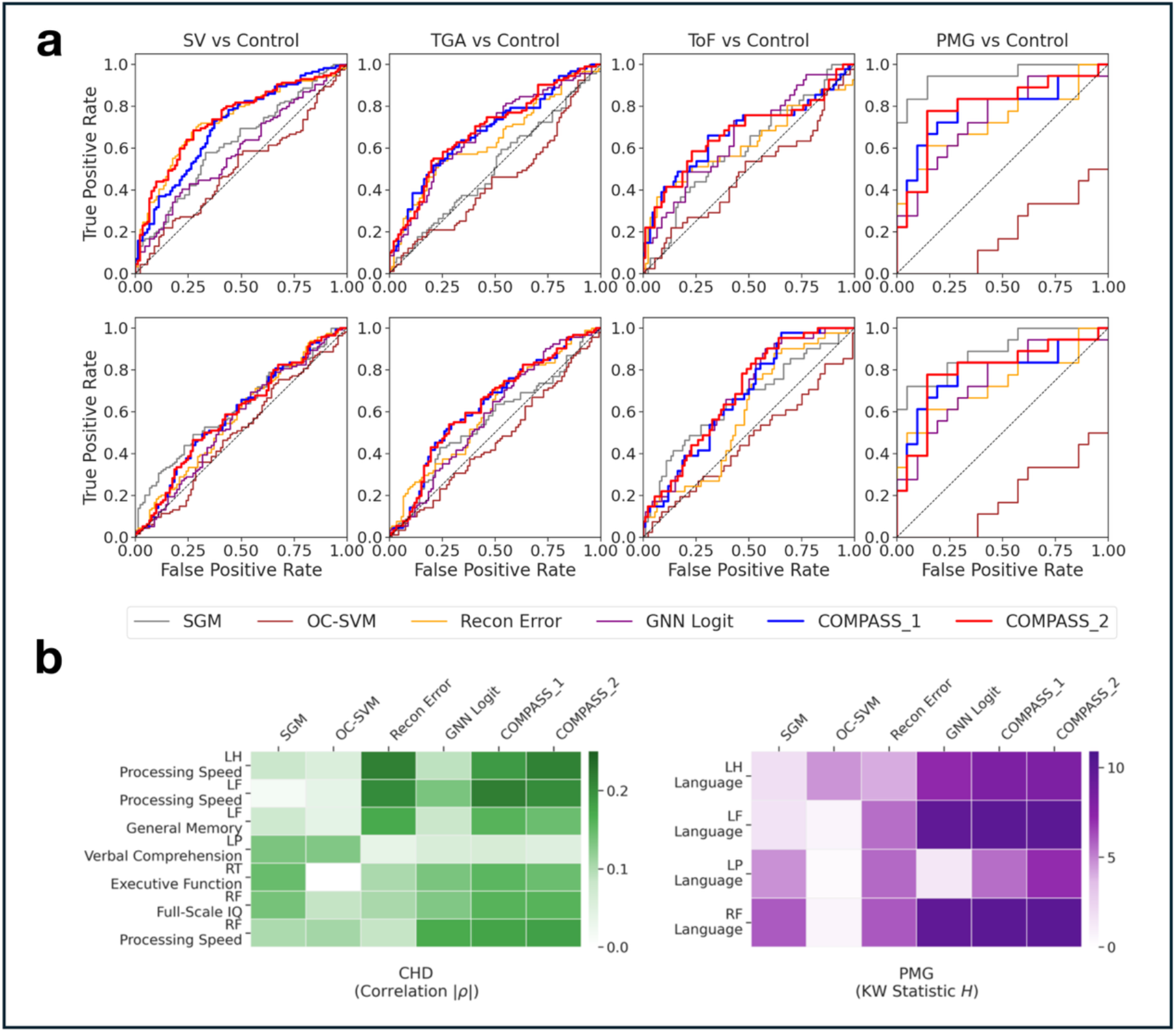
Comparison of COMPASS against baseline methods. (a) Case-versus-control discrimination. Receiver operating characteristic curves for each patient group versus controls (columns: SV, TGA, ToF, and PMG) in the left (top) and right (bottom) hemispheres, comparing six methods: SGM, OC-SVM, the reconstruction error, the GNN logit, COMPASS_1, and COMPASS_2. No single method is best everywhere, but COMPASS_1 ranks at or near the top across cohorts while each competitor fails somewhere in the panel. (b) Outcome associations. Strength of association between each method’s scores and clinical outcomes, for the CHD (green, Pearson |*ρ*|), and PMG (purple, Kruskal-Wallis *H*) cohorts; rows are region-outcome pairs reaching significance for at least one method. The composite scores recover the broadest and strongest associations across cohorts. Both panels use unadjusted statistics for a fair methodto-method comparison. SGM and COMPASS_2 carry an in-sample advantage, SGM through cohort-specific reference selection and COMPASS_2 through in-sample fusion-weight tuning, and should be read as favorable-case references.

Although no single method was uniformly best, COMPASS_1 demonstrated the highest consistency across all evaluated cohorts. The reconstruction error performed well on overt phenotypes, such as PMG and SV in LH, but lacked sensitivity to subtle anomalies. Conversely, GNN logit captured subtle deviations (e.g., ToF) while showing limited performance in SV. COMPASS_1 integrated the stronger branch in each regime, maintaining balanced sensitivity throughout. Among the external baselines, OC-SVM performed near or below chance across most cohorts and failed on PMG. SGM remained competitive, outperforming COMPASS_1 in three of ten comparisons, specifically both hemispheres of PMG (0.947 and 0.886 vs. 0.784) and RH of SV (0.620 vs. 0.591). Although SGM showed the strong performance in PMG, COMPASS_1 matched or exceeded SGM across the remaining subtle anomalies. Results were highly consistent across data splits and seeds in both hemispheres.

COMPASS_1 and COMPASS_2 identified the broadest set of significant outcome associations with the largest effect sizes across most cohorts. Crucially, the strongest associations detected by COMPASS 1 aligned with the core clinical hallmarks of each condition. These included RH executive function and LH processing speed in CHD, as well as LF language in PMG (COMPASS_1 *H* = 9.905, *q* = 0.007 *vs*. SGM *H* = 1.545, *q* = 0.462), both matching the core alterations reported in our previous studies^9,16,34,35^. The one exception was the association between LP region and verbal comprehension in CHD, detected by SGM and OC-SVM but missed by COMPASS variants. Overall, COMPASS_1 was the only metric that consistently ranked near the top across all evaluations without cohort-specific tuning, predefined reference, or heavy computational overhead. In contrast, other methods exhibited clear systematic weaknesses, such as OC-SVM’s geometric mismatch, the reconstruction error’s insensitivity to subtle deviations, the GNN logit’s instability, and SGM’s costly in-sample reference. This comprehensive reliability establishes COMPASS as a general-purpose normative metric.

## Discussion

We introduced COMPASS, an unsupervised normative framework for sulcal patterns that combines a deep generative model with a diffusion-based GNN discriminator. Trained on 10,349 TD sulcal patterns, it establishes an unprecedented scale of normative reference. Across two clinical cohorts (CHD and PMG), COMPASS detected condition-related global and local structural deviations and linked them to neurodevelopmental outcomes. By incorporating domain-tailored HA-VGAE and LSS perturbations, COMPASS effectively captured the complex manifold properties where conventional models fail.

Our central conceptual finding is that generative and discriminative branches are complementary across essential demands on normative modeling. First, the discriminative branch provides high sensitivity to detect subtle deviations and the generative branch stratifies deviation severity continuously. Second, the generative branch transforms more reliably across unseen data acquisition sites while the discriminative branch better resists demographic and quality variations. While neither standalone branch satisfies all demands, COMPASS combines their strengths, suppressing each weakness and providing balanced reliability.

Aggregating a normative population at this scale unavoidably introduce multi-site variation, which we evaluated through cohort comparisons, mixed-effects demographic modeling, and LOCO validation. Although age distributions varied across clinical cohorts (e.g., PMG), coupling reconstruction errors with age-invariant GNN logits effectively diluted non-pathological age drift. Cross-cohort differences were largely explained by biological age rather than site-related batch effects (e.g., scanner variability). Consequently, COMPASS achieved dual robustness, generalizing to unseen cohorts while remaining stable against demographic and data quality variations.

Several limitations define the path forward. First, the model was trained without explicit covariate conditioning. Although no sex main effects were observed in COMPASS, a modest age dependence persisted in LH, highlighting the need for age- and sex-conditioned normative modeling. Second, our developmental scope was restricted to individuals between 3 and 40 years of age, excluding early maturation and aging-related volumetric loss. Third, our evaluations relied on cross-sectional data, meaning longitudinal cohorts are required to establish predictive validity for long-term outcomes. Finally, the small sample size of the PMG cohort increased estimate variance. Larger evaluation cohorts will be necessary to stabilize effect estimates and resolve subtle regional alterations. Overall, COMPASS shows that large-scale normative modeling of sulcal patterns can detect and localize structural deviations and correlate with clinical outcomes, establishing a scalable foundation for future risk stratification and personalized treatment planning.

## Methods

### Datasets

To train the AI-based normative modeling of sulcal patterns, we used TD T1-weighted brain MRIs from eleven publicly available datasets (total n=10,349, age: 3-39 years): Human Connectome Project (HCP)^22^, HCP-development (HCPD)^23^, Chinese HCP (CHCP)^24^, Child Mind Institute (CMI)^25^, Beijing Normal University (BeijingEN)^26^, Brain Genomics Superstruct Project (BGSP)^27^, Pediatric Imaging, Neurocognition, and Genetics (PING)^28^, Queensland Twin IMaging (QTIM)^29^, Southwest University Longitudinal Imaging Multimodal (SWU-SLIM)^30^, Narratives collection^31^, and Adolescent Brain Cognitive Development (ABCD)^32^. To avoid potential confounding effect by aging-related volumetric loss^43^, we excluded subsets of participants over 40 years old (101 subjects from CHCP and 5 subjects from Narratives). To prevent overfitting to specific age range and to balance the age distribution across the total training dataset, we employed a sex-balanced, random subset of ABCD (1,000 males and 1,000 females). Supplementary Table 27 presents further details of demographic information for each dataset.

The first validation dataset consisted of a CHD cohort derived from three retrospective studies^9,19,34,35^. This cohort encompassed patients with specific CHD subtypes: SV (67 males, 48 females; age 14.8 ± 3.0 [mean±standard deviation (SD), years]), ToF (27 males, 14 females; age

14.8 ± 1.0), and TGA (70 males, 21 females; age 16.2 ± 0.7), alongside a healthy control group (46 males, 48 females; age 15.4 ± 1.9; Supplementary Table 29). To investigate the relationship between structural anomalies and clinical outcomes, this dataset included comprehensive neurodevelopmental assessments evaluating general intelligence, academic achievement, and executive functions (Supplementary Table 30).

The second clinical cohort is a PMG dataset^16^. This included 18 PMG patients (12 males, 6 females; age 8.8 ± 5.3), alongside an age-matched normative control group of 26 TD individuals (8 males, 18 females; age 8.9 ± 5.0; Supplementary Table 31). The ordinal language scores for 18 PMG patients were assessed based on language development evaluations (1: intact [n=8], 2: mild-to-moderate impairment [n=6], and 3: profound impairment [n=4]). Each lobar region across both hemispheres was visually evaluated by pediatric neuroradiologist (P.E.G) and categorized as involved or uninvolved (Fig. 3d).

All of these studies were approved by the Boston Children’s Hospital (BCH) institutional review board and written informed consent was obtained from the participants (aged more than 18 years) or their parents or guardians (aged <18 years). Further details regarding inclusion/exclusion criteria, MRI acquisition parameters, outcome assessment and patient factors were described in the previous studies.

### MRI Processing

We used FreeSurfer pipeline (v 8.1.0) to extract left and right white matter surfaces and aligned those surfaces to the standard MNI-305 space using the linear transformation, and visually inspected the output surfaces of each participant for accuracy (Supplementary Fig. 4)^33^. The sulcal pattern was represented using a graph structure with the primary sulcal folds as nodes, consisting of the sulcal pits and their surrounding catchment basins^15^. Cortical sulcal depth maps on the inner cortical surface were generated using FreeSurfer and were smoothed using a surface heat kernel with a full width of half-maximum of 10mm to prevent noisy and over-extraction of the sulcal pits. Sulcal pits and their sulcal catchment basins were identified using a watershed segmentation on the smoothed sulcal depth maps. Finally, the sulcal pattern graphs were constructed by connecting sulcal pits with graph edges, when their sulcal basins coincided. These graphs are inherently attributed graphs, which means that each graph node (sulcal pit) is attributed with geometric features: 3D position, and sulcal depth of the sulcal pit, the area of the sulcal basin. See Supplementary Fig. 5 for feature distribution as well as the number of sulcal pits and Euler planar characteristics (Methods, “Graph formulation and characteristics”) across training and evaluation cohorts used in this paper.

### Graph formulation and characteristics

Given the sulcal pattern graph ***G*** = (***V, E***) with a set of nodes 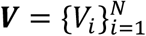 and edges ***E*** = ***V*** × ***V***, an adjacency matrix *A*_*i*_ ∈ {0,1}^*N*×*N*^ describe binary topology (1 for edges) of the graph ***G***. Each node *V*_*i*_ is associated with an attribute vector ***x***_*i*_ ∈ ℝ^5^, comprising of 3D sulcal position, sulcal depth, and area of sulcal basin. Node feature matrix is defined as 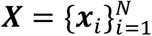. Node degree matrix ***D*** ∈ W^*N*×*N*^ is defined as diagonal matrix based on a node degree vector ***d*** = {*d*_*i*_}, where node degree indicates the number of neighboring nodes *V*_*j*_ ∈ ***ϕ***_*i*_ for each node *V*_*i*_ as: 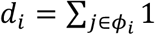.

Given the definition, sulcal pattern graph exhibits very unique, constrained geometric and topological characteristics^39^ (Supplementary Note 1). Furthermore, the sulcal pattern graphs also show distinct geometric feature behaviors. The most GNNs based on the common graph domains (e.g., social networks, molecular graphs, and functional brain connectivity) are designed based on the homophily principle^45^, meaning that the attributes of the connected node pairs are more likely to be similar. In sulcal pattern graphs, some features (3D positions) follow this principle, varying smoothly along the graph structure, such that adjacent nodes are spatially proximate. However, morphological attributes such as sulcal area and depth may differ substantially between neighboring pits. Consequently, sulcal graphs exhibit a geometrically low-homophily structure, characterized by locally homophilic spatial features but heterophilic morphological attributes.

### Model Overview

Our model consists of four components: (i) a graph autoencoder, (ii) a latent diffusion model, (iii) a perturbation conditioning module, and (iv) a GNN discriminator. Firstly, the graph autoencoder is pretrained to accurately map the graph to the low-dimensional, latent representation (encoder) and subsequently reconstruct the original graph using the representation (decoder). After that, the latent diffusion model perturbs the latent features, to generate PA patterns closely resembling the original ones. A GNN discriminator is trained to distinguish between original and PA graphs (reconstructed by the graph autoencoder using PA patterns), enabling the model to learn the decision boundary of normative sulcal patterns without relying on abnormal dataset.

### Heterophily-Aware Variational Graph Autoencoder

The graph autoencoder consists of graph encoder and decoder, where encoder maps node features to a latent space, from which a decoder reconstructs graph structure and attributes. Standard graph autoencoders^46^ have been widely used in various domains, but their underlying assumptions are not well aligned with the aforementioned geometric and topological characteristics of sulcal pattern graphs (Methods, “Graph formulation and characteristics”). To address this, we introduced modifications in both the encoder and decoder, proposing a HA-VGAE.

Conventional graph encoders rely on message-passing GNN, which learns node representations by gradually aggregating local neighboring nodes. In the spectral graph domain, this can be interpreted as low-pass graph filters under a homophily assumption^46^. However, sulcal pattern graph is associated with heterophilic, high-frequency features such as sulcal area and depth suggesting that the conventional message-passing GNN may suppress them, leading to suboptimal representation learning. To address this issue, we adopt a multi-band spectral GNN encoder, a Beta-wavelet GNN^47^. Specifically, the Beta-wavelet transform constructed by wavelets 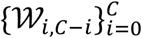 is defined given a node attribute vector ***x***:

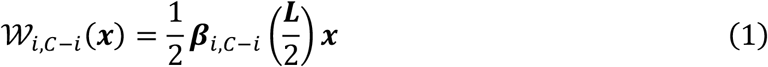

where ***β***_*i,C*-*i*_(***L***) is a Beta distribution over the normalized graph Laplacian ***L*** = ***D*** − ***A***. *C* is the wavelet order, determining the number of wavelets. Our HA-VGAE encoder computes the *c*-dimensional hidden embeddings ***H***_*i*_ ∈ ℝ^*c*×1^ for each wavelet, and then multi-layer perceptron (MLP) calculates the mean and log-variance of the latent node representations as follows:

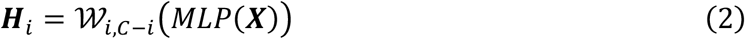

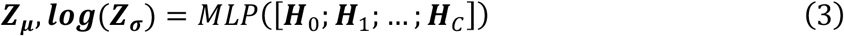

Finally, latent embeddings ***Z*** ∈ ℝ^*c*×1^ are then sampled as 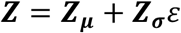, the standard reparameterization trick. *ε* ∈ *N*(0, *I*) is a random normal noise.

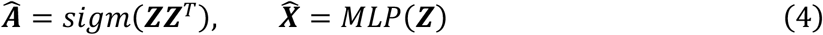

where Â ∈ {0,1}^*N*×*N*^ is a reconstructed adjacency matrix and *sigm*(·) denotes the sigmoid function. However, this strictly relies on homophilic assumption and may be ill-suited for reconstructing heterophily sulcal pattern features. Thus, HA-VGAE replaces it with an edge prediction layer that estimate the probability for each edge between nodes *i* and *j* as:

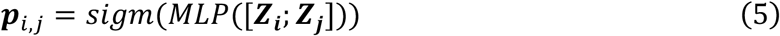

This formulation allows the decoder to model heterogeneous edges explicitly. For node reconstruction 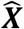, we retain the standard MLP-based decoder.

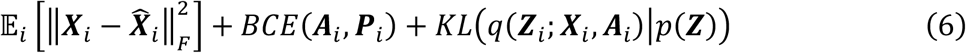

Where *i* denotes subject index, and 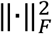 is a Frobenious matrix norm. *BCE*(·,·) means a binary cross-entropy loss function for edge reconstruction, and the last term represents a Kullback-Leibler (KL) divergence regularization for encouraging the approximated posterior *q*(***Z***_*i*_; ***X***_*i*_, ***A***_*i*_) toward the normal prior *p*(***Z***). See Supplementary Fig. 6 for the reconstruction performance of the trained HA-VGAE on the normative training dataset.

### Latent Diffusion Model

Given the structured latent node embeddings learned by the HA-VGAE framework, a latent diffusion model is employed to synthesize PA node embeddings via a continuous-time score-matching formulation. Following the idealized design space proposed by Karras et al. (2022)^37^, we parameterize the diffusion process such that the noise schedule directly dictates the perturbation level, setting σ(*t*) = *t*. In the forward diffusion chain, Gaussian noise is progressively applied to the initial clean latent state ***Z***_0_ = ***Z***, yielding the perturbed state ***Z***_*t*_ = ***Z***_0_ + *t****ϵ***, where ***ϵ*** ∈ *N*(0, *I*). The reverse generative trajectory is determined by the empirical Probability Flow Ordinary Differential Equation (ODE):

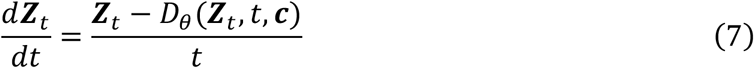

Where *D*_θ_ (***Z***_*t*_, *t*, ***c***) denotes the preconditioned denoising network designed to predict the clean target ***Z***_0_ from the noisy input ***Z***_*t*_. To stabilize training dynamics across widely fluctuating noise scales, *D*_θ_ is formulated using scale-activating preconditioning coefficients:

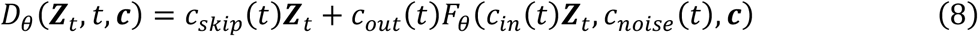

where *c*_skip_(*t*), *c*_out_(*t*), *c*_in_(*t*), and *c*_noise_(*t*) are functions of *t* that preserve unit variance throughout the network layers, and *F*_θ_ is the primary neural network.

### Latent Semantic Shifting

Usually, the perturbation condition ***c*** is introduced via a conditioning MLP τ to enforce controlled structural degradations^38^:

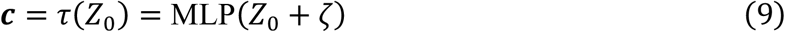

where *ζ* ~ *N*(0, *ηI*) represents a controlled stochastic modifier regulated by the perturbation intensity parameter *η*. While this previous perturbation model relying on independent, random Gaussian noise has demonstrated efficacy in non-medical graph domains (e.g., molecular or chemical graphs), sulcal pattern graphs exhibit unique topological constraints bounded by the planar surface manifold. Moreover, cortical folding develops through interdependent inter-sulcal relationships and underlying white-matter connectivity. Consequently, structural abnormalities in real clinical populations typically present as spatially interdependent abnormalities driven by disease-specific factors, rather than independent noise. For instance, CHD patients often exhibit localized reductions in sulcal area and depth within specific cortical regions due to restricted fetal blood supply and altered neurogenesis^9,34^. Relying entirely on pure Gaussian perturbation may lead to biologically unrealistic PA graphs, thereby limiting the model’s capacity to learn valid clinical decision boundaries.

To address this limitation, we propose LSS, a novel graph perturbation framework that integrates biological prior knowledge by inducing controlled, spatially clustered morphological shifts directly upon the learned normative manifold. We first define a stochastic regional mask *M* by randomly sampling anatomical lobes with a 50% probability per lobe. A global random perturbation direction Γ ∈ {−1,1} is sampled to dynamically dictate whether the target morphological features will be collectively magnified or diminished within the selected regions.

Using the differentiable decoder of the HA-VGAE, we reconstruct the intermediate node attributes to isolate the target structural features, specifically the sulcal area 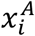 and depth 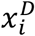. To enforce a coherent semantic drift, a semantic loss function *L*_*sem*_ is formulated to calculate the directed sum of these target features exclusively within the masked anatomical regions:

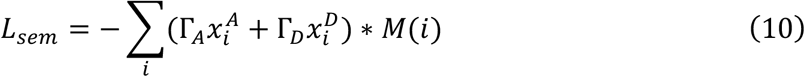

By computing the gradients of *L*_*sem*_ with respect to the input latent embeddings ***Z***, we obtain a precise directional vector that maximizes or minimizes the target features in the reconstructed graph space. The latent embeddings are subsequently shifted along this normalized gradient to generate biologically plausible structural variations without disrupting the overall topological validity:

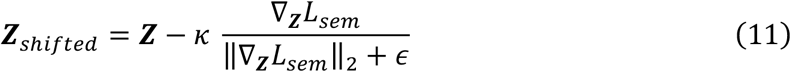

Here,κdenotes a scalar bounding the severity of the semantic shift, and ϵ is a minor constant to ensure numerical stability.

Following this directed semantic shift, a low-intensity, graph-regularized Gaussian noise is injected to introduce natural stochasticity and fully transform the embeddings into PA states. To ensure this unstructured noise respects the underlying cortical geometry, the raw Gaussian noise is smoothed via a random-walk aggregation across the neighborhood topology, yielding a localized noise tensor *ζ*_*smooth*_. The finalized perturbed embedding is computed as:

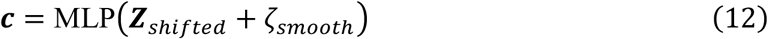

This multi-stage perturbation paradigm guarantees that the generated PA graphs present with biologically plausible, clustered structural anomalies while retaining sufficient topological stochasticity for robust anomaly classifier optimization. See Supplementary Fig. 7 for the example results of the LSS perturbation. To reconstruct topologically valid PA graphs from these synthesized PA embeddings, the perturbed latent representations are processed directly through the pre-trained HA-VGAE decoder. To construct a discrete and valid planar graph structure, the decoded edge probabilities *P* are thresholded to filter out low-confidence connections (excluding low prob < 0.05), followed by symmetric Bernoulli sampling to finalize the synthetic binary edges.

To test whether simpler graph perturbations could suffice, we evaluated heuristic and explicit perturbations, such as adding Gaussian feature noise or randomly rewiring edges. However, such unconstrained perturbations ignore the complex topology unique to sulcal patterns, generating unrealistic structures that risk reducing diagnostic sensitivity to real clinical deviations.

In contrast, our diffusion-based LSS operates directly on the learned normative manifold, which inherently encodes normative local sulcal morphology and inter-sulcal relationships, producing controlled and anatomically plausible anomalies. See Supplementary Note 2for further details.

### GNN Anomaly Discriminator

To establish a robust normative decision boundary, a graph-level GNN discriminator is optimized to distinguish between original graphs and generated PA graphs. For each input graph ***G***, the network processes node features to calculate a finalized graph-level anomaly score ***h***_***G***_. Given the input features, we employ a BWGNN, same as used in HA-VGAE encoder, to obtain the multi-scale node representation matrix ***H***_*node*_:

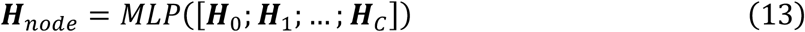

Rather than utilizing a standard global readout pooling (e.g., mean or sum pooling) that spatially dilutes focal structural breakdowns, we implement an anatomically constrained regional aggregation framework. For a specific anatomical lobe *k*, a localized regional embedding ***H***_*k*_ is calculated by computing the mean of its constituent node features:

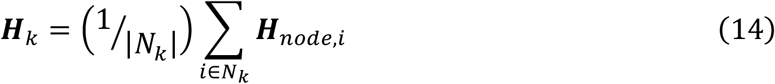

where *N*_*k*_ represents the set of nodes belonging to lobe *k*. The regional embedding matrix is then evaluated through a MLP layer to yield a set of regional anomaly logits *L*_*k*_ = *MLP*(***H***_*k*_).

To synthesize these individual lobar evaluations into a single graph-level anomaly score ***h***_***G***_ while explicitly prioritizing focal, localized cortical pathology, a differentiable LogSumExp (LSE) pooling operator is applied across the regional logits:

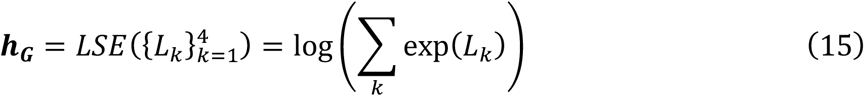

This regional logit aggregation guarantees that a severe structural anomaly confined to specific anatomical lobe(s) can effectively drive the global prediction score, thereby maximizing the anomaly detection sensitivity.

The diffusion denoiser and the downstream GNN discriminator are jointly optimized within an end-to-end framework. To align strictly with the ODE framework, the standard residual-noise prediction loss is replaced with a variance-weighted denoising score-matching objective. This formulation minimizes the expected L2 reconstruction error directly in the data space across a log-normal distribution of noise levels. The joint objective function combines a binary classification loss for anomaly detection with the score-matching loss for high-fidelity PA graph generation:

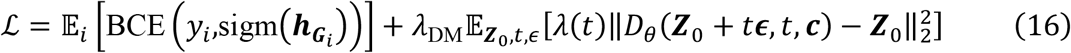

where *y*_*i*_ ∈ {0,1} represents the ground-truth graph label (1 for original normative graphs and 0 for generated PA graphs) for each subject *i*, and sigm(⋅) denotes the logistic sigmoid function mapping the graph-level discriminator outputs 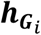. The term λ(*t*) is the continuous weighting function derived as 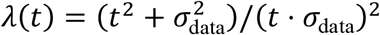 to ensure minimum variance in the score estimates, where σ_data_ represents the standard variance of the empirical latent dataset. The hyperparameter λ_DM_ serves as a regularization weight that balances the gradient contributions between the discriminative classification task and the diffusion generative modeling task.

### COMPASS

As described, COMPASS integrates two distinct indices: a generative anomaly score derived from the HA-VGAE and a discriminative anomaly score from the GNN discriminator. To establish the generative anomaly scores at the regional and global levels, we first define the reconstruction residual error for each individual node as follows:

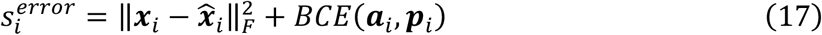

Where ***x***_*i*_ and 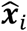 represent the original and reconstructed node feature vectors, respectively. ***a***_*i*_ and ***p***_*i*_ denote the ground-truth binary adjacency vector and predicted edge probability vector for node *i*.The node-level reconstruction residuals are subsequently averaged within a specific anatomical lobe *k* to yield the regional reconstruction error and averaged across the entire hemisphere to produce the global reconstruction error. Concurrently, the discriminative anomaly scores are directly extracted from the optimized GNN discriminator, yielding regional logits (-*L*_*k*_ in Eq. (15)) and a finalized global logit (-***h***_***G***_ in Eq. (15)).

To map these two functionally distinct modalities onto a standardized normative scale, we apply Z-score transformations to both the reconstruction errors and the GNN logits using the mean and standard deviation derived from the normative training dataset. These Z-scores quantify the statistical deviation of an individual’s sulcal pattern from the typical neurodevelopmental trajectory. Using those Z-scores, COMPASS combines the two indices by late fusion:

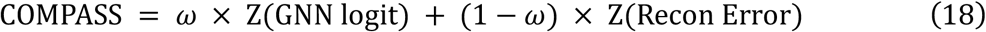

We adopt the parameter-free COMPASS_1 (*ω* = 0.5) as the primary metric in this study. This design was selected by systematically comparing three fusion levels against six composite scoring functions (Supplementary Fig. 8). The weight *ω* can also be determined in-sample (COMPASS_2), by maximizing case-control separation (AUROC) within the target validation cohort, and therefore represents an optimistic upper bound rather than an independently validated operating point. We report COMPASS_2 only as a sensitivity analysis bounding the additional separation that cohort-specific tuning could at most provide. This distinction also delimits their intended use: COMPASS_1 is hypothesis-free and applicable to prospective screening of single, unseen subjects, where cohort-level optimization is unavailable, whereas COMPASS_2 requires a formal casecontrol design in which the weight can be estimated and its optimistic bias explicitly acknowledged. Where a strong prior exists, *ω* may instead be set by hypothesis rather than fitted; for instance, lowering *ω* increases the relative contribution of the reconstruction error, which our simulations indicate is favourable for overtly severe malformation.

### Implementation Details

Models were implemented in PyTorch 2.4.1 and DGL 2.4.0. Pretraining and anomaly detection training together required approximately 2 hours on a single NVIDIA RTX A5000 GPU. The normative dataset was split into training and validation partitions comprising 95% and 5% of subjects, respectively, stratified across cohorts to preserve their relative proportions in both partitions.

The HA-VGAE was pretrained using the AdamW optimizer (batch size 24, weight decay 5e-04) for a maximum of 200 epochs, with an initial learning rate (LR) of 1e-02 reduced by a factor of 0.5 after 25 epochs without improvement, down to a minimum of 1e-04. The hidden dimension c and the wavelet order *C* were set to 32 (RH) and 64 (LH) and 3, respectively, and the KL-regularization term was weighted by 0.001. To prevent the network from learning an identity shortcut mapping, we imposed an additional information constraint on the latent space: the output dimension of the final MLP layer in Eq. (3) was restricted to 16, thereby limiting the information content of the latent vectors^36^.

The anomaly detection model was trained with AdamW (batch size 32, weight decay 1e-02) using two parameter groups with discriminative LRs: 1e-03 for the diffusion components and 1e-04 for all remaining GNN parameters. This asymmetry reflects the fact that the diffusion branch is trained from random initialization on a comparatively unconstrained objective, whereas the GNN encoder is simultaneously constrained by the classification and auxiliary losses and is therefore prone to destabilization under large updates. LRs followed a one-cycle schedule over 100 epochs, and to reduce sensitivity to the terminal point of the optimization trajectory we applied stochastic weight averaging over the final 25% of epochs^48^. The diffusion model used *T* = 25 time steps, a hidden dimension of 64, a perturbation noise scale of *η* = 0.05, and λ_DM_ = 0.01. The primary denoising network *F*_θ_ adopted the same MLP backbone as Karras et al. (2022)^37^, and the LSS parameter κ was sampled from a uniform distribution *U*(1.5, 2.5). The GNN discriminator comprised BWGNN layers with a hidden dimension of 32 and three wavelets (*C* = 2).

To mitigate the potential confounding effects arising from minor misalignments in cortical surface registration and overfitting to non-biological site-specific noise (Supplementary Fig. 9), we implement a domain randomization framework using data augmentation tailored for sulcal pattern graphs. Specifically, to explicitly simulate misalignment-driven batch effects and enhance model generalizability against spatial distortions, the global 3D coordinates of the sulcal pits undergo random rigid-body perturbations, including 3D rotations across yaw, pitch, and roll axes, as well as 3D translations along x, y, and z axes. The magnitude of these spatial perturbations is dynamically scaled according to the 3D position variances, ***σ***_*pos*_, calculated across the training dataset. The transformation parameters are randomly sampled from a uniform distribution spanning [0, 0.1* ***σ***_*pos*_], with an independent augmentation probability of 20% applied to each geometric direction. The final input features are subsequently projected into the initial hidden representation ***H***_**0**_.

See Supplementary Notes 3 and 4 for ablation analyses across alternative model configurations and hyperparameter settings.

### Baseline Methods

We compared COMPASS against well-established unsupervised baselines: (i) SGM, the previous method used for sulcal pattern analysis; (ii) OC-SVM, a widely used benchmark for unsupervised anomaly detection; and (iii) the stand-alone deep learning components of our own framework, namely the HA-VGAE reconstruction error and the diffusion-trained GNN discriminator.

SGM quantifies sulcal pattern normality as a mean similarity index (SI), obtained by comparing an individual graph against every graph in a predefined normative reference set^15^. Because this cost scales linearly with the size of the reference set, a strictly matched comparison, in which SI is computed against the same 10,349 normative graphs used to train COMPASS, would not practically feasible for the clinical cohorts evaluated here. We therefore used the previously published SI values for each cohort: CHD^35^ and PMG^16^. This unavoidable substitution introduces deviations from a like-for-like comparison, each of which favors SGM rather than our method. First, each cohort’s SI was computed against a cohort-specific reference set drawn from that cohort’s own healthy controls, giving SGM an in-sample reference advantage in precisely the casecontrol contrasts evaluated here, whereas COMPASS scores every subject against a single, fixed normative model that never saw these cohorts. Second, the reference sets differ in size and composition across cohorts, so SGM values are not strictly comparable between them. We therefore interpret SGM throughout as a favorable-case reference rather than a matched baseline. For OC-SVM, each graph was represented by the mean of its 16-dimensional latent node embeddings (Eq. 3) extracted from the HA-VGAE encoder, and the model was trained on exactly the same normative graphs used to train COMPASS.

### Statistical Analyses

All analyses were performed in Python v3.12.2 using statsmodels v0.14.4, scipy 1.14.1, and pingouin v0.6.1. Two-sided tests were used throughout, with significance set at 0.05. Where multiple comparisons were performed within a family of tests, p-values were corrected by the Benjamini-Hochberg FDR procedure, and corrected values are reported as *q*. The family over which correction was applied is specified for each analysis below. LH and RH were analyzed separately throughout.

In the CHD cohort, global anomaly scores were compared between each patient subtype and healthy controls using general linear model (GLM)-based analysis of covariance (ANCOVA), adjusting for age, sex, and socioeconomic status.

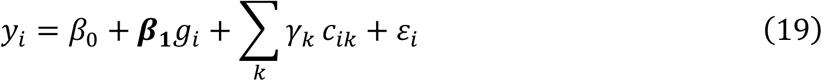

Where *ε*_*i*_ ~ *N*(0, σ^2^). For each subject *i, y*_*i*_ is global anomaly score, *g*_*i*_ is a gorup indicator, ***β***_**1**_ is the reported group effect, and k is the number of covariates *c*_*ik*_ (|*k*| = 3 for age, sex, and socioeconomic status). Regional scores were analyzed using mixed model for repeated measures (MMRM) with an unstructured covariance matrix, treating the four lobar regions within each hemisphere as repeated measures on each subject and including the same covariates.

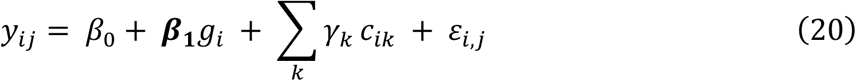

Where *ε*_*i,j*_ ~ *N*(0, *Σ*), *j* ∈ {frontal, temporal, parietal, occipital} and *Σ* is an unstructured 4×4 covariance matrix capturing correlation among lobes within each subject. A lobe-by-subtype interaction term was tested first and being non-significant in both hemispheres, was removed from subsequent models. Post-hoc regional comparisons were performed with independent two-sample t-tests. FDR correction was applied across subtypes within each analysis. The PMG cohort was assessed with independent two-sample t-tests (Fig 3c; global and local group differences) and paired t-tests and Cohen’s *d* effect size (Fig 3d; local intra-subject comparisons). Effect sizes are reported as standardized β for the GLM and MMRM analyses and as mean differences with 95% CIs for two-sample comparisons. As a sensitivity analysis, the CHD group comparisons were reestimated with the surface-quality proxy added as an additional covariate; this was restricted to CHD because the PMG cohort was evaluated with unadjusted two-sample tests that admit no covariates.

In the CHD cohort, associations between anomaly scores and neurodevelopmental outcomes were assessed with GLMs adjusting for age, sex, socioeconomic status, and cognitive test type (WISC or Wechsler Adult Intelligence Scale [WAIS], according to age). These models were likewise re-estimated with the surface-quality proxy as an additional covariate as a sensitivity analysis.

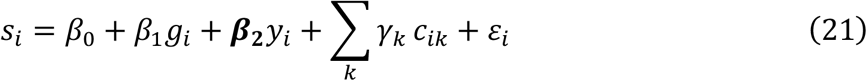

For each subject *i, s*_*i*_ is clinical outcome, ***β***_2_ is the reported association with anomaly score *y*_*i*_, and *k* is the number of covariates *c*_*ik*_ (|*k*| = 4 for age, sex, socioeconomic status, and cognitive test type). In the PMG cohort, the association between anomaly scores and the three-level ordinal language score was assessed with Kruskal-Wallis (KW) tests, as used in our previous study. FDR correction was applied across all region-outcome pairs within each cohort.

To assess generalization to unseen cohorts, we compared scores from models trained with and without each target cohort on a fixed evaluation set. Evaluation used the 5% validation partition, stratified so that every cohort contributes the same proportion of held-out subjects. The ID model was the main model, trained on the remaining 95% of all eleven cohorts. For each target cohort, a OoD model was retrained from scratch on the other ten cohorts only. Both were applied to the same validation subjects of the target cohort, giving paired ID and OoD scores per subject. See Supplementary Fig. 1 for further details.

Differences in anomaly scores across the eleven normative cohorts were assessed by oneway analysis of variance (ANOVA), followed by Tukey Honestly Significant Difference (HSD) post-hoc pairwise comparisons on the unadjusted scores. The analysis was then repeated as an ANCOVA adjusting for age and sex, and again with the surface-quality proxy added as a third covariate; cohort effects *F* were evaluated by type II sums of squares. FDR correction was applied across the anomaly metrics within each model specification. To quantify demographic and quality effects while accounting for residual between-cohort variability, linear mixed models (LMMs) were fitted with cohort as a random intercept.

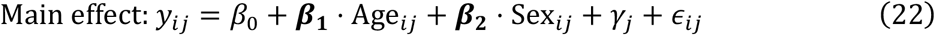

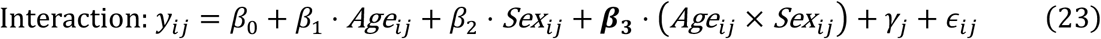

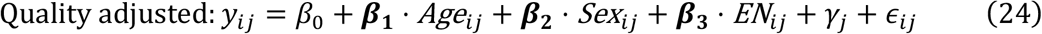

where *γ*_*j*_~*N*(0, τ^2^) is the random intercept for cohort *j*, ϵ_*ij*_~*N*(0, σ^2^) the residual, and *EN*_*ij*_ the surface data quality proxy (negative-log Euler number). An age-by-sex interaction was tested first, with Bonferroni correction across the three metrics within each hemisphere. Because the interaction was significant only for the GNN logit in RH, it was removed from the remaining models, and main effects of age and sex were estimated from the reduced specification. The surface-quality proxy was then added to these main-effect models to assess its contribution and its effect on the age coefficients. To test for hemispheric differences, a paired-sample LMM was implemented. To account for the within-subject correlation between LH and RH from the same individual, a subject-level random intercept was specified, while cohort-level variations were controlled via fixed-effect covariates:

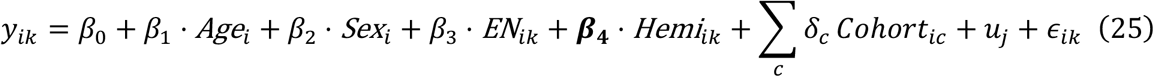

where *y*_*ik*_ denotes the anomaly score for subject *i* and hemisphere 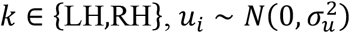 is the subject-level random intercept, Hemi_*ic*_ is an indicator for hemisphere (with LH as the reference), and *Cohort*_*ic*_ represents categorical variables for the cohorts. FDR correction was applied across all metric-predictor combinations within each model specification and hemisphere.

Baseline comparison was performed by using AUROC between each patient group and its controls, with bootstrap over 1,000 resamples. Outcome associations were quantified by unadjusted Pearson correlations for the continuous CHD outcomes, and KW statistics for the ordinal PMG language scores.

## Acknowledgements

This research was supported by the Thrasher Research Fund through the Thrasher Early Career Award Program.

