## Supplementary Information for "A Large-Scale Deep Normative Modeling of Primary Sulcal Patterns Reveals Deviations in a Spectrum of Disorders"

**Common Nomenclature**

GLM: General linear model
ANCOVA: Analysis of covariance

ANOVA: Analysis of variance
SV: Single ventricle
TGA: Transposition of the great arteries
ToF: Tetralogy of Fallot
SGM: Spectral graph matching
OC-SVM: One-class support vector machine
GNN: Graph neural network
COMPASS: Composite anomaly scoring system
FDR: False discovery rate
LH: Left hemisphere
RH: Right hemisphere
MMRM: Mixed model for repeated measures
LMM: Linear mixed model
PMG: Polymicrogyria

ND: Neurodevelopment

IQ: Intelligence Quotient

WISC: Wechsler Intelligence Scale for Children

WAIS: Wechsler Adult Intelligence Scale

TOWRE: Test of Word Reading Efficiency

KW-test: Kruskal-Wallis test

OoD: Out-of-distribution

ID: In-distribution

LOCO: Leave-one-cohort-out

ICC: Intraclass correlation

AUROC: Area under the receiver operating characteristic curve

QC: Quality control

MRI: Magnetic resonance imaging

HCP: Human Connectome Project

HCPD: HCP-development

CHCP: Chinese HCP

CMI: Child Mind Institute

BGSP: Brain Genomics Superstruct Project

PING: Pediatric Imaging, Neurocognition, and Genetics

QTIM: Queensland Twin Imaging

SWU: Southwest University Longitudinal Imaging Multimodal

ABCD: Adolescent Brain Cognitive Development

SES: Socioeconomic status

PA: Psuedo-Abnormal

MSE: Mean square error

MAE: Mean absolute error

BCE: Binary cross-entropy

GAT: Graph attention networks

GIN: Graph isomorphism networks

APPNP: Approximate personalized propagation of neural predictions

GPRGNN: Generalized PageRank graph neural networks

t-SNE: t-Stochastic Neighbor Embedding

**Supplementary Notes**

**1. Topological characteristics of sulcal pattern graphs.**

The main text treats the sulcal pattern graph as topologically equivalent to a planar triangulation and uses two Euler characteristics, the planar deficit and the average node degree, to verify that the synthesized PA graphs remain valid sulcal patterns (Fig. 2a). This note formalizes that equivalence, deriving how the pit-basin-adjacency construction yields a geodesic Delaunay triangulation and, from Euler's formula, the testable constraints |E| ≤ 3|V| − 6 and $\bar{d}<6$ used there ($\bar{d}$ denotes average node degree).

Let $\mathcal{M}\subset\mathbb{R}^{3}$ be a smooth, compact 2D Riemannian manifold representing the cortical surface mesh. Applying the watershed transform (Methods, “MRI processing”), local minima of the sulcal depth map $H\left( p \right)$are defined as sulcal pits, forming the finite set $\mathcal{P}=\{p_{1},..,p_{N}\}$, and it partitions the surface into basins each associated with pit $p_{i}$ (Supplementary Fig. 10A):

$$\begin{aligned} \mathcal{M}=\bigcup_{i} B_{i}, B_{i}\cap B_{j}=\emptyset for i\neq j\#\left( 1 \right) \end{aligned}$$

**Definition 1 (Geodesic Voronoi Diagram).** For a given pit set $\mathcal{P}$ on $\mathcal{M}$, the geodesic Voronoi diagram is:

$$\begin{aligned} Vor\left( p_{i} \right)=\{q\mathcal{\in M|}d_{\mathcal{M}}\left( q,p_{i} \right)\leq d_{\mathcal{M}}\left( q,p_{j} \right),\forall j\neq i\}\#\left( 2 \right) \end{aligned}$$

where $d_{\mathcal{M}}$is the geodesic distance on $\mathcal{M}$. Each watershed basin $B_{i}$ is equivalent to $Vor\left( p_{i} \right)$ under $d_{\mathcal{M}}$ (Supplementary Fig. 10B).

**Definition 2 (Sulcal Pattern Graph).** The sulcal pattern graph $G=(V,E)$ is the adjacency graph of the basins:

$$\begin{aligned} V={\{v_{i}\}}_{i=1}^{N}, \left( v_{i},v_{j} \right)\in E \mathrm{iff} \partial B_{i}\cap\partial B_{j}\neq\emptyset\#\left( 3 \right) \end{aligned}$$

Each node corresponds to a sulcal pit and edges indicate geodesic adjacency between basins.

**Lemma 1 (Duality).** The sulcal pattern graph $G$ is topologically equivalent to the geodesic Delaunay triangulation of $\mathcal{P}$:

$$\begin{aligned} G\cong{Del}_{\mathcal{M}}\left( \mathcal{P} \right)\#\left( 4 \right) \end{aligned}$$

This follows from the well-known dual relationship between Voronoi diagrams and Delaunay triangulations under the same metric, illustrated in Supplementary Fig. 10C-D.

**Theorem 1 (Topological Constraint for Realistic Graphs).** If $G$ is Delaunay-equivalent on a closed 2-manifold, it satisfies Euler’s formula:

$$\begin{aligned} \left| V \right|-\left| E \right|+\left| F \right|=2-2g\#\left( 5 \right) \end{aligned}$$

where $g$ is the genus of $\mathcal{M}$ ($=0$ for a hemispheric surface, topologically a sphere) and $F$ is a set of faces. Consequently,

$$\begin{aligned} \left| E \right|\leq3\left| V \right|-6,\mathrm{and}\bar{d}<6\#\left( 6 \right) \end{aligned}$$

These constraints define necessary geometric consistency conditions for any generated sulcal pattern graph. The planar deficit used in the main text, (3|V| − 6) − |E|, measures the slack in the first inequality; a non-negative deficit together with $\bar{d}<6$ certifies that a graph remains a valid genus-0 planar triangulation. Fig. 2a evaluates both quantities on the generated PA graphs, confirming that planarity is preserved across the clinically relevant perturbation range and breaks down only under extreme, non-physiological settings.

**2. Comparison against explicit perturbation strategies.**

For the GNN disctimative branch, our framework generates PA graphs by sampling from a conditional latent diffusion model (Methods, “Latent Diffusion Model”). A reasonable question is whether this added complexity is necessary. One could instead corrupt node features with Gaussian noise directly, optionally combined with random edge rewiring. We implemented two such baselines and compared them against the proposed method under matched conditions. All three strategies use the same upstream pipeline (lobar masking, and random-walk smoothing of the perturbation). They differ only in how the PA graph is constructed, as follows:

**Diffusion (proposed).** The shifted latent embedding is used as a conditioning input, and the PA embedding is generated by reverse diffusion process, then decoded through the frozen HA-VGAE decoder.

**Node.** No diffusion sampling. Gaussian noise of magnitude $\eta_{node}$ is added directly to the features of masked nodes. Topology is unchanged.

**Node+Edge.** As above, with an additional random removal and re-allocation of a fraction $\eta_{edge}$ of undirected edges. Candidate edges are restricted by empirically chosen Euclidean distance threshold (<1.5) so that rewiring does not create anatomically implausible long-range connections. Because this adds a second free parameter, $\eta_{node}$ was fixed at the best value from the **Node** setting in each hemisphere (LH $\eta_{node}=1$, RH $\eta_{node}=5$).

Sulcal pits do not vary independently. The position, area, and depth of each pit are constrained by those of its neighbours, and the HA-VGAE learns this complex structure from large-scale data. Sampling within the learned latent space lets us perturb a graph while preserving these dependencies. LSS moves the representation along the latent manifold in a semantically directed way, and a small conditioning noise ($\eta= 0.05$) keeps the sampled latent close to the region the decoder was trained on. The resulting PA graphs are altered in the intended respect but still resemble real sulcal patterns, which is what makes them useful as a effective negative class. In contrast, explicit perturbation strategies cannot achieve this. Corrupting node features directly does not produce the corresponding change in topology, and rewiring edges at random does not produce the corresponding change in features. The two are decoupled, so the resulting graphs are less plausible as anatomical variants.

Our t-SNE analysis shown in Supplementary Fig. 11a shows this directly^1^. Diffusion-generated PA embeddings of masked nodes remain interleaved with the original distribution, whereas **Node** and **Node+Edge** embeddings spread into regions where real data are sparse. Supplementary Fig. 11b further hows the quantitative result. The diffusion strategy performs best at small $\eta$ (mean AUROC 0.712 ± 0.065 for LH; 0.677 ± 0.065 for RH) and stays above both explicit baselines across the low perturbation range in both hemispheres. Its advantage narrows at large $\eta$, which is consistent with strong conditioning noise pushing samples away from the learned latent region, where the generative prior no longer constrains the output. As the error bars of the three strategies overlap substantially, explicit perturbation might remain reasonable as a form of domain randomisation. Training on a wide range of corrupted graphs can still yield a usable decision boundary. Within a cohort of 10,349 subjects, however, PA graphs that stay close to the data manifold provided a more effective training signal, and did so consistently across perturbation levels and both hemispheres.

**3. Sensitivity analyses to model configuration and hyperparameters.**

To verify that our findings do not depend on a particular architectural or hyperparameter choice, we systematically varied each major component of the framework and re-evaluated case-versus-control discrimination. Each configuration was trained from scratch and evaluated by the mean AUROC across all clinical cohorts, with uncertainty estimated by 1,000 bootstrap resamples; the two hemispheres were modeled and evaluated separately. All other settings were held at their final values while a single factor was varied (Supplementary Fig. 12).

We probed four groups of factors. (i) HA-VGAE pretraining (Methods, “Heterophily-Aware Variational Graph Autoencoder”): the encoder architecture (comparing our BWGNN against GraphSage, GAT, GIN, APPNP; Eq. (2) and (3); see Supplementary Note 4 for details), the edge decoder framwork (conventional inner product vs. our edge predictor; Eq. (5)), and the three principal BWGNN hyperparameters, the hidden dimension $c$, the bottleneck (latent) dimension, and the wavelet order $C$. (ii) Input augmentation (Methods, “Implementation details”): the presence or absence of the proposed random rigid-body rotation and translation of sulcal pit coordinates. (iii) Perturbation and diffusion (Methods, “Latent Diffusion Model” and “Latent Semantic Shifting”): the number of diffusion time steps $T$, the hidden dimension of the denoising network $F_{\theta}$, the weight of the diffusion loss $\lambda_{\text{DM}}$ (Eq. (16)), and the two LSS parameters governing perturbation magnitude $\eta$ (Eq. (9)) and the semantic-shift step size $\kappa$ (Eq. (11)). (iv) Anomaly classifier (Methods, “GNN Anomaly Discriminator”): the classifier backbone, again comparing BWGNN against standard message-passing and heterophily-friendly alternatives (Supplementary Notes 4), together with its hidden dimension and wavelet order.

Across all factors, performance varied within a narrow band and no single choice altered the qualitative conclusions. Two patterns were nonetheless consistent. First, the spectral BWGNN encoder and classifier outperformed conventional message-passing alternatives, consistent with the mixed-homophily structure of sulcal pattern graphs described in Methods: architectures built on the homophily assumption suppress the heterophilic morphological attributes that carry much of the anomaly signal. Second, removing the rigid-body augmentation reduced performance, supporting its role in mitigating residual registration misalignment (Supplementary Fig. 9). The remaining hyperparameters produced only modest variation, indicating that the framework is not finely tuned to any specific setting.

**4. GNN baselines for sulcal pattern graphs.**

To assess whether our spectral encoder and classifier are justified, we compared the BWGNN against alternatives: conventional message-passing architectures, which are the standard baselines across graph learning, and architectures explicitly designed for heterophilic graphs.

**Message-passing baselines.** GraphSAGE^2^, GAT^3^, and GIN^4^ all follow the message-passing paradigm, in which each node's representation is iteratively updated by aggregating the representations of its neighbors. GraphSAGE applies a permutation-invariant pooling operator (mean, max, or LSTM^5^; we used mean aggregator in this study) over sampled neighbors. GAT weights neighbors by learned attention coefficients. GIN uses a sum aggregator followed by a MLP, chosen to maximize discriminative power with respect to the Weisfeiler-Lehman isomorphism test. They share a common inductive bias: repeated neighborhood averaging acts as a low-pass filter on the graph signal^6^, smoothing node features toward their local neighborhood mean. This is well matched to homophilic graphs, in which connected nodes tend to share similar attributes.

However, sulcal pattern graphs violate this assumption. As described in Methods “Graph formulation and characteristics”, they are partially homophilic: the sulcal pit coordinates vary smoothly along the graph, since adjacent basins are by construction spatially proximate, but the morphological attributes do not. Sulcal depth and basin area can differ substantially between neighboring pits, reflecting the local variability of cortical folding. Because low-pass aggregation suppresses precisely the high-frequency component of the signal, message-passing encoders tend to average away these morphological contrasts, which is problematic here because subtle deviations in depth and area are a principal carrier of the anomaly signal. In both the HA-VGAE encoder and the anomaly classifier, replacing BWGNN with GraphSAGE, GAT, or GIN reduced case-versus-control AUROC (e.g., 0.712 to 0.590 - 0.683 for LH classifier; 0.677 to 0.600 - 0.667 for RH classifier; Supplementary Fig. 12).

**Heterophily-oriented baselines.** We additionally compared three architectures developed specifically to handle heterophilic graphs, providing a stricter test than the message-passing baselines. APPNP^7^ decouples feature transformation from propagation, where node features are first transformed by a MLP and then propagated by a personalized PageRank iterations, in which a teleport probability $\alpha_{PR}$ retains a fixed fraction of each node's own representation at every propagation step. This restrains over-smoothing and preserves node-level information across hops, but the propagation weights remain fixed and non-negative, so the operator is still fundamentally low-pass. GPRGNN^8^ generalizes this by learning the propagation coefficients of a generalized PageRank expansion. Because these coefficients may take negative values, the model can in principle realize high-pass as well as low-pass filters and thus adapt its spectral response to the graph at hand. H2GCN^9^ instead modifies the architecture directly, combining three designs shown to help under heterophily: separating a node's own embedding from those of its neighbors rather than mixing them, aggregating over higher-order (order $K_{h2gcn})$ neighborhoods so that structurally similar but non-adjacent nodes can inform one another, and concatenating representations from all intermediate layers rather than using only the final one.

These architectures narrowed but did not close the gap to BWGNN (Supplementary Fig. 12). We interpret this as reflecting the particular structure of sulcal pattern graphs, which are neither homophilic nor heterophilic as a whole but mixed across feature types. Methods that adapt a single global filtering behavior, whether fixed as in APPNP or learned as in GPRGNN, must trade one regime against the other. A multi-band spectral decomposition instead retains low- and high-frequency components in parallel, allowing the smooth spatial signal and the contrastive morphological signal to be represented simultaneously.

**Baseline configuration.** To ensure a fair comparison, all baselines were matched to our model in receptive-field depth and tuned over their principal architecture-specific hyperparameter. Because BWGNN was used with a wavelet order of C = 2 for GNN discriminator, and 3 for HA-VGAE, giving three to four filters including the low-pass band, all message-passing baselines (GraphSAGE, GAT, GIN) were likewise given three to four propagation layers, so that every architecture aggregates information over a comparable neighborhood radius. For GAT, the number of attention heads was varied over 1, 2, and 4, and 2 was selected. GIN used the standard two-layer MLP in its aggregation step. For APPNP and GPRGNN, we had no prior expectation for the teleport probability $\alpha_{PR}$ and therefore evaluated 0.1, 0.2, and 0.3, selecting 0.1; the number of propagation steps $K_{PR}$ was fixed at the standard value of 10. For H2GCN, the neighborhood order was fixed at the standard $K_{h2gcn}$ = 2. All remaining settings, including the hidden dimension, optimizer, and training schedule, were held identical to those of our model (Methods, "Implementation details").

**Supplementary Figures**

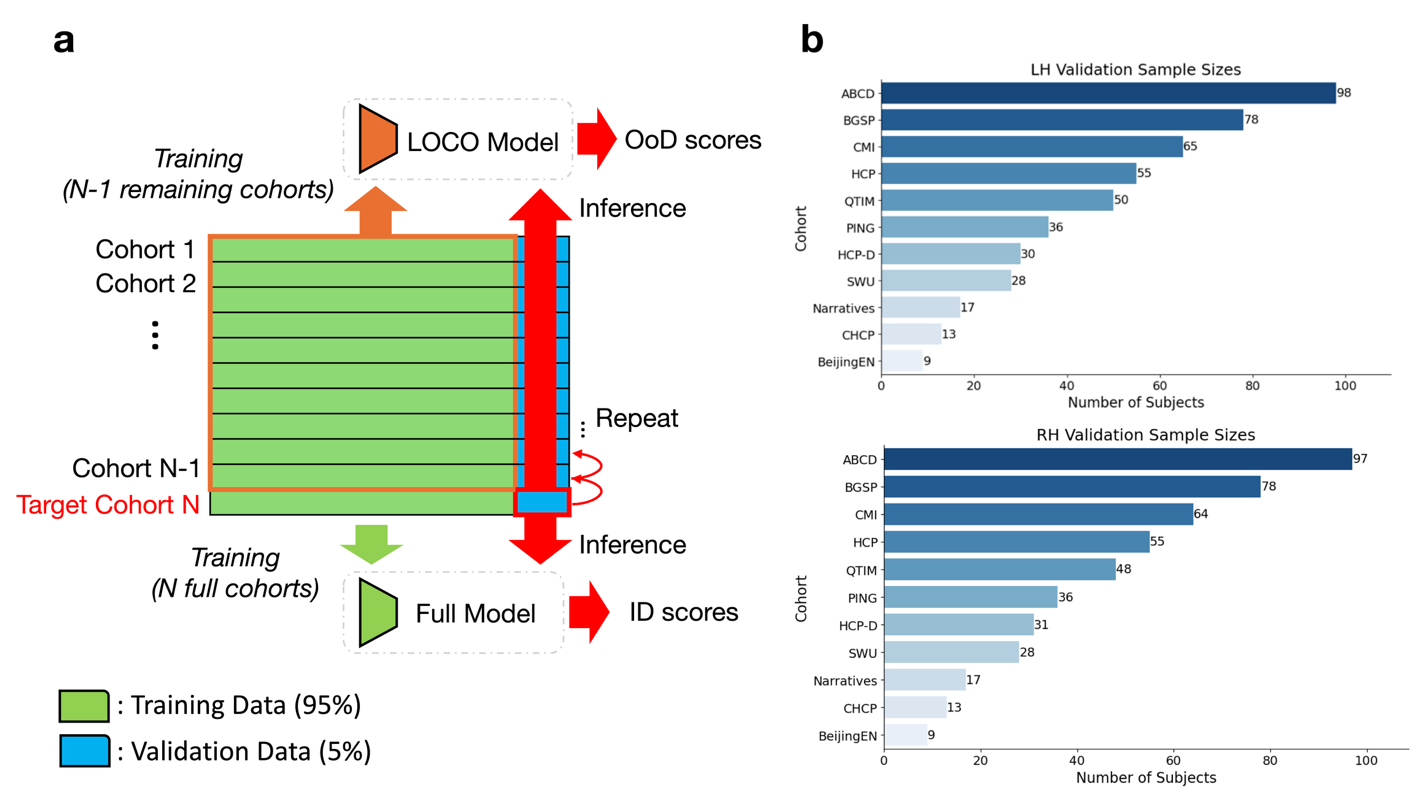

**Supplementary Fig. 1. LOCO validation.** (a) Schematic of the LOCO design. Each row represents one of the N=11 normative cohorts, partitioned into a 95% training set (green) and a 5% validation set (blue). For each target cohort (red outline), a LOCO model is trained on the training partitions of the ten remaining cohorts only (orange outline), while the Full model is the main model trained on the training partitions of all eleven cohorts. Both models are then applied to the same validation subjects of the target cohort, yielding paired OoD and ID deviation scores. Because the evaluated subjects are identical and only the training composition differs, the comparison isolates the cohort effect. (b) Number of validation subjects per cohort, for the left (top) and right (bottom) hemispheres, indicating the sample on which the paired scores are computed. The smaller cohorts yield correspondingly wider CIs in the agreement statistics (Fig. 5a-b).

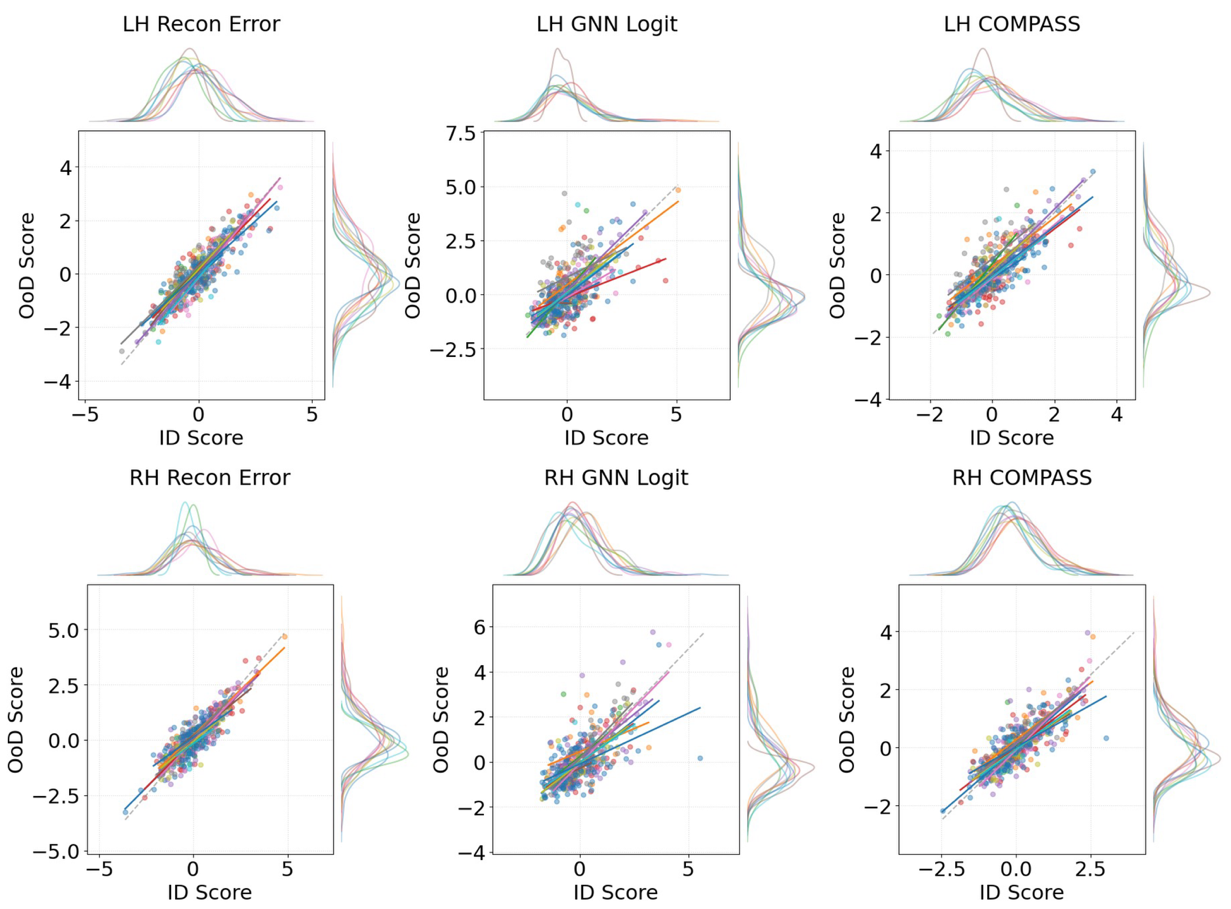

**Supplementary Fig. 2. Cohort-wise agreement between ID and OoD anomaly scores under LOCO validation.** Scatter plots of OoD versus ID anomaly scores for each subject, shown for the reconstruction error, GNN logit, and COMPASS (columns) in the left (top row) and right (bottom row) hemispheres under the LOCO design. OoD and ID are computed on the same subjects, so proximity to the diagonal (dashed line) reflects the stability of a score when its cohort is unseen during training. Points are colored by cohort, with per-cohort regression lines overlaid and marginal kernel-density estimates of the ID (top) and OoD (right) distributions shown along each axis. The reconstruction error and COMPASS cluster tightly along the diagonal with consistent per-cohort slopes, whereas the GNN logit shows greater scatter and cohort-dependent slope divergence, consistent with the ICCs reported in Fig. 5a-b.

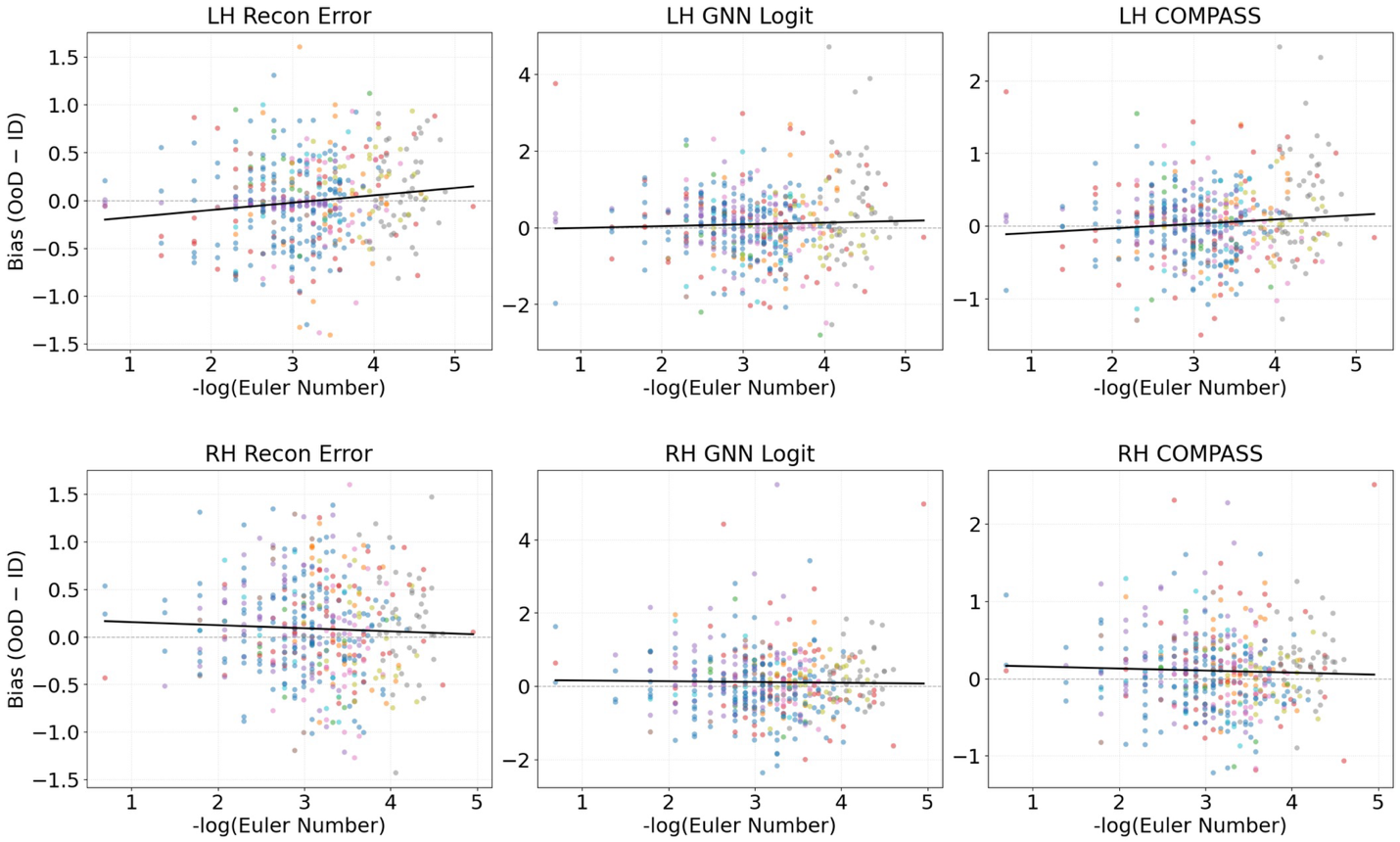

**Supplementary Fig. 3. Association between OoD bias and cortical surface quality.** Relationship between the LOCO bias and cortical surface quality for the reconstruction error, GNN logit, and COMPASS (columns) in the left (top row) and right (bottom row) hemispheres. For each subject, the y-axis is the bias (OoD - ID) between OoD and ID scores, and the x-axis is the surface-quality proxy (negative-log-transformed FreeSurfer Euler number, higher values denoting poorer quality). Each point is one subject, colored by cohort, with the fitted regression line overlaid. Corresponding LMM estimates are reported in Supplementary Table 19.

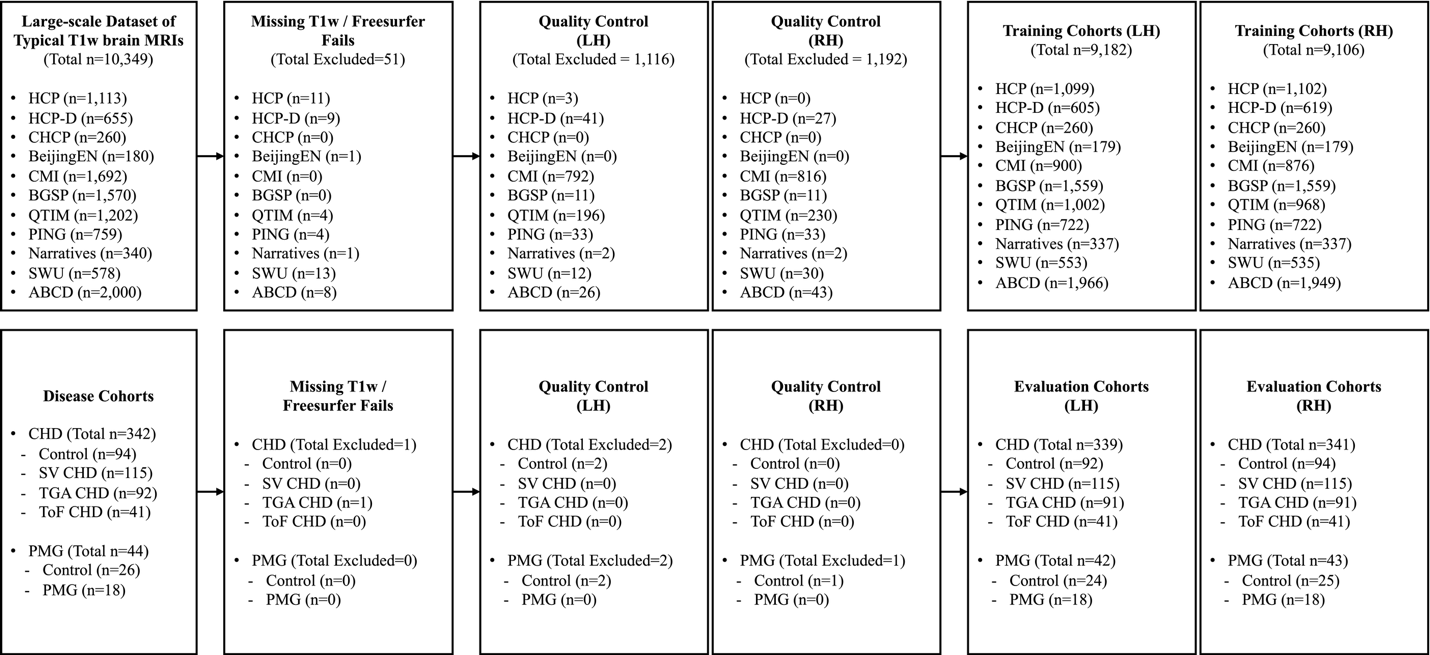

**Supplementary Fig. 4. Overview of the QC workflow for training and evaluation dataset.**

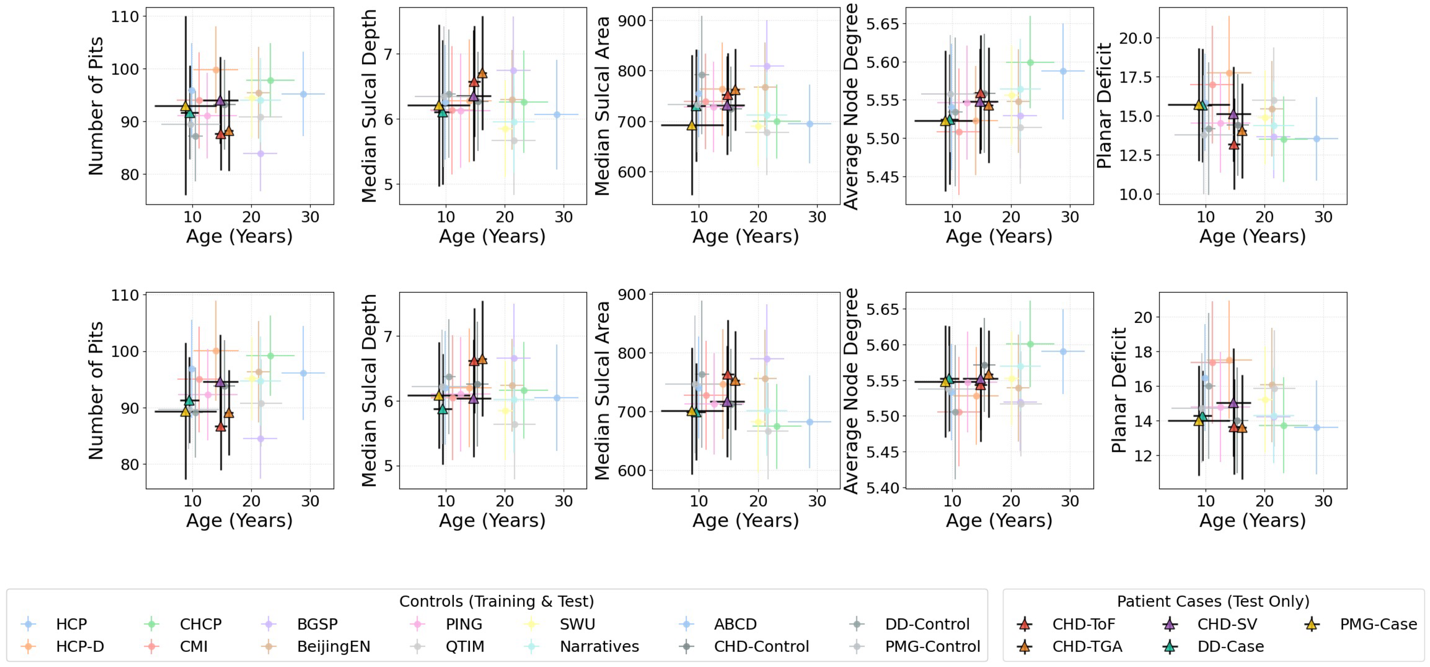

**Supplementary Fig. 5. Age-related distributions of sulcal graph properties across training and evaluation cohorts.** Distributions of five sulcal pattern graph properties as a function of age, shown for LH (top row) and RH (bottom row). Columns report, in order: the number of sulcal pits (graph nodes); the median sulcal depth and the median sulcal basin area (node attributes); and the average node degree and the average planar deficit (topological descriptors). Distributions are colored by cohort, spanning the eleven normative training cohorts (thin lines with circles) and the evaluation cohorts, with the latter separated into controls (thin lines with circles) and the SV, TGA, ToF, and PMG patient groups (thick lines with triangles). The planar deficit is defined as (3N − 6) − E for N nodes and E edges, with non-negative values and a mean degree below six indicating a valid genus-0 planar triangulation. Due to the profound inter-subject variability in cortical surfaces and sulcal folds, the number of sulcal pits and graph topology exhibit high variability.

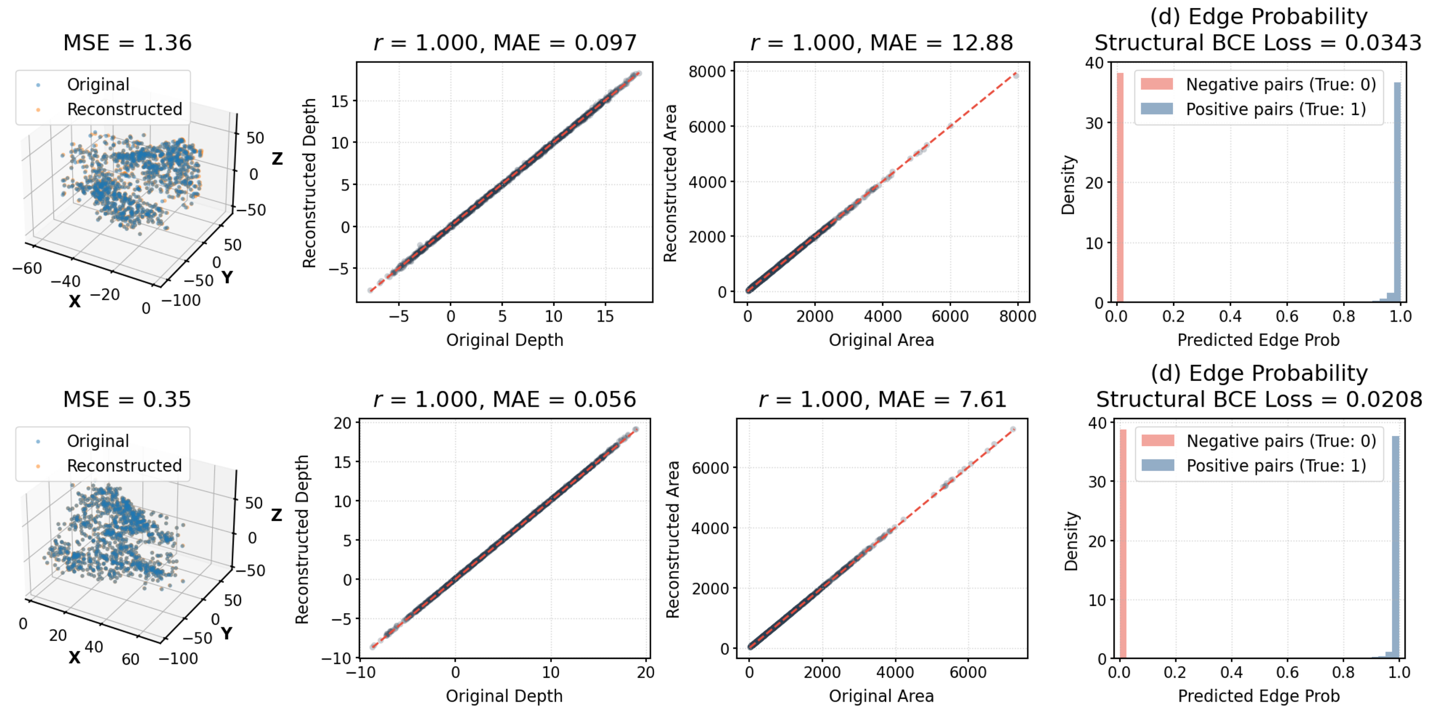

**Supplementary Fig. 6. Reconstruction fidelity of the HA-VGAE.** Node- and edge-level reconstruction quality of the HA-VGAE on normative training graphs, shown for the LH (top row) and RH (bottom row). First column: original versus reconstructed three-dimensional sulcal pit positions, plotted as nodes in space, summarized by the MSE between original and reconstructed coordinates. Second and third columns: original versus reconstructed node attributes, sulcal depth and sulcal basin area, respectively, with the Pearson correlation (r) and MAE between the two. Fourth column: distributions of the predicted edge probability for node pairs that are connected (positive pairs, true label 1) versus unconnected (negative pairs, true label 0) in the original graph, with the structural BCE loss reported; clear separation between the two distributions indicates faithful recovery of graph topology. Across both hemispheres, node positions, attributes, and edges were reconstructed with high fidelity (near-unity correlations, low MSE/MAE, and well-separated edge-probability distributions).

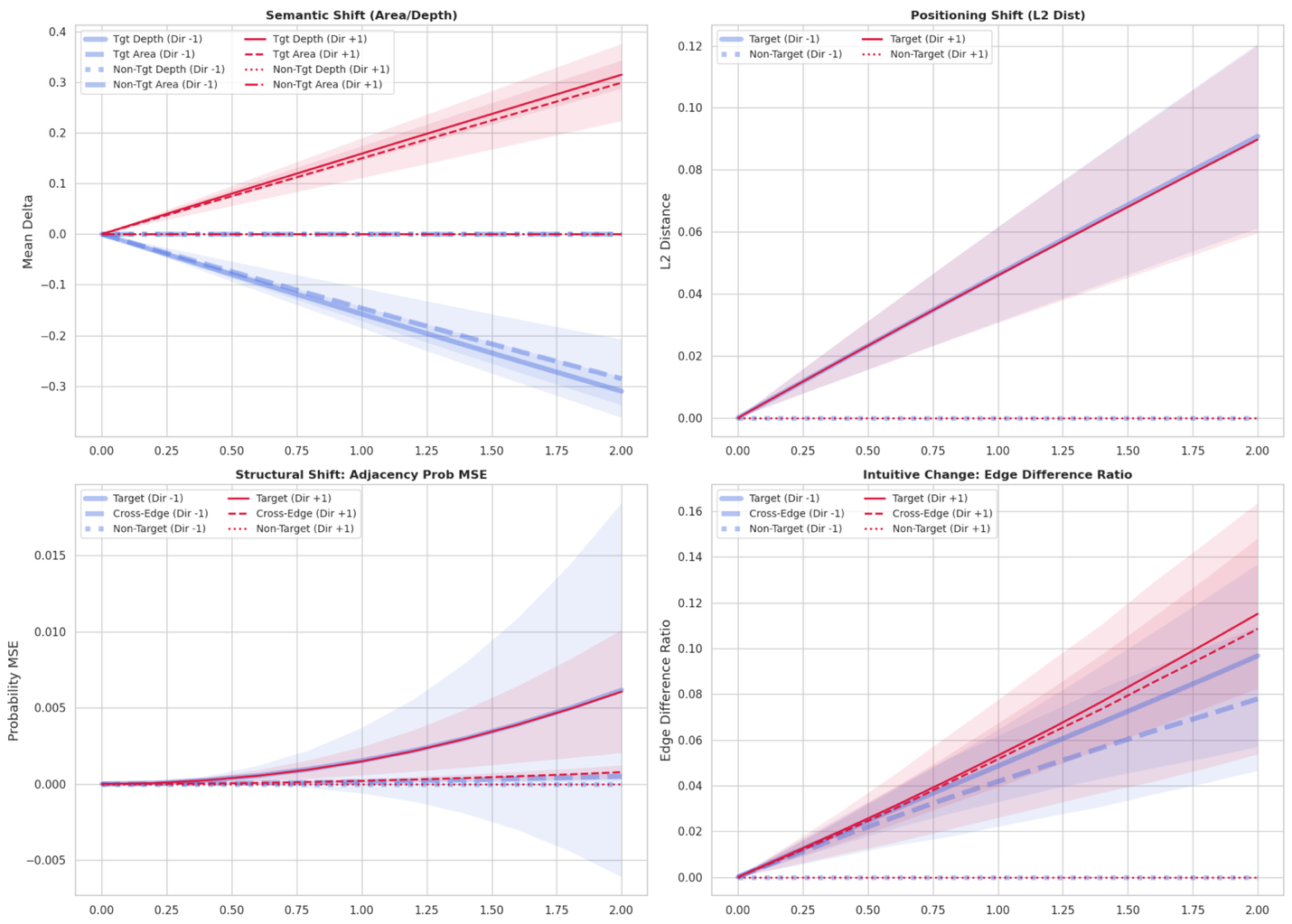

**Supplementary Fig. 7. Selectivity and directional control of the proposed LSS perturbation.** Effect of the LSS perturbation on sulcal graph properties as a function of the semantic shift magnitude $\kappa$ (x-axis) quantifying that the perturbation alters only its intended targets and in the intended direction, $\Gamma$ in Eq. 10. In all panels, color denotes the shift direction (blue, feature diminution, $\Gamma$ [Dir] -1; red, feature magnification, Dir +1), line style denotes the region relative to the stochastic mask $M$ (solid, target lobes; dashed, cross-boundary edges; dotted, non-target lobes), and shaded bands are SD across graphs. (Top left) mean change in the target morphological features (sulcal depth and area) within target versus non-target lobes. Target-region features shift monotonically and symmetrically with $\kappa$, increasing under Dir +1 and decreasing under Dir -1, whereas non-target features remain essentially unchanged, confirming that the shift is both directionally controlled and spatially confined. (Top right) L2 displacement of sulcal pit positions. Target-region positions shift progressively while non-target positions are unaffected, and the two directions coincide, indicating that positional change is driven by regional membership rather than shift direction. (Bottom left) MSE of the reconstructed adjacency probabilities. Edge structure changes gradually and predominantly within target regions, with smaller effects at cross-boundary edges and negligible change at non-target edges, showing that topological perturbation stays localized. (Bottom right) fraction of edges added or removed relative to the original graph. Edge turnover rises smoothly with $\kappa$, concentrated in target and cross-boundary regions, confirming that LSS induces spatially clustered structural change while leaving non-target topology intact.

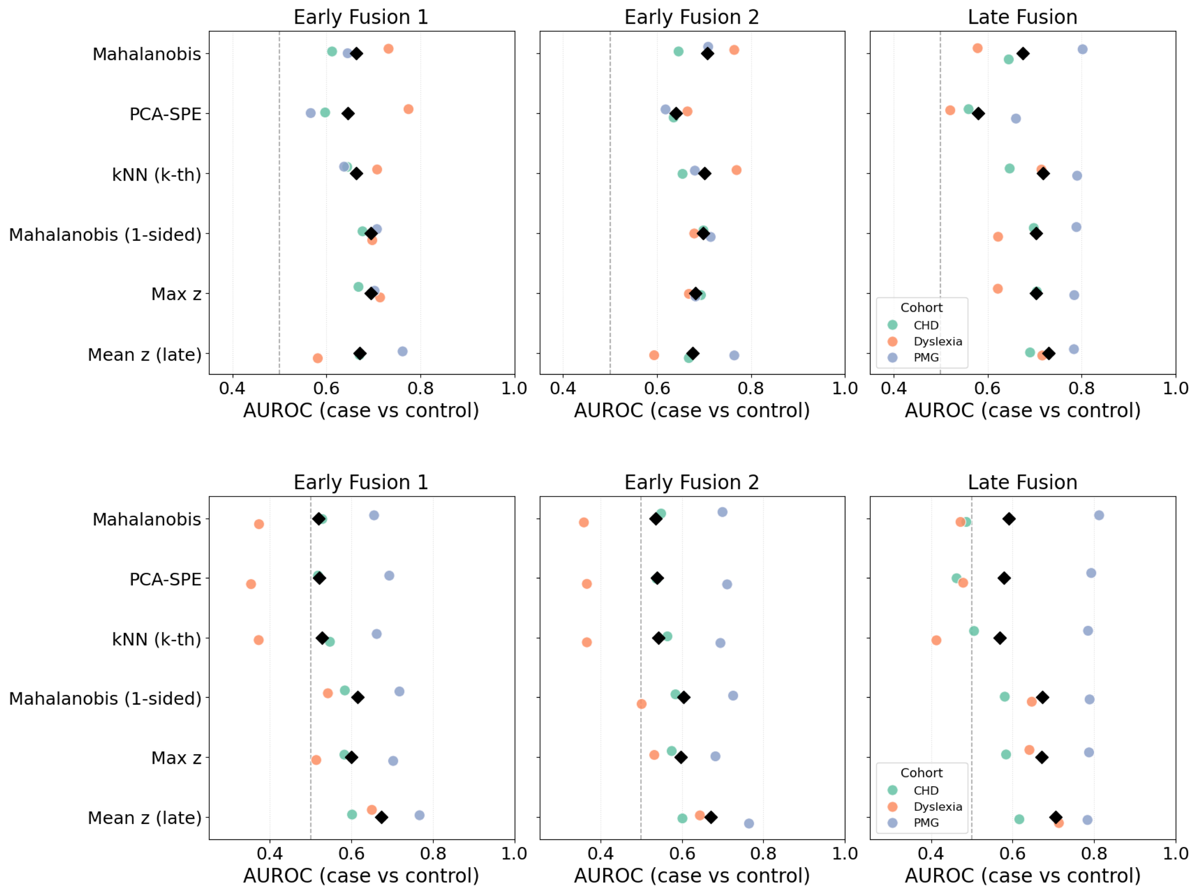

**Supplementary Fig. 8. Comparison of fusion levels and functions.** Case-versus-control AUROC for each combination of fusion level (columns) and function (rows) for LH (top) and RH (bottom). Fusion levels: [Early Fusion 1], the reconstruction error and the GNN logit are averaged within each lobe (4-D); [Early Fusion 2], both measures are retained for every lobe (8-D); [Late Fusion], each measure is first aggregated to the whole brain and the two resulting scalars are combined (2-D). Composite scoring functions map the resulting feature vector to a single anomaly score: *Mahalanobis*, distance from the normative mean weighted by the inverse covariance (Ledoit-Wolf shrinkage); *PCA-SPE*, squared residual outside the normative principal subspace (95% of variance retained); *kNN*, Euclidean distance to the 10th nearest normative neighbour; *one-sided Mahalanobis*, as Mahalanobis but with deviations below the normative mean set to zero; *max z*, the largest component; *mean z*, the unweighted mean of the components. All functions were fitted on the normative training set only and applied unchanged to the clinical cohorts, with no exposure to case labels; features were re-standardised using normative means and standard deviations. Each point is one clinical cohort (CHD, and PMG); black diamonds denote the across-cohort mean; the dashed line indicates chance (=0.5). AUROC uncertainty was estimated by 1,000 bootstrap resamples. The *late-fusion mean* achieved the highest across-cohort mean AUROC (0.73 for LH, 0.70 for RH) and is adopted as the composite score in the main analysis (COMPASS_1).

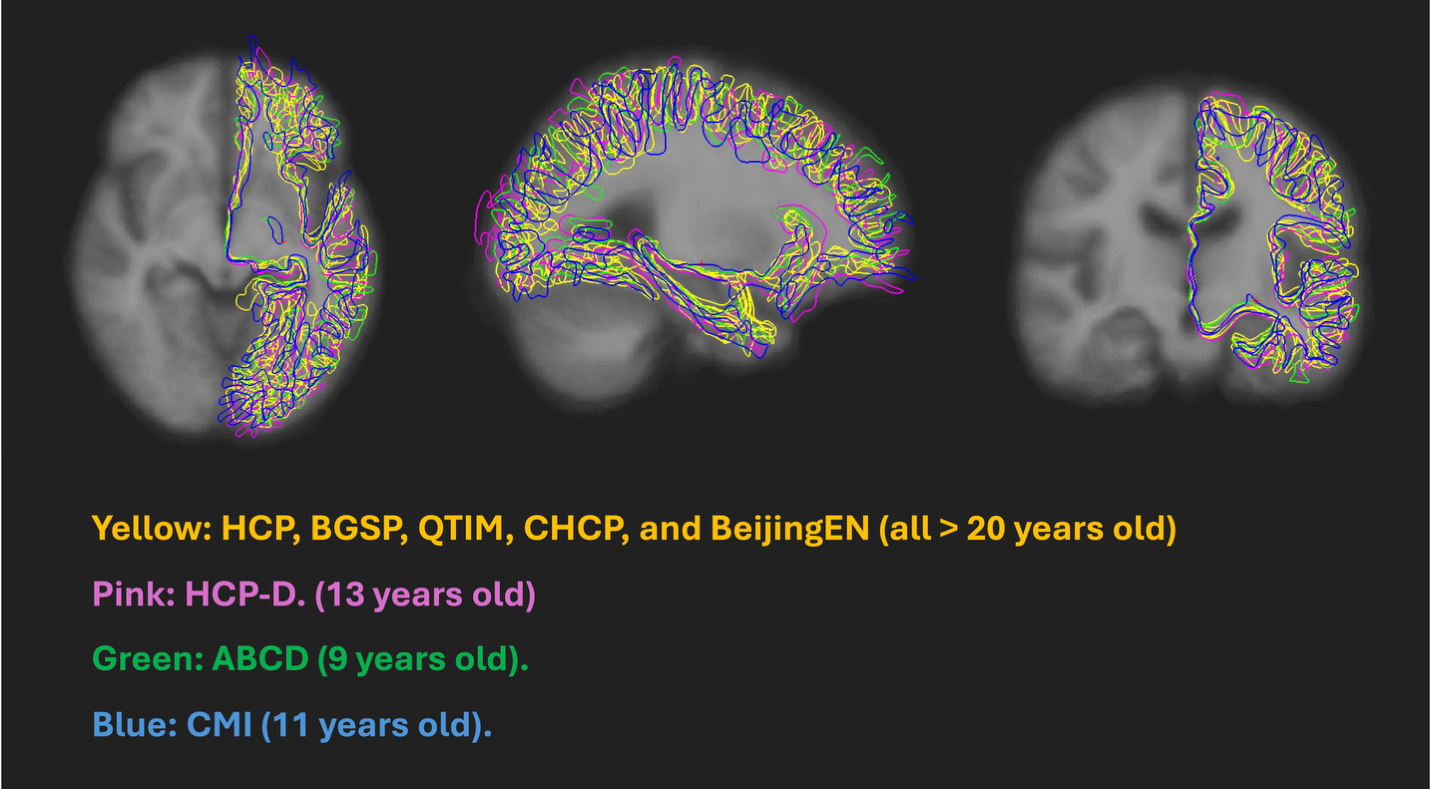

**Supplementary Fig. 9. Age-related residual misalignment of cortical surfaces after standard registration.** Cortical white matter surface contours from representative subjects of different cohorts, overlaid on a common brain template in axial, sagittal, and coronal views (left to right) after standard linear registration to MNI-305 space. Contours are colored by cohort and age: yellow denotes the examplar adult cohorts (HCP, BGSP, QTIM, CHCP, and BeijingEN, all > 20 years), pink HCP-D (13 years), green ABCD (9 years), and blue CMI (11 years). While the adult surfaces align closely, the youngest cohorts show small but systematic deviations from the adult contours, particularly at the cortical boundary, reflecting the greater difficulty of stereotaxic normalization in younger brains owing to lower tissue contrast and smaller brain volume. These residual, non-biological misalignments motivate the rigid-body coordinate augmentation applied during training (see Methods, "Implementation Details"), which perturbs sulcal pit positions to make the model robust to such site- and age-related spatial variation.

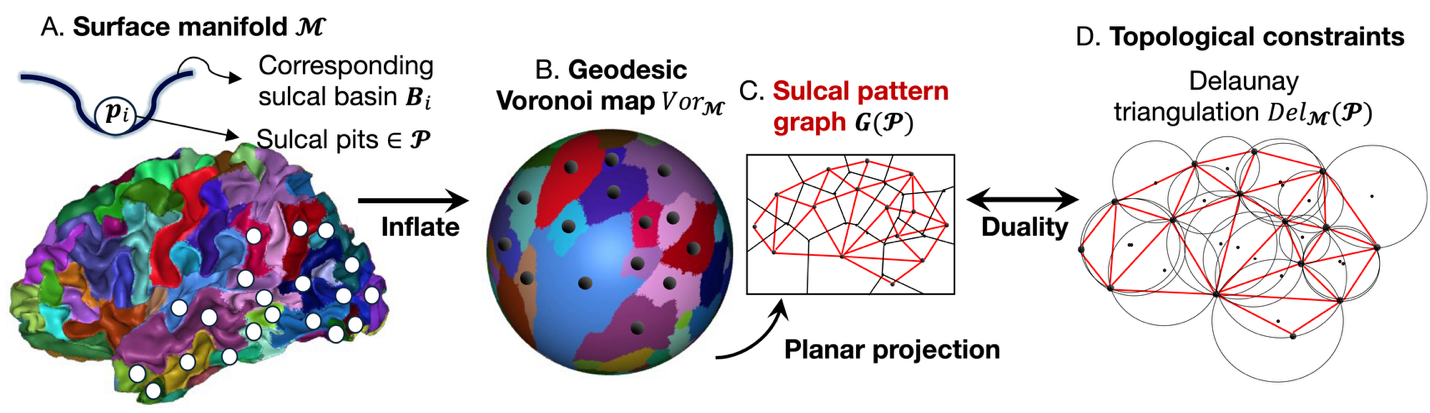

**Supplementary Fig. 10. Construction and topological duality of the sulcal pattern graph.** Schematic of how the sulcal pattern graph is defined and why it is constrained to a planar topology. (A) On the cortical surface manifold, local minima of the sulcal depth map are extracted as sulcal pits, each defining a surrounding sulcal basin. Inflating the surface to a sphere, the basins form a geodesic Voronoi map, partitioning the manifold into regions, one per pit. (C) The sulcal pattern graph is the adjacency graph of these basins, with nodes at pits and edges connecting basins that share a boundary; its planar projection is shown. (D) By the duality between Voronoi diagrams and Delaunay triangulations under the geodesic metric, sulcal pattern graph is topologically equivalent to the geodesic Delaunay triangulation, which on a genus-0 surface satisfies Euler's formula and the resulting constraints |E| ≤ 3|V| − 6 and average node degree below six. Full definitions and derivation are given in Supplementary Note 1.

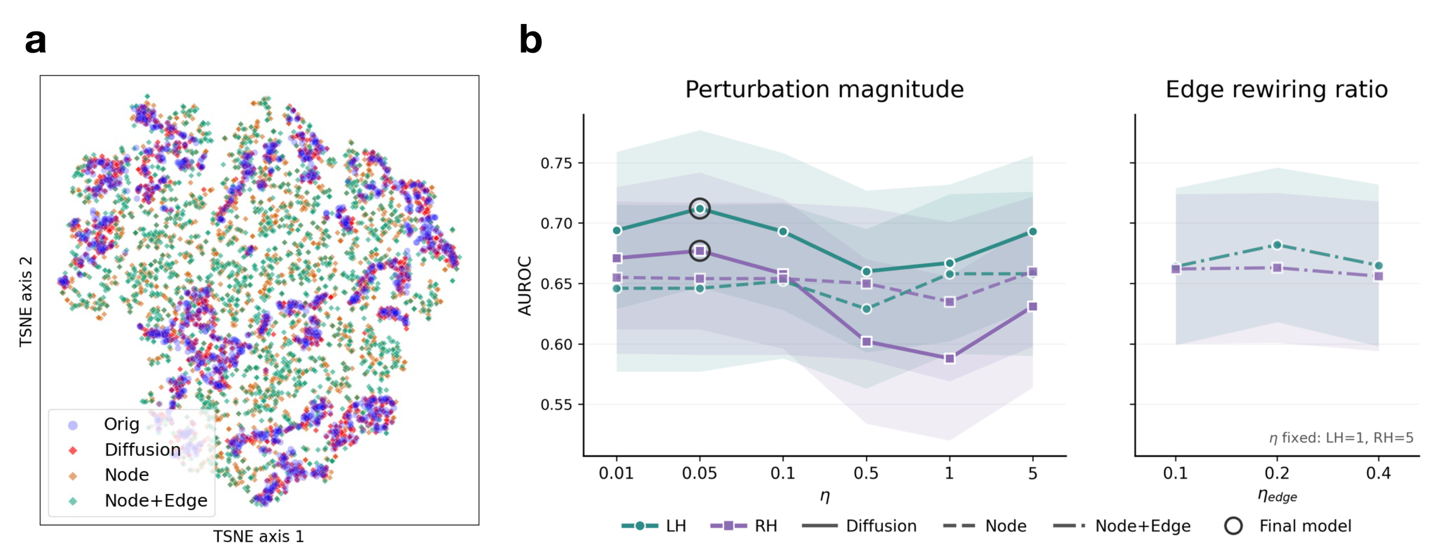

**Supplementary Fig. 11. Comparison of the proposed diffusion-based perturbation against explicit perturbation strategies.** (a) t-SNE projection of latent embeddings of masked nodes from validation normative graphs. Original node encodings (blue circles) are shown together with PA embeddings generated by the diffusion+LSS (red), **Node** (orange), and **Node+Edge** (green) strategies. For the two explicit strategies, corrupted graphs were re-encoded through the frozen HA-VGAE encoder. Diffusion-generated embeddings remain interleaved with the original distribution, whereas explicitly perturbed embeddings spread into regions where real data are sparse. (b) Mean AUROC across evaluation clinical cohorts and perturbation settings. (left) Effect of the perturbation magnitude $\eta$ (and $\eta_{node}$) for the diffusion (solid) and **Node** (dashed) strategies. (right) Effect of the edge rewiring ratio $\eta_{edge}$ for the **Node+Edge** strategy (dash-dotted), with $\eta$ fixed at the best value from the **Node** sweep in each hemisphere (LH $\eta_{node}=1$, RH $\eta_{node}=5$). Colours indicate hemisphere (teal, LH; purple, RH). Markers show the mean AUROC across five evaluation clinical cohorts and shaded bands the corresponding standard deviation. Circled points mark the setting used in the final model. All three strategies share the same lobar masking and random-walk smoothing, and differ only in how the PA graph is constructed.

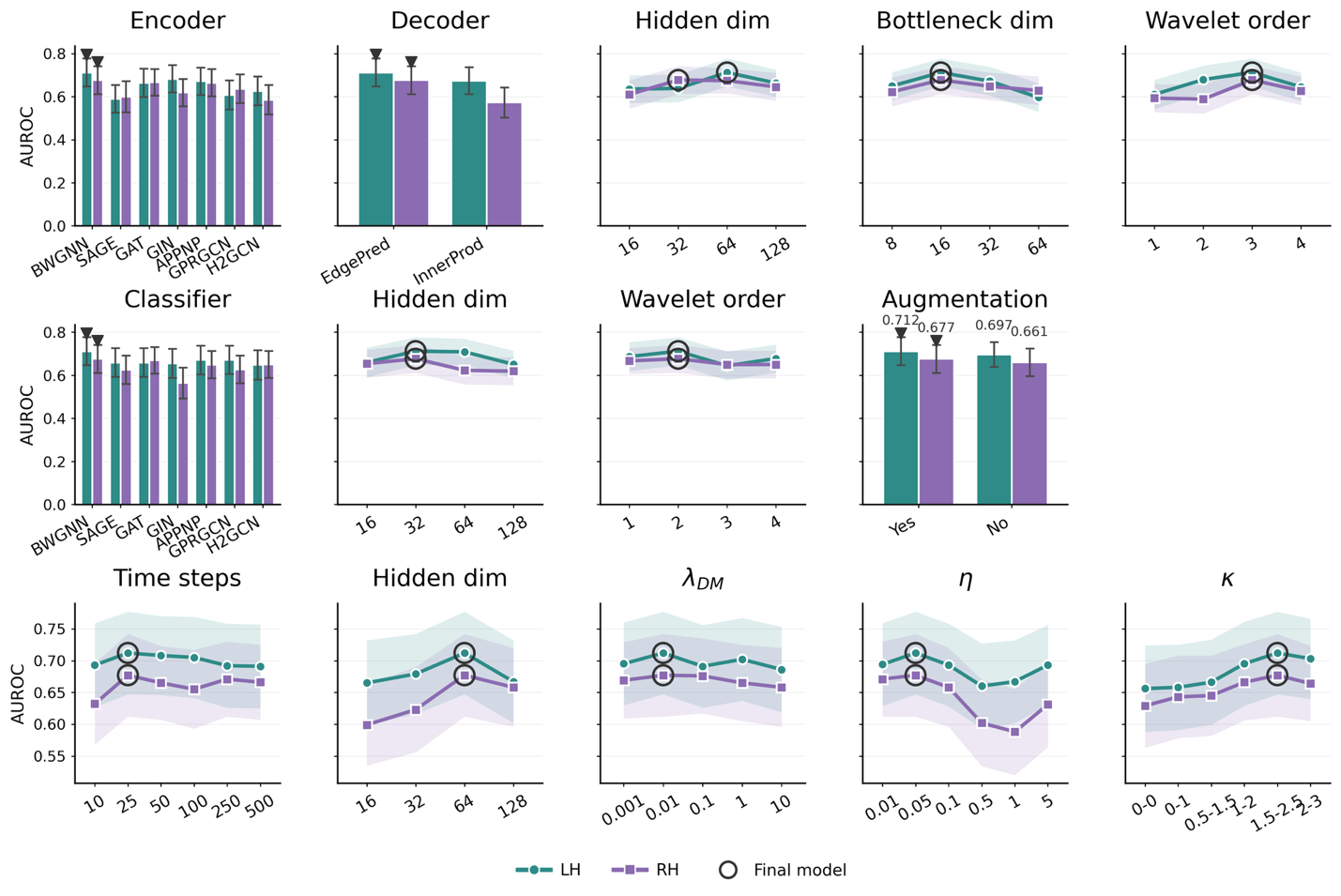

**Supplementary Fig. 12. Sensitivity of COMPASS to model configuration and hyperparameters.** Case-versus-control AUROC (mean ± SD over 1,000 bootstrap resamples) as each component of the framework is varied while all other settings are held at their final values. **(top row)** HA-VGAE pretraining: encoder architecture, decoder formulation, and the encoder's hidden dimension, bottleneck (latent) dimension, and Beta-wavelet order. **(middle row)** Anomaly classifier and input augmentation: classifier backbone, its hidden dimension and Beta-wavelet order, and the presence or absence of the proposed rigid-body rotation and translation of sulcal pit coordinates**. (bottom row)** Diffusion and perturbation: number of diffusion time steps, hidden dimension of the denoising network, diffusion-loss weight ($\lambda_{\text{DM}}$), and the two LSS parameters governing perturbation magnitude (η) and semantic-shift step size (κ). Green denotes LH and purple RH; bars show categorical comparisons with error bars, and lines show continuous hyperparameters with shaded ±SD bands. Markers indicate the configuration adopted in the final model. Across all factors, performance varied within a narrow band, with the spectral BWGNN encoder and classifier outperforming conventional alternatives and the rigid-body augmentation contributing a consistent gain.

**Supplementary Tables**

**Supplementary Table 1.** **Global anomaly-score group effects in CHD cohort.** Standardized effect sizes for the difference between each CHD subtype and healthy controls, estimated by GLM-based ANCOVA on global (whole-hemisphere) anomaly scores in both hemispheres, adjusting for age, sex, and socioeconomic status. Rows denote CHD subtypes; columns denote the six anomaly-scoring methods compared (SGM, OC-SVM, reconstruction error, GNN logit, COMPASS_1, and COMPASS_2). Each cell reports the standardized regression coefficient $\beta$, its 95% confidence interval [95% CI], and the FDR-corrected q-value. Positive $\beta$ indicates higher anomaly scores in patients than controls. FDR correction (Benjamini–Hochberg) was applied across each method; asterisks denote *q < 0.05, **q < 0.01, ***q < 0.001. SGM and COMPASS_2 carry an in-sample advantage and should be read as favorable-case references (see Methods, "Baseline methods" and "COMPASS"); COMPASS_1 is the primary, parameter-free metric.

| **CHD**  **Subtype** | **LH**  **SGM** | **LH**  **OCSVM** | **LH**  **Recon Error** | **LH**  **GNN Logit** | **LH COMPASS_1** | **LH COMPASS_2** |
| --- | --- | --- | --- | --- | --- | --- |
| SV | -0.005  [-0.007, -0.002] q<0.001*** | -0.028  [-0.282, 0.225] q=0.826 | 0.728  [0.428, 1.028] q<0.001*** | 0.387  [0.078, 0.696] q=0.014* | 0.558  [0.314, 0.801] q<0.001*** | 0.639  [0.383, 0.896] q<0.001*** |
| TGA | -0.002  [-0.004, 0.001] q=0.155 | -0.102  [-0.377, 0.173] q=0.826 | 0.595  [0.269, 0.920] q<0.001*** | 0.669  [0.333, 1.004] q<0.001*** | 0.632  [0.368, 0.895] q<0.001*** | 0.614  [0.336, 0.892] q<0.001*** |
| ToF | -0.004  [-0.007, -0.001] q=0.016* | -0.096  [-0.432, 0.240] q=0.826 | 0.414  [0.017, 0.812] q=0.041* | 0.754  [0.345, 1.163] q<0.001*** | 0.584  [0.262, 0.906] q<0.001*** | 0.503  [0.163, 0.842] q=0.004** |
| **CHD**  **Subtype** | **RH**  **SGM** | **RH**  **OCSVM** | **RH**  **Recon Error** | **RH**  **GNN Logit** | **RH COMPASS_1** | **RH COMPASS_2** |
| SV | -0.004  [-0.007, -0.002] q<0.001*** | -0.107  [-0.405, 0.191] q=0.480 | 0.348  [0.079, 0.618] q=0.017* | 0.240  [-0.002, 0.482] q=0.052 | 0.294  [0.085, 0.503] q=0.006** | 0.290  [0.082, 0.498] q=0.006** |
| TGA | -0.002  [-0.004, 0.001] q=0.206 | -0.186  [-0.509, 0.138] q=0.390 | 0.565  [0.272, 0.858] q<0.001*** | 0.424  [0.161, 0.687] q=0.003** | 0.494  [0.268, 0.721] q<0.001*** | 0.489  [0.263, 0.714] q<0.001*** |
| ToF | -0.004  [-0.007, -0.001] q=0.025* | -0.281  [-0.676, 0.114] q=0.390 | 0.416  [0.059, 0.774] q=0.023* | 0.678  [0.357, 0.999] q<0.001*** | 0.547  [0.270, 0.824] q<0.001*** | 0.557  [0.282, 0.833] q<0.001*** |

**Supplementary Table 2.** **Regional anomaly-score group effects in CHD cohort.** Standardized effect sizes for the difference between each CHD subtype and healthy controls, estimated by a MMRM with an unstructured covariance matrix on regional (lobar) anomaly scores in both hemispheres, treating the four lobar regions of each subject as repeated measures and adjusting for age, sex, and socioeconomic status. A lobe-by-subtype interaction was tested first and, being non-significant as mentioned in main body, removed; the reported coefficients therefore reflect the pooled regional group effect across lobes. Rows denote CHD subtypes; columns denote the six anomaly-scoring methods compared. Each cell reports the standardized coefficient $\beta$, its 95% confidence interval [95% CI], and the FDR-corrected q-value. Positive $\beta$ indicates higher anomaly scores in patients than controls. FDR correction (Benjamini–Hochberg) was applied across each method; asterisks denote *q < 0.05, **q < 0.01, ***q < 0.001. SGM and COMPASS_2 carry an in-sample advantage and should be read as favorable-case references (see Methods, "Baseline methods" and "COMPASS"); COMPASS_1 is the primary, parameter-free metric.

| **CHD**  **Subtype** | **LH**  **SGM** | **LH**  **OCSVM** | **LH**  **Recon Error** | **LH**  **GNN Logit** | **LH COMPASS_1** | **LH COMPASS_2** |
| --- | --- | --- | --- | --- | --- | --- |
| SV | -0.003  [-0.005, -0.001] q=0.013* | -0.109  [-0.271, 0.053] q=0.187 | 0.471  [0.279, 0.663] q<0.001*** | 0.090  [-0.065, 0.246] q=0.254 | 0.297  [0.156, 0.438] q<0.001*** | 0.386  [0.227, 0.546] q<0.001*** |
| TGA | -0.002  [-0.005, 0.000] q=0.104 | -0.288  [-0.470, -0.105] q=0.006** | 0.355  [0.139, 0.571] q=0.002** | 0.311  [0.136, 0.486] q=0.001** | 0.348  [0.189, 0.507] q<0.001*** | 0.352  [0.173, 0.532] q<0.001*** |
| ToF | -0.004  [-0.007, -0.001] q=0.014* | -0.170  [-0.387, 0.047] q=0.186 | 0.210  [-0.047, 0.466] q=0.109 | 0.180  [-0.027, 0.388] q=0.133 | 0.227  [0.038, 0.416] q=0.019* | 0.231  [0.018, 0.445] q=0.034* |
| **CHD**  **Subtype** | **RH**  **SGM** | **RH**  **OCSVM** | **RH**  **Recon Error** | **RH**  **GNN Logit** | **RH COMPASS_1** | **RH COMPASS_2** |
| SV | -0.003  [-0.005, -0.001] q=0.024* | -0.040  [-0.187, 0.107] q=0.660 | 0.177  [0.008, 0.346] q=0.060 | 0.190  [0.048, 0.331] q=0.009** | 0.195  [0.068, 0.322] q=0.004** | 0.195  [0.068, 0.321] q=0.004** |
| TGA | -0.002  [-0.004, 0.001] q=0.154 | -0.037  [-0.203, 0.128] q=0.660 | 0.257  [0.067, 0.448] q=0.025* | 0.251  [0.091, 0.411] q=0.003** | 0.249  [0.106, 0.392] q=0.002** | 0.248  [0.106, 0.391] q=0.002** |
| ToF | -0.004  [-0.007, -0.001] q=0.024* | -0.142  [-0.339, 0.054] q=0.468 | 0.206  [-0.020, 0.432] q=0.075 | 0.306  [0.116, 0.496]  q=0.003** | 0.245  [0.074, 0.415] q=0.005** | 0.248  [0.079, 0.418] q=0.004** |

**Supplementary Table 3.** **Global anomaly-score group effects after adjusting for surface quality proxy (Euler number) in CHD cohort.** In addition to the basleine covariates (age, sex and socioeconomic status), the Euler number was added to control the effect of surface quality, indexed by Freesurfer-induced Euler number. The Euler number has been used as an index of the topological defects of raw cortical surfaces. As shown in this results, there is no significant impact on the group effect identified in Supplementary Table 1. Each cell reports the standardized coefficient $\beta$, its 95% confidence interval [95% CI], and the FDR-corrected q-value. Positive $\beta$ indicates higher anomaly scores in patients than controls. FDR correction (Benjamini–Hochberg) was applied across each method; asterisks denote *q < 0.05, **q < 0.01, ***q < 0.001.

| **CHD**  **Subtype** | **LH**  **SGM** | **LH**  **OCSVM** | **LH**  **Recon Error** | **LH**  **GNN Logit** | **LH COMPASS_1** | **LH COMPASS_2** |
| --- | --- | --- | --- | --- | --- | --- |
| SV | -0.003  [-0.006, -0.001] q=0.009** | 0.005  [-0.247, 0.257] q=0.971 | 0.684  [0.387, 0.981] q<0.001*** | 0.357  [0.049, 0.666] q=0.023* | 0.521  [0.281, 0.761] q<0.001*** | 0.586  [0.338, 0.835] q<0.001*** |
| TGA | -0.002  [-0.004, 0.001] q=0.2315 | -0.067  [-0.340, 0.206] q=0.947 | 0.548  [0.226, 0.870] q=0.001** | 0.637  [0.302, 0.971] q<0.001*** | 0.592  [0.332, 0.853] q<0.001*** | 0.575  [0.305, 0.844] q<0.001*** |
| ToF | -0.002  [-0.005, 0.001] q=0.2315 | -0.132  [-0.465, 0.201] q=0.947 | 0.461  [0.068, 0.854] q=0.022* | 0.786  [0.379, 1.194] q<0.001*** | 0.624  [0.306, 0.941] q<0.001*** | 0.559  [0.230, 0.887] q<0.001*** |
| **CHD**  **Subtype** | **RH**  **SGM** | **RH**  **OCSVM** | **RH**  **Recon Error** | **RH**  **GNN Logit** | **RH COMPASS_1** | **RH COMPASS_2** |
| SV | -0.004  [-0.006, -0.001] q=0.006** | -0.107  [-0.405, 0.191] q=0.481 | 0.341  [0.073, 0.609] q=0.0172* | 0.237  [-0.005, 0.478] q=0.055 | 0.289  [0.082, 0.496] q=0.006** | 0.285  [0.078, 0.491] q=0.007** |
| TGA | -0.001  [-0.004, 0.001] q=0.234 | -0.185  [-0.509, 0.140] q=0.396 | 0.542  [0.250, 0.833] q<0.001*** | 0.411  [0.148, 0.674] q=0.004** | 0.476  [0.251, 0.702] q<0.001*** | 0.471  [0.247, 0.696] q<0.001*** |
| ToF | -0.002  [-0.005, 0.001] q=0.234 | -0.282  [-0.677, 0.114] q=0.396 | 0.433  [0.077, 0.788] (q=0.0172) | 0.688  [0.367, 1.009] q<0.001*** | 0.560  [0.285, 0.835] q<0.001*** | 0.570  [0.297, 0.844] q<0.001*** |

**Supplementary Table 4.** **Regional anomaly-score group effects after adjusting for surface quality proxy (Euler number) in CHD cohort.** In addition to the basleine covariates (age, sex and socioeconomic status), the Euler number was added to control the effect of surface quality, indexed by Freesurfer-induced Euler number. The Euler number has been used as an index of the topological defects of raw cortical surfaces. As shown in this results, there is no significant impact on the group effect identified in Supplementary Table 2. Each cell reports the standardized coefficient $\beta$, its 95% confidence interval [95% CI], and the FDR-corrected q-value. Positive $\beta$ indicates higher anomaly scores in patients than controls. FDR correction (Benjamini–Hochberg) was applied across each method; asterisks denote *q < 0.05, **q < 0.01, ***q < 0.001.

| **CHD**  **Subtype** | **LH**  **SGM** | **LH**  **OCSVM** | **LH**  **Recon Error** | **LH**  **GNN Logit** | **LH COMPASS_1** | **LH COMPASS_2** |
| --- | --- | --- | --- | --- | --- | --- |
| SV | -0.003  [-0.005, -0.001] q=0.013* | -0.087  [-0.248, 0.074] q=0.291 | 0.451  [0.261, 0.641] q<0.001*** | 0.082  [-0.074, 0.237] q=0.304 | 0.280  [0.141, 0.420] q<0.001*** | 0.355  [0.201, 0.508] q<0.001*** |
| TGA | -0.002  [-0.004, 0.001] q=0.148 | -0.268  [-0.448, -0.087] q=0.011* | 0.332  [0.119, 0.545] q=0.003** | 0.303  [0.128, 0.478] q=0.002** | 0.333  [0.176, 0.489] q<0.001*** | 0.333  [0.161, 0.506] q<0.001*** |
| ToF | -0.004  [-0.007, -0.001] q=0.010* | -0.201  [-0.416, 0.015]  q=0.102 | 0.258  [0.004, 0.512] q=0.047 | 0.203  [-0.006, 0.411] q=0.085 | 0.259  [0.073, 0.446] q=0.007** | 0.272  [0.066, 0.477] q=0.010* |
| **CHD**  **Subtype** | **RH**  **SGM** | **RH**  **OCSVM** | **RH**  **Recon Error** | **RH**  **GNN Logit** | **RH COMPASS_1** | **RH COMPASS_2** |
| SV | -0.003  [-0.005, -0.001] q=0.017* | -0.037  [-0.183, 0.109] q=0.731 | 0.176  [0.008, 0.344] q=0.056 | 0.190  [0.048, 0.331] q=0.009** | 0.196  [0.069, 0.323] q=0.003** | 0.196  [0.070, 0.321] q=0.003** |
| TGA | -0.002  [-0.004, 0.001] q=0.184 | -0.029  [-0.194, 0.136] q=0.731 | 0.249  [0.059, 0.438] q=0.030* | 0.248  [0.088, 0.407] q=0.004** | 0.243  [0.100, 0.386] q=0.003** | 0.242  [0.100, 0.384] q=0.003** |
| ToF | -0.004  [-0.007, -0.001] q=0.017* | -0.152  [-0.348, 0.044] q=0.385 | 0.220  [-0.006, 0.445] q=0.056 | 0.313  [0.122, 0.503] q=0.004** | 0.257  [0.087, 0.427] q=0.003** | 0.260  [0.092, 0.429] q=0.003** |

**Supplementary Table 5. Regional post-hoc group differences across neurodevelopmental cohorts.** Post-hoc comparisons of regional COMPASS_1 anomaly scores between each patient group and its controls, computed per lobe by independent two-sample t-tests. Rows denote the five diseases; columns denote the eight cortical regions. Each cell reports the mean difference (patients - controls), its 95% CI, and the FDR-corrected q-value. Positive values indicate higher anomaly scores in patients. FDR correction (Benjamini–Hochberg) was applied across the four lobes within each cohort and hemisphere; asterisks denote *q < 0.05, **q < 0.01, ***q < 0.001.

| **Disease Subtype** | **Left Frontal** | **Left Temporal** | **Left Parietal** | **Left Occipital** |
| --- | --- | --- | --- | --- |
| SV | 0.495 [0.282, 0.707] q<0.001*** | 0.604 [0.401, 0.807] q<0.001*** | 0.254 [0.036, 0.472] q=0.030* | 0.188 [-0.032, 0.409] q=0.094 |
| TGA | 0.344 [0.123, 0.565] q=0.005** | 0.475 [0.254, 0.695] q<0.001*** | 0.208 [-0.036, 0.451] q=0.126 | 0.180 [-0.069, 0.429] q=0.156 |
| ToF | 0.142 [-0.200, 0.484] q=0.409 | 0.462 [0.113, 0.810] q=0.041* | 0.212 [-0.058, 0.481] q=0.244 | 0.169 [-0.114, 0.452] q=0.318 |
| PMG | 1.528 [0.192, 2.864] q=0.039 | -0.313 [-1.029, 0.403] q=0.381 | 1.181 [0.131, 2.231] q=0.039* | 0.902 [0.099, 1.706] q=0.039* |
| **Disease Subtype** | **Right Frontal** | **Right Temporal** | **Right Parietal** | **Right Occipital** |
| SV | 0.429 [0.214, 0.643] q<0.001*** | 0.143 [-0.050, 0.336] q=0.292 | 0.126 [-0.076, 0.328] q=0.293 | 0.008 [-0.217, 0.234] q=0.943 |
| TGA | 0.222 [0.012, 0.431] q=0.153 | 0.132 [-0.078, 0.341] q=0.358 | 0.122 [-0.095, 0.340] q=0.358 | 0.083 [-0.145, 0.311] q=0.472 |
| ToF | 0.333 [0.114, 0.551] q=0.013* | 0.327 [0.044, 0.609] q=0.048* | 0.115 [-0.152, 0.383] q=0.525 | 0.063 [-0.218, 0.345] q=0.656 |
| PMG | 0.384 [-0.132, 0.900] q=0.187 | 0.754 [-0.011, 1.520] q=0.106 | 0.361 [-0.369, 1.092] q=0.320 | 0.928 [0.273, 1.583] q=0.027* |

**Supplementary Table 6. Global anomaly-score group differences in the PMG cohort.** Group differences in global (whole-hemisphere) anomaly scores between PMG patients (n = 18) and typical controls (n = 26), reported for each hemisphere by two complementary tests: an independent two-sample t-test (parametric) and a Mann–Whitney U test (non-parametric), the former used as the primary test. Rows are grouped by hemisphere (left, right), with the t-test and U-test results on separate rows within each; columns denote the six anomaly-scoring methods compared. For the t-test rows, each cell reports the mean difference (patients - controls), its 95% CI, and the p-value; for the U-test rows, each cell reports the U statistic and its p-value. Asterisks denote *p < 0.05, **p < 0.01, and ***p < 0.001.

| **Test Type** | **LH**  **SGM** | **LH**  **OCSVM** | **LH**  **Recon Error** | **LH**  **GNN Logit** | **LH COMPASS_1** | **LH COMPASS_2** |
| --- | --- | --- | --- | --- | --- | --- |
| U test | U=42  p<0.001*** | U=71  p=0.001** | U=276 p=0.015* | U=281 p=0.010* | U=297 p=0.002** | U=301 p=0.002** |
| T-test | -0.018  [-0.024, -0.011] p<0.001*** | -2.033  [-3.230, -0.835] p=0.002** | 1.622  [0.074, 3.170] p=0.041* | 1.841  [0.384, 3.299] p=0.016* | 1.732  [0.377, 3.086] p=0.015* | 1.765  [0.411, 3.118] p=0.013* |
| **Test Type** | **RH**  **SGM** | **RH**  **OCSVM** | **RH**  **Recon Error** | **RH**  **GNN Logit** | **RH COMPASS_1** | **RH COMPASS_2** |
| U test | U=44 p<0.001*** | U=145  p=0.153 | U=303 p=0.004** | U=278 p=0.031* | U=299 p=0.006** | U=309 p=0.003** |
| T-test | -0.018  [-0.024, -0.011] p<0.001*** | -0.848  [-1.851, 0.155] p=0.094 | 0.977  [0.242, 1.712] p=0.011* | 0.711  [-0.273, 1.696] p=0.152 | 0.844  [0.166, 1.522] p=0.016* | 0.937  [0.263, 1.611] p=0.008** |

**Supplementary Table 7. Associations between global LH anomaly scores and ND outcomes in CHD.** Associations between each LH anomaly score and ND outcomes in the CHD cohort, estimated by GLMs adjusting for age, sex, socioeconomic status, and cognitive test type (WISC or WAIS, according to age). Rows denote the ND outcomes assessed; columns denote the six anomaly-scoring methods compared. Each cell reports the standardized regression coefficient $\beta$, its 95% CI, and the FDR-corrected q-value. A negative $\beta$ indicates that higher anomaly scores are associated with poorer outcome. FDR correction (Benjamini–Hochberg) was applied across the eight outcomes within each method; asterisks denote *q < 0.05.

| **ND Outcome** | **SGM** | **OCSVM** | **Recon Error** | **GNN Logit** | **COMPASS_1** | **COMPASS_2** |
| --- | --- | --- | --- | --- | --- | --- |
| Executive Function | 23.880  [-2.416, 50.176] (q=0.150) | -0.015  [-0.256, 0.225] (q=0.926) | -0.085  [-0.285, 0.116] (q=0.543) | 0.001  [-0.200, 0.202] (q=0.993) | -0.065  [-0.316, 0.185] (q=0.811) | -0.086  [-0.322, 0.150] (q=0.630) |
| Full-scale IQ | 166.658  [-24.922, 358.238] (q=0.150) | 0.966  [-0.791, 2.722] (q=0.769) | -1.183  [-2.646, 0.280] (q=0.300) | -0.472  [-1.906, 0.963] (q=0.993) | -1.285  [-3.098, 0.528] (q=0.328) | -1.382  [-3.098, 0.335] (q=0.227) |
| Verbal Comprehension | 161.274  [-27.451, 350.000] (q=0.150) | 0.492  [-1.241, 2.224] (q=0.769) | -0.125  [-1.572, 1.321] (q=0.865) | 0.058  [-1.356, 1.472] (q=0.993) | -0.050  [-1.841, 1.741] (q=0.956) | -0.106  [-1.803, 1.591] (q=0.902) |
| Perceptual Reasoning | 72.770  [-114.416, 259.957] (q=0.445) | 0.851  [-0.844, 2.545] (q=0.769) | -0.919  [-2.346, 0.508] (q=0.330) | -0.840  [-2.231, 0.551] (q=0.658) | -1.378  [-3.140, 0.384] (q=0.328) | -1.251  [-2.923, 0.421] (q=0.227) |
| General Memory | 175.111  [-43.310, 393.531] (q=0.154) | 0.576  [-1.413, 2.566] (q=0.769) | -1.980  [-3.634, -0.325] (q=0.077) | 0.239  [-1.383, 1.861] (q=0.993) | -1.319  [-3.377, 0.738] (q=0.333) | -1.925  [-3.871, 0.021] (q=0.210) |
| Processing Speed | 181.209  [-19.626, 382.045] (q=0.150) | 1.369  [-0.453, 3.191] (q=0.769) | -2.186  [-3.709, -0.664] (q=0.040*) | -1.083  [-2.580, 0.414] (q=0.658) | -2.541  [-4.426, -0.657] (q=0.067) | -2.632  [-4.415, -0.848] (q=0.032*) |
| Reading Composite Score | 110.187  [-77.051, 297.425] (q=0.283) | -0.080  [-1.773, 1.614] (q=0.926) | -0.484  [-1.907, 0.938] (q=0.575) | 0.063  [-1.362, 1.488] (q=0.993) | -0.329  [-2.106, 1.448] (q=0.818) | -0.473  [-2.144, 1.199] (q=0.661) |
| Math Composite Score | 273.669  [24.181, 523.157] (q=0.150) | 0.982  [-1.283, 3.247] (q=0.769) | -1.354  [-3.254, 0.546] (q=0.324) | -1.124  [-3.028, 0.781] (q=0.658) | -1.930  [-4.301, 0.441] (q=0.328) | -1.786  [-4.017, 0.446] (q=0.227) |

**Supplementary Table 8. Associations between global RH anomaly scores and ND outcomes in CHD.** Associations between each RH anomaly score and ND outcomes in the CHD cohort, estimated by GLMs adjusting for age, sex, socioeconomic status, and cognitive test type (WISC or WAIS, according to age). Rows denote the ND outcomes assessed; columns denote the six anomaly-scoring methods compared. Each cell reports the standardized regression coefficient $\beta$, its 95% CI, and the FDR-corrected q-value. A negative $\beta$ indicates that higher anomaly scores are associated with poorer outcome. FDR correction (Benjamini-Hochberg) was applied across the eight outcomes within each method.

| **ND Outcome** | **SGM** | **OCSVM** | **Recon Error** | **GNN Logit** | **COMPASS_1** | **COMPASS_2** |
| --- | --- | --- | --- | --- | --- | --- |
| Executive Function | 23.881  [-2.304, 50.066] (q=0.163) | 0.172  [-0.028, 0.372] (q=0.367) | -0.145  [-0.366, 0.076] (q=0.278) | -0.321  [-0.566, -0.076] (q=0.083) | -0.336  [-0.620, -0.052] (q=0.133) | -0.347  [-0.632, -0.062] (q=0.138) |
| Full-scale IQ | 163.740  [-27.145, 354.625] (q=0.163) | 0.472  [-1.009, 1.952] (q=0.979) | -1.342  [-2.970, 0.286] (q=0.278) | -1.067  [-2.886, 0.753] (q=0.399) | -1.826  [-3.923, 0.271] (q=0.175) | -1.812  [-3.919, 0.296] (q=0.183) |
| Verbal Comprehension | 148.198  [-40.038, 336.435] (q=0.163) | -0.102  [-1.561, 1.358] (q=0.979) | -1.459  [-3.062, 0.144] (q=0.278) | -1.279  [-3.070, 0.512] (q=0.399) | -2.064  [-4.127, -0.001] (q=0.133) | -2.056  [-4.129, 0.018] (q=0.139) |
| Perceptual Reasoning | 80.580  [-106.045, 267.205] (q=0.396) | -0.019  [-1.464, 1.426] (q=0.979) | -0.206  [-1.800, 1.388] (q=0.800) | -0.649  [-2.426, 1.128] (q=0.541) | -0.604  [-2.657, 1.449] (q=0.563) | -0.631  [-2.694, 1.432] (q=0.548) |
| General Memory | 180.217  [-37.390, 397.824] (q=0.163) | 0.197  [-1.476, 1.870] (q=0.979) | -1.179  [-3.020, 0.661] (q=0.278) | -0.449  [-2.507, 1.608] (q=0.668) | -1.288  [-3.670, 1.094] (q=0.329) | -1.246  [-3.640, 1.148] (q=0.351) |
| Processing Speed | 183.539  [-16.269, 383.348] (q=0.163) | 1.478  [-0.067, 3.022] (q=0.367) | -1.667  [-3.371, 0.036] (q=0.278) | -1.368  [-3.274, 0.537] (q=0.399) | -2.297  [-4.490, -0.104] (q=0.133) | -2.282  [-4.486, -0.077] (q=0.139) |
| Reading Composite Score | 106.761  [-79.612, 293.134] (q=0.298) | 0.149  [-1.275, 1.573] (q=0.979) | -1.074  [-2.643, 0.494] (q=0.278) | -0.665  [-2.417, 1.088] (q=0.541) | -1.345  [-3.373, 0.684] (q=0.257) | -1.321  [-3.360, 0.717] (q=0.271) |
| Math Composite Score | 281.592  [33.672, 529.512] (q=0.163) | 0.957  [-0.945, 2.860] (q=0.861) | -1.133  [-3.234, 0.968] (q=0.331) | -1.524  [-3.865, 0.816] (q=0.399) | -1.971  [-4.683, 0.740] (q=0.246) | -1.997  [-4.722, 0.728] (q=0.241) |

**Supplementary Table 9. Associations between regional LH anomaly scores and ND outcomes in CHD.** Associations between each regional (lobar) LH anomaly score and neurodevelopmental outcomes in the CHD cohort, estimated by GLMs adjusting for age, sex, socioeconomic status, and cognitive test type (WISC or WAIS, according to age). Rows denote the eight ND outcomes assessed; columns are organized by anomaly-scoring method and, within each method, by cortical lobe (frontal, temporal, parietal, occipital). Each cell reports the standardized regression coefficient $\beta$, its 95% CI, and the FDR-corrected q-value. A negative $\beta$ indicates that higher anomaly scores are associated with poorer outcome. FDR correction (Benjamini–Hochberg) was applied across the eight outcomes within each method; cells shaded in light orange denote q < 0.05.

|  | **SGM** | | | | **OC-SVM** | | | | **Recon Error** | | | | **GNN Logit** | | | | **COMPASS_1** | | | | **COMPASS_2** | | | |
| --- | --- | --- | --- | --- | --- | --- | --- | --- | --- | --- | --- | --- | --- | --- | --- | --- | --- | --- | --- | --- | --- | --- | --- | --- |
| **ND Outcome** | **LF** | **LT** | **LP** | **LO** | **LF** | **LT** | **LP** | **LO** | **LF** | **LT** | **LP** | **LO** | **LF** | **LT** | **LP** | **LO** | **LF** | **LT** | **LP** | **LO** | **LF** | **LT** | **LP** | **LO** |
| Executive Function | 5.745 [-17.215, 28.706] (q=0.623) | 8.734 [-4.913, 22.380] (q=0.527) | 5.304 [-13.083, 23.692] (q=0.979) | **21.490 [6.434, 36.546] (q=0.042)** | -0.191 [-0.402, 0.020] (q=0.321) | 0.149 [-0.063, 0.361] (q=0.927) | 0.213 [0.003, 0.423] (q=0.074) | -0.033 [-0.259, 0.192] (q=0.982) | -0.104 [-0.324, 0.116] (q=0.469) | 0.011 [-0.196, 0.218] (q=0.999) | -0.023 [-0.236, 0.191] (q=0.979) | -0.065 [-0.278, 0.148] (q=0.733) | -0.012 [-0.214, 0.190] (q=0.930) | 0.017 [-0.198, 0.232] (q=0.974) | -0.161 [-0.381, 0.059] (q=0.242) | 0.040 [-0.165, 0.246] (q=0.794) | -0.091 [-0.364, 0.182] (q=0.585) | 0.029 [-0.277, 0.336] (q=0.955) | -0.151 [-0.431, 0.130] (q=0.390) | -0.018 [-0.290, 0.255] (q=0.963) | -0.114 [-0.376, 0.147] (q=0.451) | 0.021 [-0.248, 0.290] (q=0.979) | -0.082 [-0.341, 0.176] (q=0.723) | -0.053 [-0.309, 0.203] (q=0.786) |
| Full-scale IQ | 120.305 [-46.272, 286.881] (q=0.382) | 40.180 [-59.263, 139.623] (q=0.527) | -5.468 [-139.318, 128.382] (q=0.979) | 93.671 [-16.772, 204.114] (q=0.247) | 0.881 [-0.660, 2.422] (q=0.550) | 0.280 [-1.265, 1.825] (q=0.927) | 1.906 [0.385, 3.426] (q=0.057) | -0.411 [-2.053, 1.231] (q=0.982) | -1.366 [-2.962, 0.230] (q=0.249) | -0.153 [-1.656, 1.350] (q=0.999) | -0.169 [-1.721, 1.384] (q=0.979) | -0.996 [-2.544, 0.552] (q=0.550) | -0.450 [-1.917, 1.017] (q=0.930) | 0.240 [-1.323, 1.803] (q=0.974) | -1.529 [-3.124, 0.067] (q=0.161) | 0.761 [-0.732, 2.253] (q=0.750) | -1.461 [-3.442, 0.519] (q=0.295) | 0.076 [-2.153, 2.306] (q=0.955) | -1.390 [-3.431, 0.651] (q=0.290) | -0.144 [-2.127, 1.839] (q=0.963) | -1.626 [-3.524, 0.271] (q=0.186) | -0.094 [-2.051, 1.864] (q=0.979) | -0.730 [-2.609, 1.150] (q=0.723) | -0.757 [-2.621, 1.106] (q=0.786) |
| Verbal Comprehension | 98.732 [-65.824, 263.287] (q=0.382) | 49.236 [-48.847, 147.318] (q=0.527) | 49.617 [-82.360, 181.594] (q=0.979) | 96.317 [-12.633, 205.268] (q=0.247) | 0.493 [-1.030, 2.016] (q=0.600) | 0.415 [-1.110, 1.940] (q=0.927) | **2.136 [0.640, 3.633] (q=0.042)** | -1.146 [-2.762, 0.470] (q=0.896) | -0.433 [-2.014, 1.149] (q=0.591) | 0.497 [-0.986, 1.979] (q=0.999) | 0.263 [-1.269, 1.795] (q=0.979) | -0.210 [-1.741, 1.322] (q=0.888) | 0.065 [-1.383, 1.513] (q=0.930) | 0.025 [-1.517, 1.568] (q=0.974) | -0.950 [-2.530, 0.630] (q=0.279) | 1.389 [-0.079, 2.856] (q=0.508) | -0.273 [-2.233, 1.688] (q=0.785) | 0.572 [-1.627, 2.772] (q=0.955) | -0.545 [-2.564, 1.474] (q=0.596) | 1.050 [-0.903, 3.004] (q=0.753) | -0.424 [-2.305, 1.457] (q=0.658) | 0.634 [-1.297, 2.565] (q=0.979) | -0.054 [-1.911, 1.802] (q=0.954) | 0.338 [-1.503, 2.179] (q=0.786) |
| Perceptual Reasoning | 81.007 [-81.314, 243.329] (q=0.436) | 36.309 [-60.442, 133.060] (q=0.527) | -32.220 [-162.380, 97.940] (q=0.979) | 53.737 [-54.018, 161.491] (q=0.374) | 1.333 [-0.161, 2.828] (q=0.321) | 0.625 [-0.877, 2.127] (q=0.927) | 0.441 [-1.052, 1.934] (q=0.562) | -0.046 [-1.644, 1.552] (q=0.982) | -0.916 [-2.472, 0.641] (q=0.397) | -0.074 [-1.536, 1.388] (q=0.999) | -0.746 [-2.255, 0.762] (q=0.979) | -0.651 [-2.159, 0.858] (q=0.635) | -0.405 [-1.832, 1.022] (q=0.930) | -0.260 [-1.780, 1.261] (q=0.974) | -1.328 [-2.882, 0.225] (q=0.187) | 0.656 [-0.797, 2.109] (q=0.750) | -1.074 [-3.003, 0.855] (q=0.439) | -0.346 [-2.514, 1.823] (q=0.955) | -1.729 [-3.711, 0.252] (q=0.290) | 0.046 [-1.883, 1.975] (q=0.963) | -1.135 [-2.986, 0.715] (q=0.365) | -0.199 [-2.103, 1.705] (q=0.979) | -1.286 [-3.110, 0.539] (q=0.666) | -0.430 [-2.244, 1.384] (q=0.786) |
| General Memory | 150.165 [-40.247, 340.578] (q=0.382) | 29.428 [-84.384, 143.240] (q=0.611) | 107.464 [-45.161, 260.089] (q=0.979) | 71.458 [-55.184, 198.100] (q=0.357) | 0.837 [-0.927, 2.600] (q=0.550) | 0.283 [-1.485, 2.050] (q=0.927) | 1.524 [-0.224, 3.271] (q=0.100) | -0.467 [-2.345, 1.411] (q=0.982) | **-2.465 [-4.278, -0.652] (q=0.031)** | 0.001 [-1.718, 1.721] (q=0.999) | -1.719 [-3.484, 0.047] (q=0.451) | -1.503 [-3.271, 0.264] (q=0.381) | 0.112 [-1.567, 1.791] (q=0.930) | 0.070 [-1.718, 1.858] (q=0.974) | -1.087 [-2.918, 0.744] (q=0.279) | -0.416 [-2.126, 1.293] (q=0.794) | -1.790 [-4.054, 0.474] (q=0.295) | 0.073 [-2.478, 2.623] (q=0.955) | -2.377 [-4.703, -0.051] (q=0.290) | -1.593 [-3.855, 0.668] (q=0.753) | -2.529 [-4.692, -0.367] (q=0.088) | 0.030 [-2.209, 2.269] (q=0.979) | -2.255 [-4.392, -0.118] (q=0.309) | -1.776 [-3.901, 0.349] (q=0.607) |
| Processing Speed | 69.512 [-105.741, 244.765] (q=0.498) | 46.051 [-58.313, 150.414] (q=0.527) | 2.154 [-138.348, 142.656] (q=0.979) | 81.948 [-34.143, 198.038] (q=0.265) | 0.892 [-0.725, 2.510] (q=0.550) | -0.153 [-1.776, 1.469] (q=0.927) | 1.803 [0.204, 3.401] (q=0.073) | -0.092 [-1.816, 1.632] (q=0.982) | **-2.572 [-4.231, -0.914] (q=0.020)** | -1.186 [-2.758, 0.386] (q=0.999) | 0.022 [-1.608, 1.652] (q=0.979) | -1.431 [-3.053, 0.190] (q=0.381) | -1.278 [-2.812, 0.256] (q=0.519) | 0.300 [-1.340, 1.941] (q=0.974) | -1.871 [-3.542, -0.200] (q=0.117) | 0.260 [-1.310, 1.829] (q=0.794) | **-3.146 [-5.202, -1.091] (q=0.023)** | -0.999 [-3.337, 1.339] (q=0.955) | -1.504 [-3.646, 0.638] (q=0.290) | -0.940 [-3.019, 1.139] (q=0.753) | **-3.251 [-5.219, -1.284] (q=0.010)** | -1.366 [-3.415, 0.683] (q=0.979) | -0.645 [-2.619, 1.328] (q=0.723) | -1.425 [-3.377, 0.527] (q=0.607) |
| Reading Composite Score | 102.772 [-62.109, 267.653] (q=0.382) | 37.820 [-60.538, 136.179] (q=0.527) | 1.803 [-130.575, 134.182] (q=0.979) | 45.089 [-64.515, 154.693] (q=0.419) | -0.228 [-1.755, 1.299] (q=0.769) | -0.071 [-1.600, 1.457] (q=0.927) | 1.425 [-0.085, 2.935] (q=0.086) | -1.003 [-2.624, 0.617] (q=0.896) | -0.658 [-2.242, 0.927] (q=0.474) | 0.177 [-1.309, 1.664] (q=0.999) | 0.048 [-1.488, 1.584] (q=0.979) | -0.110 [-1.645, 1.425] (q=0.888) | -0.292 [-1.743, 1.159] (q=0.930) | -0.338 [-1.884, 1.207] (q=0.974) | -0.812 [-2.396, 0.772] (q=0.314) | 0.920 [-0.555, 2.395] (q=0.750) | -0.772 [-2.735, 1.191] (q=0.585) | -0.149 [-2.354, 2.056] (q=0.955) | -0.619 [-2.642, 1.404] (q=0.596) | 0.719 [-1.241, 2.679] (q=0.753) | -0.816 [-2.699, 1.068] (q=0.451) | 0.085 [-1.851, 2.021] (q=0.979) | -0.238 [-2.099, 1.622] (q=0.916) | 0.255 [-1.590, 2.100] (q=0.786) |
| Math Composite Score | 216.369 [-4.297, 437.036] (q=0.382) | 62.420 [-69.625, 194.465] (q=0.527) | 55.461 [-122.231, 233.153] (q=0.979) | 115.167 [-31.634, 261.969] (q=0.247) | 0.854 [-1.195, 2.903] (q=0.550) | -0.190 [-2.243, 1.863] (q=0.927) | 2.162 [0.137, 4.187] (q=0.073) | 0.025 [-2.157, 2.207] (q=0.982) | -1.590 [-3.713, 0.532] (q=0.283) | -0.239 [-2.235, 1.758] (q=0.999) | 0.061 [-2.002, 2.123] (q=0.979) | -0.939 [-2.998, 1.120] (q=0.635) | -1.500 [-3.442, 0.443] (q=0.519) | -0.436 [-2.512, 1.640] (q=0.974) | -2.352 [-4.467, -0.238] (q=0.117) | -0.264 [-2.250, 1.722] (q=0.794) | -2.595 [-5.219, 0.028] (q=0.210) | -0.706 [-3.667, 2.255] (q=0.955) | -1.861 [-4.572, 0.849] (q=0.290) | -0.999 [-3.631, 1.633] (q=0.753) | -2.322 [-4.841, 0.198] (q=0.186) | -0.477 [-3.077, 2.123] (q=0.979) | -0.775 [-3.273, 1.723] (q=0.723) | -1.111 [-3.586, 1.364] (q=0.786) |

**Supplementary Table 10. Associations between regional RH anomaly scores and ND outcomes in CHD.** Associations between each regional (lobar) RH anomaly score and neurodevelopmental outcomes in the CHD cohort, estimated by GLMs adjusting for age, sex, socioeconomic status, and cognitive test type (WISC or WAIS, according to age). Rows denote the eight ND outcomes assessed; columns are organized by anomaly-scoring method and, within each method, by cortical lobe (frontal, temporal, parietal, occipital). Each cell reports the standardized regression coefficient $\beta$, its 95% CI, and the FDR-corrected q-value. A negative $\beta$ indicates that higher anomaly scores are associated with poorer outcome. FDR correction (Benjamini–Hochberg) was applied across the eight outcomes within each method; cells shaded in light orange denote q < 0.05.

|  | **SGM** | | | | **OC-SVM** | | | | **Recon Error** | | | | **GNN Logit** | | | | **COMPASS_1** | | | | **COMPASS_2** | | | |
| --- | --- | --- | --- | --- | --- | --- | --- | --- | --- | --- | --- | --- | --- | --- | --- | --- | --- | --- | --- | --- | --- | --- | --- | --- |
| **ND Outcome** | **RF** | **RT** | **RP** | **RO** | **RF** | **RT** | **RP** | **RO** | **RF** | **RT** | **RP** | **RO** | **RF** | **RT** | **RP** | **RO** | **RF** | **RT** | **RP** | **RO** | **RF** | **RT** | **RP** | **RO** |
| Executive Function | 6.659 [-16.092, 29.410] (q=0.565) | 8.751 [-4.863, 22.365] (q=0.535) | 4.414 [-13.911, 22.740] (q=0.955) | 20.563 [5.558, 35.569] (q=0.059) | 0.280 [0.054, 0.506] (q=0.061) | 0.129 [-0.089, 0.348] (q=0.653) | -0.084 [-0.317, 0.149] (q=0.900) | -0.022 [-0.252, 0.208] (q=0.852) | -0.166 [-0.386, 0.053] (q=0.183) | -0.260 [-0.486, -0.035] (q=0.152) | -0.033 [-0.253, 0.188] (q=0.934) | 0.120 [-0.103, 0.342] (q=0.760) | -0.093 [-0.294, 0.107] (q=0.360) | -0.248 [-0.478, -0.019] (q=0.253) | -0.033 [-0.254, 0.188] (q=0.960) | 0.112 [-0.104, 0.327] (q=0.442) | -0.242 [-0.531, 0.047] (q=0.115) | **-0.431 [-0.726, -0.137] (q=0.034)** | -0.060 [-0.357, 0.238] (q=0.929) | 0.193 [-0.089, 0.476] (q=0.375) | -0.234 [-0.520, 0.052] (q=0.125) | **-0.429 [-0.723, -0.134] (q=0.035)** | -0.059 [-0.356, 0.237] (q=0.903) | 0.191 [-0.090, 0.473] (q=0.363) |
| Full-scale IQ | 125.028 [-40.287, 290.344] (q=0.367) | 38.742 [-60.640, 138.124] (q=0.535) | -14.542 [-148.140, 119.056] (q=0.955) | 87.237 [-22.969, 197.443] (q=0.275) | 1.810 [0.162, 3.459] (q=0.075) | 0.462 [-1.131, 2.056] (q=0.690) | -0.337 [-2.034, 1.360] (q=0.900) | -0.362 [-2.036, 1.312] (q=0.805) | -1.750 [-3.343, -0.157] (q=0.079) | -1.015 [-2.669, 0.640] (q=0.328) | -0.340 [-1.944, 1.264] (q=0.934) | 0.640 [-0.981, 2.260] (q=0.760) | -1.306 [-2.761, 0.149] (q=0.213) | -1.429 [-3.106, 0.249] (q=0.253) | -0.276 [-1.886, 1.334] (q=0.960) | 1.007 [-0.563, 2.576] (q=0.442) | **-2.888 [-4.979, -0.797] (q=0.028)** | -2.065 [-4.228, 0.099] (q=0.099) | -0.560 [-2.727, 1.607] (q=0.929) | 1.384 [-0.675, 3.442] (q=0.375) | **-2.819 [-4.890, -0.748] (q=0.031)** | -2.085 [-4.246, 0.076] (q=0.103) | -0.553 [-2.715, 1.609] (q=0.903) | 1.400 [-0.650, 3.450] (q=0.363) |
| Verbal Comprehension | 89.358 [-74.139, 252.855] (q=0.377) | 45.013 [-53.072, 143.099] (q=0.535) | 41.395 [-90.438, 173.227] (q=0.955) | 92.043 [-16.705, 200.792] (q=0.275) | 1.731 [0.102, 3.359] (q=0.075) | -0.357 [-1.931, 1.216] (q=0.690) | -1.233 [-2.904, 0.437] (q=0.900) | -0.319 [-1.972, 1.334] (q=0.805) | -1.750 [-3.323, -0.178] (q=0.079) | -0.965 [-2.599, 0.669] (q=0.328) | -0.067 [-1.652, 1.517] (q=0.934) | 0.443 [-1.158, 2.044] (q=0.760) | -0.813 [-2.254, 0.628] (q=0.345) | -1.496 [-3.152, 0.159] (q=0.253) | -0.718 [-2.306, 0.869] (q=0.960) | 0.974 [-0.576, 2.524] (q=0.442) | -2.372 [-4.444, -0.299] (q=0.054) | -2.078 [-4.214, 0.058] (q=0.099) | -0.712 [-2.851, 1.427] (q=0.929) | 1.197 [-0.837, 3.230] (q=0.396) | -2.272 [-4.325, -0.220] (q=0.057) | -2.107 [-4.240, 0.026] (q=0.103) | -0.756 [-2.890, 1.378] (q=0.903) | 1.225 [-0.800, 3.250] (q=0.376) |
| Perceptual Reasoning | 97.898 [-63.287, 259.083] (q=0.373) | 36.824 [-59.957, 133.605] (q=0.535) | -39.762 [-169.791, 90.267] (q=0.955) | 48.548 [-59.051, 156.147] (q=0.429) | 1.005 [-0.608, 2.618] (q=0.232) | 0.396 [-1.156, 1.948] (q=0.690) | -0.106 [-1.759, 1.547] (q=0.900) | 0.600 [-1.030, 2.229] (q=0.805) | -0.456 [-2.018, 1.106] (q=0.566) | -0.039 [-1.654, 1.575] (q=0.962) | 0.067 [-1.496, 1.630] (q=0.934) | 0.287 [-1.292, 1.866] (q=0.760) | -0.892 [-2.312, 0.529] (q=0.345) | -1.023 [-2.660, 0.614] (q=0.293) | 0.077 [-1.491, 1.645] (q=0.960) | 0.625 [-0.905, 2.156] (q=0.482) | -1.330 [-3.385, 0.725] (q=0.204) | -0.887 [-3.003, 1.230] (q=0.431) | 0.131 [-1.980, 2.242] (q=0.929) | 0.770 [-1.238, 2.779] (q=0.515) | -1.346 [-3.380, 0.689] (q=0.194) | -0.950 [-3.064, 1.164] (q=0.431) | 0.131 [-1.975, 2.237] (q=0.903) | 0.788 [-1.212, 2.788] (q=0.501) |
| General Memory | 160.431 [-28.585, 349.446] (q=0.367) | 32.121 [-81.664, 145.906] (q=0.579) | 102.863 [-49.607, 255.332] (q=0.955) | 60.928 [-65.502, 187.359] (q=0.429) | 1.523 [-0.370, 3.416] (q=0.152) | 0.521 [-1.303, 2.344] (q=0.690) | 0.718 [-1.223, 2.659] (q=0.900) | -1.975 [-3.879, -0.071] (q=0.337) | -1.911 [-3.736, -0.086] (q=0.079) | -0.832 [-2.727, 1.064] (q=0.444) | -0.438 [-2.273, 1.398] (q=0.934) | 1.524 [-0.325, 3.373] (q=0.760) | -0.954 [-2.625, 0.716] (q=0.345) | -0.338 [-2.266, 1.591] (q=0.731) | -0.047 [-1.890, 1.796] (q=0.960) | 0.889 [-0.909, 2.687] (q=0.442) | -2.659 [-5.063, -0.256] (q=0.054) | -0.998 [-3.485, 1.490] (q=0.431) | -0.442 [-2.922, 2.038] (q=0.929) | 1.999 [-0.353, 4.351] (q=0.375) | -2.555 [-4.936, -0.175] (q=0.057) | -0.961 [-3.446, 1.525] (q=0.447) | -0.412 [-2.886, 2.063] (q=0.903) | 1.945 [-0.397, 4.288] (q=0.363) |
| Processing Speed | 79.973 [-93.754, 253.700] (q=0.418) | 48.453 [-55.710, 152.616] (q=0.535) | -3.978 [-144.069, 136.113] (q=0.955) | 74.273 [-41.435, 189.981] (q=0.332) | 2.216 [0.492, 3.940] (q=0.061) | 1.060 [-0.607, 2.727] (q=0.653) | 0.258 [-1.522, 2.038] (q=0.900) | -1.074 [-2.825, 0.678] (q=0.805) | -1.722 [-3.394, -0.050] (q=0.079) | -1.396 [-3.128, 0.335] (q=0.227) | -0.451 [-2.132, 1.231] (q=0.934) | 0.264 [-1.437, 1.965] (q=0.760) | -1.860 [-3.380, -0.341] (q=0.132) | -1.150 [-2.913, 0.612] (q=0.293) | 0.105 [-1.583, 1.793] (q=0.960) | 0.206 [-1.444, 1.856] (q=0.806) | **-3.443 [-5.628, -1.258] (q=0.017)** | -2.161 [-4.430, 0.108] (q=0.099) | -0.316 [-2.589, 1.957] (q=0.929) | 0.391 [-1.773, 2.555] (q=0.722) | **-3.411 [-5.574, -1.247] (q=0.017)** | -2.137 [-4.403, 0.130] (q=0.103) | -0.274 [-2.542, 1.993] (q=0.903) | 0.385 [-1.770, 2.540] (q=0.726) |
| Reading Composite Score | 104.386 [-58.863, 267.634] (q=0.373) | 36.230 [-61.819, 134.279] (q=0.535) | -4.628 [-136.430, 127.174] (q=0.955) | 42.575 [-66.464, 151.613] (q=0.443) | 0.996 [-0.639, 2.630] (q=0.232) | -0.319 [-1.891, 1.254] (q=0.690) | -0.911 [-2.583, 0.760] (q=0.900) | -0.765 [-2.415, 0.885] (q=0.805) | -1.578 [-3.152, -0.005] (q=0.079) | -1.361 [-2.990, 0.267] (q=0.227) | -0.174 [-1.757, 1.408] (q=0.934) | 0.321 [-1.279, 1.921] (q=0.760) | -0.757 [-2.197, 0.683] (q=0.345) | -0.311 [-1.973, 1.351] (q=0.731) | -0.371 [-1.959, 1.217] (q=0.960) | 0.807 [-0.742, 2.357] (q=0.442) | -2.163 [-4.237, -0.090] (q=0.055) | -1.431 [-3.572, 0.709] (q=0.252) | -0.495 [-2.633, 1.642] (q=0.929) | 0.954 [-1.079, 2.988] (q=0.475) | -2.075 [-4.129, -0.022] (q=0.064) | -1.355 [-3.494, 0.784] (q=0.285) | -0.507 [-2.640, 1.626] (q=0.903) | 0.981 [-1.044, 3.006] (q=0.455) |
| Math Composite Score | 229.348 [11.103, 447.593] (q=0.316) | 64.246 [-67.317, 195.809] (q=0.535) | 50.809 [-126.067, 227.684] (q=0.955) | 110.533 [-35.486, 256.551] (q=0.275) | 1.859 [-0.331, 4.049] (q=0.152) | 1.322 [-0.784, 3.429] (q=0.653) | 0.304 [-1.944, 2.552] (q=0.900) | -0.594 [-2.811, 1.624] (q=0.805) | -1.392 [-3.512, 0.728] (q=0.226) | -2.309 [-4.490, -0.127] (q=0.152) | 0.519 [-1.605, 2.644] (q=0.934) | 0.606 [-1.542, 2.753] (q=0.760) | -1.722 [-3.649, 0.205] (q=0.213) | -1.561 [-3.786, 0.665] (q=0.293) | -0.667 [-2.799, 1.464] (q=0.960) | 1.557 [-0.520, 3.634] (q=0.442) | -3.012 [-5.794, -0.230] (q=0.054) | -3.289 [-6.147, -0.430] (q=0.097) | -0.131 [-3.002, 2.741] (q=0.929) | 1.830 [-0.897, 4.556] (q=0.375) | -2.999 [-5.754, -0.245] (q=0.057) | -3.226 [-6.083, -0.370] (q=0.103) | -0.216 [-3.080, 2.649] (q=0.903) | 1.882 [-0.833, 4.597] (q=0.363) |

**Supplementary Table 11. Associations between global LH anomaly scores and ND outcomes after adjusting for surface quality proxy (Euler number) in CHD.** Associations between each LH anomaly score and ND outcomes in the CHD cohort, estimated by GLMs adjusting for age, sex, socioeconomic status, cognitive test type (WISC or WAIS, according to age), and surface quality proxy (Euler number). Rows denote the ND outcomes assessed; columns denote the six anomaly-scoring methods compared. Each cell reports the standardized regression coefficient $\beta$, its 95% CI, and the FDR-corrected q-value. A negative $\beta$ indicates that higher anomaly scores are associated with poorer outcome. FDR correction (Benjamini–Hochberg) was applied across the eight outcomes within each method; asterisks denote *q < 0.05.

| **ND Outcome** | **SGM** | **OCSVM** | **Recon Error** | **GNN Logit** | **COMPASS_1** | **COMPASS_2** |
| --- | --- | --- | --- | --- | --- | --- |
| Executive Function | 23.880  [-2.416, 50.176] (q=0.150) | -0.016  [-0.256, 0.225] (q=0.925) | -0.084  [-0.285, 0.116] (q=0.546) | 0.001  [-0.200, 0.202] (q=0.993) | -0.065  [-0.316, 0.186] (q=0.813) | -0.084  [-0.324, 0.156] (q=0.655) |
| Full-scale IQ | 166.658  [-24.922, 358.238] (q=0.150) | 0.965  [-0.792, 2.722] (q=0.770) | -1.188  [-2.650, 0.275] (q=0.296) | -0.472  [-1.906, 0.963] (q=0.993) | -1.289  [-3.102, 0.524] (q=0.325) | -1.395  [-3.141, 0.350] (q=0.216) |
| Verbal Comprehension | 161.274  [-27.451, 350.000] (q=0.150) | 0.491  [-1.242, 2.224] (q=0.770) | -0.130  [-1.576, 1.316] (q=0.859) | 0.058  [-1.356, 1.472] (q=0.993) | -0.054  [-1.845, 1.737] (q=0.953) | -0.104  [-1.830, 1.622] (q=0.905) |
| Perceptual Reasoning | 72.770  [-114.416, 259.957] (q=0.445) | 0.851  [-0.844, 2.545] (q=0.770) | -0.921  [-2.348, 0.505] (q=0.328) | -0.840  [-2.231, 0.551] (q=0.658) | -1.380  [-3.142, 0.382] (q=0.325) | -1.295  [-2.994, 0.405] (q=0.216) |
| General Memory | 175.111  [-43.310, 393.531] (q=0.154) | 0.576  [-1.414, 2.565] (q=0.770) | -1.981  [-3.635, -0.327] (q=0.076) | 0.239  [-1.383, 1.861] (q=0.993) | -1.321  [-3.378, 0.737] (q=0.332) | -1.865  [-3.845, 0.114] (q=0.216) |
| Processing Speed | 181.209  [-19.626, 382.045] (q=0.150) | 1.368  [-0.454, 3.190] (q=0.770) | -2.195  [-3.717, -0.674] (q=0.039*) | -1.083  [-2.580, 0.414] (q=0.658) | -2.549  [-4.434, -0.665] (q=0.065) | -2.672  [-4.485, -0.858] (q=0.032*) |
| Reading Composite Score | 110.187  [-77.051, 297.425] (q=0.283) | -0.081  [-1.775, 1.612] (q=0.925) | -0.486  [-1.908, 0.935] (q=0.573) | 0.063  [-1.362, 1.488] (q=0.993) | -0.331  [-2.108, 1.447] (q=0.817) | -0.460  [-2.162, 1.241] (q=0.680) |
| Math Composite Score | 273.669  [24.181, 523.157] (q=0.150) | 0.980  [-1.285, 3.246] (q=0.770) | -1.359  [-3.258, 0.541] (q=0.321) | -1.124  [-3.028, 0.781] (q=0.658) | -1.934  [-4.306, 0.437] (q=0.325) | -1.842  [-4.113, 0.428] (q=0.216) |

**Supplementary Table 12. Associations between global RH anomaly scores and ND outcomes after adjusting for surface quality proxy (Euler number) in CHD.** Associations between each RH anomaly score and ND outcomes in the CHD cohort, estimated by GLMs adjusting for age, sex, socioeconomic status, cognitive test type (WISC or WAIS, according to age), and surface quality proxy (Euler number). Rows denote the ND outcomes assessed; columns denote the six anomaly-scoring methods compared. Each cell reports the standardized regression coefficient $\beta$, its 95% CI, and the FDR-corrected q-value. A negative $\beta$ indicates that higher anomaly scores are associated with poorer outcome. FDR correction (Benjamini–Hochberg) was applied across the eight outcomes within each method.

| **ND Outcome** | **SGM** | **OCSVM** | **Recon Error** | **GNN Logit** | **COMPASS_1** | **COMPASS_2** |
| --- | --- | --- | --- | --- | --- | --- |
| Executive Function | 22.054  [-4.232, 48.339] (q=0.189) | 0.172  [-0.027, 0.371] (q=0.363) | -0.125  [-0.347, 0.098] (q=0.378) | -0.307  [-0.552, -0.061] (q=0.116) | -0.312  [-0.599, -0.026] (q=0.232) | -0.324  [-0.611, -0.036] (q=0.221) |
| Full-scale IQ | 144.238  [-47.124, 335.601] (q=0.189) | 0.440  [-1.033, 1.914] (q=0.949) | -1.143  [-2.781, 0.495] (q=0.378) | -0.905  [-2.724, 0.915] (q=0.526) | -1.561  [-3.672, 0.550] (q=0.294) | -1.548  [-3.669, 0.573] (q=0.304) |
| Verbal Comprehension | 143.425  [-46.216, 333.065] (q=0.189) | -0.112  [-1.573, 1.350] (q=0.949) | -1.423  [-3.043, 0.197] (q=0.378) | -1.237  [-3.038, 0.564] (q=0.503) | -2.018  [-4.106, 0.069] (q=0.232) | -2.009  [-4.106, 0.088] (q=0.241) |
| Perceptual Reasoning | 63.910  [-123.184, 251.004] (q=0.502) | -0.047  [-1.487, 1.394] (q=0.949) | -0.017  [-1.620, 1.586] (q=0.984) | -0.508  [-2.287, 1.271] (q=0.656) | -0.357  [-2.424, 1.710] (q=0.734) | -0.387  [-2.463, 1.690] (q=0.714) |
| General Memory | 163.525  [-55.003, 382.052] (q=0.189) | 0.191  [-1.478, 1.860] (q=0.949) | -1.012  [-2.867, 0.842] (q=0.378) | -0.307  [-2.368, 1.754] (q=0.770) | -1.056  [-3.457, 1.345] (q=0.443) | -1.015  [-3.427, 1.398] (q=0.467) |
| Processing Speed | 153.956  [-44.756, 352.669] (q=0.189) | 1.427  [-0.099, 2.953] (q=0.363) | -1.354  [-3.055, 0.347] (q=0.378) | -1.117  [-3.009, 0.775] (q=0.503) | -1.880  [-4.072, 0.312] (q=0.247) | -1.867  [-4.070, 0.335] (q=0.257) |
| Reading Composite Score | 94.229  [-92.707, 281.165] (q=0.368) | 0.146  [-1.275, 1.567] (q=0.949) | -0.950  [-2.529, 0.630] (q=0.378) | -0.552  [-2.309, 1.204] (q=0.656) | -1.170  [-3.214, 0.874] (q=0.348) | -1.147  [-3.201, 0.907] (q=0.364) |
| Math Composite Score | 265.057  [16.376, 513.738] (q=0.189) | 0.953  [-0.945, 2.851] (q=0.864) | -0.934  [-3.048, 1.180] (q=0.440) | -1.370  [-3.715, 0.976] (q=0.503) | -1.712  [-4.444, 1.021] (q=0.348) | -1.740  [-4.484, 1.005] (q=0.341) |

**Supplementary Table 13. Associations between anomaly scores and ordinal language outcome in the PMG cohort.** Associations between each anomaly score and the three-level ordinal language outcome (1, intact; 2, mild-to-moderate impairment; 3, profound impairment) in the PMG cohort, assessed by KW tests. Rows denote the scored regions. Columns denote the anomaly-scoring methods compared. Each cell reports the KW H statistic and the FDR-corrected q-value; a higher H reflects stronger separation of anomaly scores across the three language-severity groups. FDR correction (Benjamini–Hochberg) was applied across the five regions within each hemisphere (the whole hemisphere and its four lobes); asterisks denote *q < 0.05, **q < 0.01.

| **ND Outcome** | **Recon Error** | **GNN Logit** | **COMPASS_1** | **COMPASS_2** |
| --- | --- | --- | --- | --- |
| LH | 4.080 (q=0.132) | 7.164 (q=0.020*) | 8.155 (q=0.010*) | 8.105 (q=0.008**) |
| LF | 5.530 (q=0.112) | 9.730 (q=0.014*) | 9.900 (q=0.009**) | 9.900 (q=0.008**) |
| LT | 2.560 (q=0.296) | 3.120 (q=0.425) | 3.270 (q=0.269) | 4.200 (q=0.164) |
| LP | 5.650 (q=0.112) | 1.340 (q=0.536) | 7.670 (q=0.025*) | 5.510 (q=0.110) |
| LO | 3.050 (q=0.296) | 1.390 (q=0.536) | 1.590 (q=0.471) | 1.410 (q=0.517) |
| RH | 4.186 (q=0.121) | 4.224 (q=0.125) | 3.129 (q=0.221) | 3.882 (q=0.148) |
| RF | 3.940 (q=0.425) | 8.230 (q=0.031*) | 9.010 (q=0.016*) | 8.930 (q=0.017*) |
| RT | 1.410 (q=0.518) | 1.140 (q=0.780) | 1.050 (q=0.621) | 1.020 (q=0.623) |
| RP | 2.390 (q=0.425) | 2.330 (q=0.660) | 2.580 (q=0.544) | 2.850 (q=0.505) |
| RO | 2.400 (q=0.425) | 0.430 (q=0.826) | 1.910 (q=0.544) | 1.590 (q=0.623) |

**Supplementary Table 14. Cohort-wise ICC between ID and OoD anomaly scores under LOCO validation.** Agreement between ID and OoD anomaly scores for each of the eleven normative cohorts, quantified by the ICC. Under the LOCO design, the ID score is obtained when all eleven cohorts contribute to training and the OoD score when the subject's own cohort is held out, both computed on the same subjects; a higher ICC therefore indicates that a cohort's scores are more stable when that cohort is unseen during training. The table is divided into three vertically stacked blocks corresponding to the three anomaly scores (reconstruction error, GNN logit, COMPASS_1), each listing the eleven cohorts as rows. Columns report, for LH and RH separately, the anomaly score type, cohort, number of subjects (N), and the ICC with its lower and upper 95% confidence bounds. Cohorts with smaller N yield correspondingly wider CIs. The per-cohort values summarized here underlie the mean ICCs reported in Fig. 5 and the scatter plots in Supplementary Fig. 1.

| **Anomaly Score** | **Cohort** | **N** | **ICC** | **95% CI_low** | **95% CI_high** | **Anomaly Score** | **Cohort** | **N** | **ICC** | **95% CI_low** | **95% CI_high** |
| --- | --- | --- | --- | --- | --- | --- | --- | --- | --- | --- | --- |
| **LH Recon Error** | HCP | 55 | 0.820 | 0.71 | 0.89 | **RH Recon Error** | HCP | 55 | 0.914 | 0.86 | 0.95 |
|  | HCPD | 30 | 0.853 | 0.72 | 0.93 |  | HCPD | 31 | 0.900 | 0.79 | 0.95 |
|  | CHCP | 13 | 0.754 | 0.32 | 0.92 |  | CHCP | 13 | 0.684 | 0.24 | 0.89 |
|  | CMI | 65 | 0.871 | 0.80 | 0.92 |  | CMI | 64 | 0.901 | 0.84 | 0.94 |
|  | BGSP | 78 | 0.998 | 0.99 | 1.00 |  | BGSP | 78 | 0.854 | 0.78 | 0.90 |
|  | BeijingEN | 9 | 0.822 | 0.38 | 0.96 |  | BeijingEN | 9 | 0.873 | 0.55 | 0.97 |
|  | PING | 36 | 0.904 | 0.82 | 0.95 |  | PING | 36 | 0.883 | 0.78 | 0.94 |
|  | Qtim | 50 | 0.841 | 0.62 | 0.92 |  | Qtim | 48 | 0.903 | 0.81 | 0.95 |
|  | SWU | 28 | 0.825 | 0.65 | 0.92 |  | SWU | 28 | 0.839 | 0.68 | 0.92 |
|  | Narratives | 17 | 0.872 | 0.69 | 0.95 |  | Narratives | 17 | 0.580 | 0.16 | 0.82 |
|  | ABCD | 98 | 0.866 | 0.81 | 0.91 |  | ABCD | 97 | 0.722 | 0.61 | 0.81 |
| **LH GNN Logit** | HCP | 55 | 0.517 | 0.29 | 0.69 | **RH GNN Logit** | HCP | 55 | 0.739 | 0.59 | 0.84 |
|  | HCPD | 30 | 0.622 | 0.35 | 0.80 |  | HCPD | 31 | 0.418 | 0.09 | 0.67 |
|  | CHCP | 13 | 0.610 | 0.10 | 0.86 |  | CHCP | 13 | 0.572 | 0.06 | 0.85 |
|  | CMI | 65 | 0.452 | 0.24 | 0.63 |  | CMI | 64 | 0.629 | 0.45 | 0.76 |
|  | BGSP | 78 | 0.901 | 0.83 | 0.94 |  | BGSP | 78 | 0.665 | 0.52 | 0.77 |
|  | BeijingEN | 9 | -0.035 | -0.76 | 0.63 |  | BeijingEN | 9 | 0.646 | -0.01 | 0.91 |
|  | PING | 36 | 0.707 | 0.50 | 0.84 |  | PING | 36 | 0.874 | 0.77 | 0.93 |
|  | Qtim | 50 | 0.301 | 0.02 | 0.53 |  | Qtim | 48 | 0.806 | 0.68 | 0.89 |
|  | SWU | 28 | 0.756 | 0.54 | 0.88 |  | SWU | 28 | 0.809 | 0.63 | 0.91 |
|  | Narratives | 17 | 0.584 | 0.15 | 0.83 |  | Narratives | 17 | 0.723 | 0.38 | 0.89 |
|  | ABCD | 98 | 0.720 | 0.61 | 0.80 |  | ABCD | 97 | 0.525 | 0.36 | 0.66 |
| **LH COMPASS_1** | HCP | 55 | 0.723 | 0.57 | 0.83 | **RH COMPASS_1** | HCP | 55 | 0.911 | 0.85 | 0.95 |
|  | HCPD | 30 | 0.695 | 0.45 | 0.84 |  | HCPD | 31 | 0.756 | 0.54 | 0.88 |
|  | CHCP | 13 | 0.828 | 0.53 | 0.94 |  | CHCP | 13 | 0.663 | 0.19 | 0.88 |
|  | CMI | 65 | 0.767 | 0.64 | 0.85 |  | CMI | 64 | 0.824 | 0.73 | 0.89 |
|  | BGSP | 78 | 0.957 | 0.91 | 0.98 |  | BGSP | 78 | 0.781 | 0.68 | 0.85 |
|  | BeijingEN | 9 | 0.705 | 0.14 | 0.92 |  | BeijingEN | 9 | 0.855 | 0.50 | 0.96 |
|  | PING | 36 | 0.887 | 0.78 | 0.94 |  | PING | 36 | 0.914 | 0.84 | 0.96 |
|  | Qtim | 50 | 0.547 | 0.13 | 0.76 |  | Qtim | 48 | 0.811 | 0.67 | 0.89 |
|  | SWU | 28 | 0.819 | 0.64 | 0.91 |  | SWU | 28 | 0.824 | 0.66 | 0.91 |
|  | Narratives | 17 | 0.754 | 0.44 | 0.90 |  | Narratives | 17 | 0.883 | 0.71 | 0.96 |
|  | ABCD | 98 | 0.840 | 0.77 | 0.89 |  | ABCD | 97 | 0.693 | 0.57 | 0.78 |

**Supplementary Table 15. Cohort-wise OoD bias under LOCO validation.** Bias values between ID and OoD anomaly scores for each of the eleven normative cohorts, quantified by the difference (OoD – ID). A higher bias indicates that a cohort's OoD score are more higher than that of ID score for each subject. Columns report, for LH and RH separately, the anomaly score type, cohort, number of subjects (N), the mean and SD of bias with its RMSE.

| **Anomaly Score** | **Cohort** | **N** | **Mean bias** | **SD bias** | **RMSE** | **Anomaly Score** | **Cohort** | **N** | **Mean bias** | **SD**  **bias** | **RMSE** |
| --- | --- | --- | --- | --- | --- | --- | --- | --- | --- | --- | --- |
| **LH Recon Error** | HCP | 55 | 0.084 | 0.461 | 0.613 | **RH Recon Error** | HCP | 55 | 0.063 | 0.430 | 0.628 |
|  | HCPD | 30 | 0.073 | 0.542 | 0.809 |  | HCPD | 31 | 0.206 | 0.539 | 0.619 |
|  | CHCP | 13 | 0.322 | 0.495 | 0.934 |  | CHCP | 13 | -0.080 | 0.458 | 0.474 |
|  | CMI | 65 | 0.045 | 0.549 | 0.700 |  | CMI | 64 | -0.083 | 0.534 | 0.646 |
|  | BGSP | 78 | 0.032 | 0.049 | 0.053 |  | BGSP | 78 | 0.061 | 0.562 | 0.711 |
|  | BeijingEN | 9 | 0.277 | 0.434 | 0.630 |  | BeijingEN | 9 | 0.130 | 0.571 | 0.382 |
|  | PING | 36 | -0.137 | 0.456 | 0.702 |  | PING | 36 | -0.091 | 0.464 | 0.735 |
|  | Qtim | 50 | 0.307 | 0.493 | 0.614 |  | Qtim | 48 | 0.155 | 0.375 | 0.472 |
|  | SWU | 28 | 0.170 | 0.471 | 0.590 |  | SWU | 28 | -0.041 | 0.551 | 0.654 |
|  | Narratives | 17 | -0.137 | 0.463 | 0.513 |  | Narratives | 17 | -0.097 | 0.528 | 0.560 |
|  | ABCD | 98 | -0.063 | 0.500 | 0.661 |  | ABCD | 97 | 0.079 | 0.577 | 0.731 |
| **LH GNN Logit** | HCP | 55 | -0.050 | 0.670 | 0.901 | **RH GNN Logit** | HCP | 55 | -0.020 | 0.709 | 0.749 |
|  | HCPD | 30 | 0.383 | 1.129 | 1.474 |  | HCPD | 31 | 0.249 | 1.028 | 0.960 |
|  | CHCP | 13 | 0.124 | 1.045 | 0.501 |  | CHCP | 13 | 0.223 | 1.147 | 0.587 |
|  | CMI | 65 | -0.308 | 1.039 | 1.162 |  | CMI | 64 | -0.041 | 0.780 | 0.797 |
|  | BGSP | 78 | 0.188 | 0.471 | 0.501 |  | BGSP | 78 | 0.155 | 0.952 | 0.991 |
|  | BeijingEN | 9 | -0.120 | 0.717 | 1.169 |  | BeijingEN | 9 | 0.055 | 0.555 | 0.623 |
|  | PING | 36 | -0.147 | 0.697 | 0.813 |  | PING | 36 | -0.082 | 0.530 | 0.775 |
|  | Qtim | 50 | 0.742 | 1.222 | 1.696 |  | Qtim | 48 | 0.156 | 0.642 | 0.763 |
|  | SWU | 28 | 0.102 | 0.644 | 0.835 |  | SWU | 28 | -0.071 | 0.612 | 0.767 |
|  | Narratives | 17 | 0.111 | 1.047 | 0.717 |  | Narratives | 17 | 0.111 | 0.791 | 0.813 |
|  | ABCD | 98 | -0.015 | 0.806 | 0.976 |  | ABCD | 97 | -0.050 | 0.978 | 1.121 |
| **LH COMPASS_1** | HCP | 55 | 0.017 | 0.423 | 0.560 | **RH COMPASS_1** | HCP | 55 | 0.022 | 0.341 | 0.471 |
|  | HCPD | 30 | 0.228 | 0.665 | 0.914 |  | HCPD | 31 | 0.228 | 0.606 | 0.623 |
|  | CHCP | 13 | 0.223 | 0.462 | 0.401 |  | CHCP | 13 | 0.072 | 0.614 | 0.361 |
|  | CMI | 65 | -0.132 | 0.569 | 0.702 |  | CMI | 64 | -0.062 | 0.486 | 0.562 |
|  | BGSP | 78 | 0.110 | 0.235 | 0.199 |  | BGSP | 78 | 0.108 | 0.565 | 0.659 |
|  | BeijingEN | 9 | 0.078 | 0.359 | 0.451 |  | BeijingEN | 9 | 0.092 | 0.417 | 0.481 |
|  | PING | 36 | -0.142 | 0.389 | 0.407 |  | PING | 36 | -0.086 | 0.351 | 0.443 |
|  | Qtim | 50 | 0.524 | 0.674 | 0.870 |  | Qtim | 48 | 0.155 | 0.421 | 0.628 |
|  | SWU | 28 | 0.136 | 0.408 | 0.432 |  | SWU | 28 | -0.056 | 0.496 | 0.599 |
|  | Narratives | 17 | -0.013 | 0.543 | 0.528 |  | Narratives | 17 | 0.007 | 0.337 | 0.515 |
|  | ABCD | 98 | -0.039 | 0.459 | 0.588 |  | ABCD | 97 | 0.015 | 0.537 | 0.589 |

**Supplementary Table 16. Effects of surface quality on OoD score bias.** Effect of cortical surface quality on the LOCO bias, estimated by LMMs with cohort as a random intercept, fitted separately for each anomaly score (reconstruction error, GNN logit, COMPASS_1) and hemisphere. The outcome is the OoD bias (OoD - ID), and the predictor is the surface-quality proxy (negative-log-transformed FreeSurfer Euler number, higher values denoting poorer quality). Each cell reports the standardized coefficient $\beta$ for the quality term, its 95% CI, and the FDR-corrected q-value. FDR correction (Benjamini–Hochberg) was applied across the three scores within each hemisphere.

| **Anomaly Score** | **Beta**  **(Euler)** | **95% CI_low** | **95% CI_high** | **FDR**  **q** | **Anomaly Score** | **Beta (Euler)** | **95% CI_low** | **95% CI_high** | **FDR**  **q** |
| --- | --- | --- | --- | --- | --- | --- | --- | --- | --- |
| LH  Recon Error | 0.0622 | -0.0006 | 0.1251 | 0.1565 | RH  Recon Error | 0.0622 | -0.0006 | 0.1251 | 0.1565 |
| LH  GNN Logit | -0.0628 | -0.2051 | 0.0796 | 0.4473 | RH  GNN Logit | -0.0628 | -0.2051 | 0.0796 | 0.4473 |
| LH COMPASS | 0.0291 | -0.046 | 0.1042 | 0.4473 | RH COMPASS | 0.0291 | -0.046 | 0.1042 | 0.4473 |

**Supplementary Table 17. Effects of age and sex on anomaly scores.** Fixed effects of age and sex on each anomaly score, estimated by LMMs with cohort as a random intercept, fitted separately for each hemisphere. Rows list the three anomaly for LH and RH. For each of age and sex, the table reports the standardized coefficient $\beta$, its 95% CI, and the FDR-corrected q-value; a negative age coefficient indicates lower scores in older subjects, and the sex coefficient is relative to male subjects. FDR correction (Benjamini-Hochberg) was applied across all metric × predictor combinations within each hemisphere. These main-effect models exclude the age-by-sex interaction; because that interaction was significant only for the GNN logit in RH (Supplementary Table 23), the main effects reported here for the RH GNN logit should not be interpreted in isolation.

| **Metric** |  | **Beta** | **FDR**  **q** | **95% CI_low** | **95% CI_High** | **Metric** |  | **Beta** | **FDR**  **q** | **95% CI_low** | **95% CI_High** |
| --- | --- | --- | --- | --- | --- | --- | --- | --- | --- | --- | --- |
| LH COMPASS | sex | -0.070 | 0.401 | -0.212 | 0.072 | RH COMPASS | sex | -0.075 | 0.417 | -0.210 | 0.061 |
|  | age | -0.032 | 0.018* | -0.054 | -0.009 |  | age | -0.020 | 0.154 | -0.043 | 0.002 |
| LH  GNN Logit | sex | -0.039 | 0.683 | -0.229 | 0.150 | RH  GNN Logit | sex | -0.177 | 0.125 | -0.347 | -0.007 |
|  | age | -0.010 | 0.365 | -0.024 | 0.005 |  | age | -0.005 | 0.761 | -0.034 | 0.023 |
| LH  Recon Error | sex | -0.101 | 0.368 | -0.270 | 0.069 | RH  Recon Error | sex | 0.027 | 0.761 | -0.147 | 0.201 |
|  | age | -0.054 | 0.001* | -0.081 | -0.027 |  | age | -0.036 | 0.101 | -0.065 | -0.006 |

**Supplementary Table 18. Effects of age, sex, and surface quality on anomaly scores.** Fixed effects of age, sex, and cortical surface quality on each anomaly score, estimated by LMMs with cohort as a random intercept, fitted separately for each hemisphere. This specification extends the age-and-sex models of Supplementary Table 20 by adding the surface-quality proxy (negative-log-transformed FreeSurfer Euler number, higher values denoting poorer quality) as a third fixed effect. Rows list the three anomaly scores for LH and RH. For each of age, sex, and quality, the table reports the standardized coefficient $\beta$, its 95% CI, and the FDR-corrected q-value; the sex coefficient is relative to male subjects. FDR correction (Benjamini-Hochberg) was applied across all metric × predictor combinations within each hemisphere. Comparing the age coefficients with those of the quality-free models (Supplementary Table 20) isolates the fraction of the age dependence attributable to surface quality.

| **Metric** |  | **Beta** | **FDR**  **q** | **95% CI_low** | **95% CI_High** | **Metric** |  | **Beta** | **FDR**  **q** | **95% CI_low** | **95% CI_High** |
| --- | --- | --- | --- | --- | --- | --- | --- | --- | --- | --- | --- |
| LH COMPASS | sex | -0.075 | 0.386 | -0.216 | 0.067 | RH COMPASS | sex | -0.087 | 0.267 | -0.222 | 0.048 |
|  | age | -0.027 | 0.042* | -0.050 | -0.005 |  | age | -0.018 | 0.191 | -0.040 | 0.005 |
|  | euler | 0.159 | 0.033* | 0.037 | 0.282 |  | euler | 0.166 | 0.061 | 0.040 | 0.292 |
| LH  GNN Logit | sex | -0.042 | 0.654 | -0.227 | 0.143 | RH  GNN Logit | sex | -0.191 | 0.062 | -0.361 | -0.021 |
|  | age | -0.007 | 0.654 | -0.037 | 0.023 |  | age | -0.002 | 0.898 | -0.030 | 0.027 |
|  | euler | 0.100 | 0.336 | -0.061 | 0.260 |  | euler | 0.200 | 0.061 | 0.041 | 0.358 |
| LH  Recon Error | sex | -0.107 | 0.336 | -0.276 | 0.061 | RH  Recon Error | sex | 0.018 | 0.898 | -0.157 | 0.192 |
|  | age | -0.048 | 0.005** | -0.075 | -0.021 |  | age | -0.033 | 0.062 | -0.063 | -0.004 |
|  | euler | 0.219 | 0.015* | 0.072 | 0.365 |  | euler | 0.133 | 0.191 | -0.030 | 0.296 |

**Supplementary Table 19. Effects of age, sex, surface quality, and hemisphere on anomaly scores.** Fixed effects of age, sex, cortical surface quality, hemisphere, and cohort on each anomaly score, estimated by LMMs with individual as a random intercept. This specification extends the age-sex-quality models of Supplementary Table 21 by adding the hemisphere as a fourth effect. Rows list the three anomaly scores. For each of age, sex, quality, and hemisphere, the table reports the standardized coefficient $\beta$, its 95% CI, and the FDR-corrected q-value; the sex coefficient is relative to male subjects and the hemisphere is relative to LH. FDR correction (Benjamini-Hochberg) was applied across all metric × predictor combinations.

| **Metric** |  | **Beta** | **FDR**  **q** | **95% CI_low** | **95% CI_High** |
| --- | --- | --- | --- | --- | --- |
| COMPASS | sex | -0.075 | 0.231 | -0.173 | 0.023 |
|  | age | -0.024 | 0.011* | -0.040 | -0.008 |
|  | euler | 0.162 | 0.002** | 0.075 | 0.250 |
|  | hemi | -0.066 | 0.251 | -0.160 | 0.028 |
| GNN Logit | sex | -0.105 | 0.209 | -0.231 | 0.022 |
|  | age | -0.004 | 0.697 | -0.025 | 0.017 |
|  | euler | 0.144 | 0.030* | 0.031 | 0.257 |
|  | hemi | -0.066 | 0.346 | -0.189 | 0.056 |
| Recon Error | sex | -0.047 | 0.491 | -0.170 | 0.075 |
|  | age | -0.043 | <0.001*** | -0.063 | -0.023 |
|  | euler | 0.177 | 0.006** | 0.068 | 0.286 |
|  | hemi | -0.067 | 0.346 | -0.185 | 0.051 |

**Supplementary Table 20. Age-by-sex interaction effects on anomaly scores.** Age-by-sex interaction effects on each anomaly score, estimated by LMMs with cohort as a random intercept and age, sex, and their interaction as fixed effects, fitted separately for each hemisphere. Rows list the three anomaly scores for LH and RH; each cell reports the standardized interaction coefficient $\beta$, its 95% CI, and the FDR-corrected q-value. FDR correction (Benjamini-Hochberg) was applied across the three anomaly scores within each hemisphere. The interaction was tested first to determine model structure: it was significant only for the RH GNN logit, which was therefore retained in its full form, whereas for all other score-hemisphere combinations the non-significant interaction was dropped and only main effects were estimated (Supplementary Tables 20-22)

| **Metric** | **Beta** | **FDR**  **q** | **95% CI_low** | **95% CI_High** | **Metric** | **Beta** | **FDR**  **q** | **95% CI_low** | **95% CI_High** |
| --- | --- | --- | --- | --- | --- | --- | --- | --- | --- |
| LH COMPASS | 0.011 | 0.425 | -0.008 | 0.030 | RH COMPASS | -0.016 | 0.173 | -0.035 | 0.004 |
| LH  GNN Logit | 0.011 | 0.425 | -0.014 | 0.037 | RH  GNN Logit | -0.037 | 0.009** | -0.062 | -0.013 |
| LH  Recon Error | 0.010 | 0.425 | -0.014 | 0.033 | RH  Recon Error | 0.005 | 0.699 | -0.020 | 0.030 |

**Supplementary Table 21. Post-hoc pairwise cohort comparisons of reconstruction error across the normative cohorts.** Post-hoc pairwise comparisons of the reconstruction error between all pairs of the eleven normative cohorts, following the omnibus ANOVA reported in Fig. 6. Rows denote the 55 unique cohort pairs; columns report, for the LH and RH separately, the mean difference between the two cohorts, its 95% CI, and the FDR-corrected q-value. P values are Tukey-adjusted for all 55 pairwise comparisons. Cells shaded in light orange denote p < 0.05.

| **LH** | | | | | | **RH** | | | | | |
| --- | --- | --- | --- | --- | --- | --- | --- | --- | --- | --- | --- |
| **Group1** | **Group2** | **Mean Diff** | **P value** | **95% CI_low** | **95% CI_High** | **Group1** | **Group2** | **Mean Diff** | **P**  **value** | **95% CI_low** | **95% CI_High** |
| ABCD | BGSP | -0.599 | 0.002 | -1.070 | -0.128 | ABCD | BGSP | -0.036 | 1.000 | -0.505 | 0.433 |
| ABCD | BeijingEN | -1.148 | 0.027 | -2.229 | -0.068 | ABCD | BeijingEN | -0.719 | 0.526 | -1.790 | 0.352 |
| ABCD | CHCP | -0.613 | 0.531 | -1.529 | 0.303 | ABCD | CHCP | -0.969 | 0.025 | -1.876 | -0.061 |
| ABCD | CMI | -0.134 | 0.999 | -0.633 | 0.365 | ABCD | CMI | -0.104 | 1.000 | -0.599 | 0.391 |
| ABCD | HCP | -0.783 | 0.000 | -1.306 | -0.261 | ABCD | HCP | -0.510 | 0.056 | -1.025 | 0.006 |
| ABCD | HCP-D | -0.522 | 0.231 | -1.161 | 0.117 | ABCD | HCP-D | -0.270 | 0.954 | -0.904 | 0.364 |
| ABCD | Narratives | -0.584 | 0.424 | -1.399 | 0.231 | ABCD | Narratives | 0.103 | 1.000 | -0.705 | 0.911 |
| ABCD | PING | -0.126 | 1.000 | -0.731 | 0.478 | ABCD | PING | 0.423 | 0.448 | -0.177 | 1.023 |
| ABCD | QTIM | -0.659 | 0.004 | -1.198 | -0.119 | ABCD | QTIM | -0.614 | 0.011 | -1.152 | -0.075 |
| ABCD | SWU | -0.441 | 0.544 | -1.106 | 0.224 | ABCD | SWU | -0.105 | 1.000 | -0.774 | 0.564 |
| BGSP | BeijingEN | -0.550 | 0.870 | -1.642 | 0.543 | BGSP | BeijingEN | -0.683 | 0.620 | -1.766 | 0.399 |
| BGSP | CHCP | -0.014 | 1.000 | -0.944 | 0.915 | BGSP | CHCP | -0.933 | 0.044 | -1.854 | -0.011 |
| BGSP | CMI | 0.465 | 0.135 | -0.059 | 0.988 | BGSP | CMI | -0.068 | 1.000 | -0.588 | 0.452 |
| BGSP | HCP | -0.185 | 0.991 | -0.731 | 0.362 | BGSP | HCP | -0.474 | 0.147 | -1.013 | 0.066 |
| BGSP | HCP-D | 0.077 | 1.000 | -0.582 | 0.735 | BGSP | HCP-D | -0.234 | 0.987 | -0.887 | 0.420 |
| BGSP | Narratives | 0.015 | 1.000 | -0.816 | 0.845 | BGSP | Narratives | 0.139 | 1.000 | -0.684 | 0.963 |
| BGSP | PING | 0.472 | 0.341 | -0.153 | 1.097 | BGSP | PING | 0.459 | 0.373 | -0.161 | 1.080 |
| BGSP | QTIM | -0.060 | 1.000 | -0.622 | 0.502 | BGSP | QTIM | -0.578 | 0.038 | -1.139 | -0.016 |
| BGSP | SWU | 0.158 | 1.000 | -0.526 | 0.841 | BGSP | SWU | -0.069 | 1.000 | -0.756 | 0.618 |
| BeijingEN | CHCP | 0.535 | 0.971 | -0.810 | 1.881 | BeijingEN | CHCP | -0.249 | 1.000 | -1.582 | 1.083 |
| BeijingEN | CMI | 1.014 | 0.105 | -0.090 | 2.119 | BeijingEN | CMI | 0.615 | 0.769 | -0.479 | 1.709 |
| BeijingEN | HCP | 0.365 | 0.993 | -0.751 | 1.481 | BeijingEN | HCP | 0.210 | 1.000 | -0.894 | 1.313 |
| BeijingEN | HCP-D | 0.626 | 0.823 | -0.549 | 1.801 | BeijingEN | HCP-D | 0.450 | 0.976 | -0.714 | 1.613 |
| BeijingEN | Narratives | 0.564 | 0.941 | -0.715 | 1.843 | BeijingEN | Narratives | 0.823 | 0.578 | -0.444 | 2.089 |
| BeijingEN | PING | 1.022 | 0.140 | -0.135 | 2.178 | BeijingEN | PING | 1.142 | 0.051 | -0.003 | 2.287 |
| BeijingEN | QTIM | 0.490 | 0.946 | -0.634 | 1.613 | BeijingEN | QTIM | 0.105 | 1.000 | -1.009 | 1.220 |
| BeijingEN | SWU | 0.707 | 0.702 | -0.482 | 1.896 | BeijingEN | SWU | 0.614 | 0.846 | -0.569 | 1.797 |
| CHCP | CMI | 0.479 | 0.864 | -0.465 | 1.423 | CHCP | CMI | 0.864 | 0.100 | -0.070 | 1.799 |
| CHCP | HCP | -0.170 | 1.000 | -1.127 | 0.786 | CHCP | HCP | 0.459 | 0.895 | -0.487 | 1.405 |
| CHCP | HCP-D | 0.091 | 1.000 | -0.934 | 1.116 | CHCP | HCP-D | 0.699 | 0.487 | -0.316 | 1.714 |
| CHCP | Narratives | 0.029 | 1.000 | -1.114 | 1.172 | CHCP | Narratives | 1.072 | 0.082 | -0.060 | 2.204 |
| CHCP | PING | 0.486 | 0.895 | -0.518 | 1.490 | CHCP | PING | 1.392 | 0.000 | 0.397 | 2.386 |
| CHCP | QTIM | -0.046 | 1.000 | -1.012 | 0.920 | CHCP | QTIM | 0.355 | 0.983 | -0.604 | 1.313 |
| CHCP | SWU | 0.172 | 1.000 | -0.870 | 1.213 | CHCP | SWU | 0.863 | 0.206 | -0.174 | 1.901 |
| CMI | HCP | -0.649 | 0.012 | -1.220 | -0.079 | CMI | HCP | -0.406 | 0.413 | -0.968 | 0.157 |
| CMI | HCP-D | -0.388 | 0.750 | -1.067 | 0.291 | CMI | HCP-D | -0.166 | 0.999 | -0.838 | 0.507 |
| CMI | Narratives | -0.450 | 0.825 | -1.297 | 0.397 | CMI | Narratives | 0.208 | 0.999 | -0.631 | 1.046 |
| CMI | PING | 0.008 | 1.000 | -0.639 | 0.654 | CMI | PING | 0.527 | 0.220 | -0.113 | 1.167 |
| CMI | QTIM | -0.525 | 0.127 | -1.110 | 0.061 | CMI | QTIM | -0.510 | 0.151 | -1.093 | 0.074 |
| CMI | SWU | -0.307 | 0.945 | -1.010 | 0.396 | CMI | SWU | -0.001 | 1.000 | -0.706 | 0.704 |
| HCP | HCP-D | 0.261 | 0.981 | -0.436 | 0.958 | HCP | HCP-D | 0.240 | 0.989 | -0.448 | 0.928 |
| HCP | Narratives | 0.199 | 1.000 | -0.662 | 1.060 | HCP | Narratives | 0.613 | 0.415 | -0.238 | 1.464 |
| HCP | PING | 0.657 | 0.056 | -0.008 | 1.322 | HCP | PING | 0.933 | 0.000 | 0.276 | 1.589 |
| HCP | QTIM | 0.125 | 1.000 | -0.482 | 0.731 | HCP | QTIM | -0.104 | 1.000 | -0.705 | 0.497 |
| HCP | SWU | 0.342 | 0.907 | -0.378 | 1.062 | HCP | SWU | 0.405 | 0.769 | -0.315 | 1.125 |
| HCP-D | Narratives | -0.062 | 1.000 | -0.998 | 0.874 | HCP-D | Narratives | 0.373 | 0.968 | -0.554 | 1.300 |
| HCP-D | PING | 0.396 | 0.844 | -0.365 | 1.156 | HCP-D | PING | 0.693 | 0.104 | -0.060 | 1.446 |
| HCP-D | QTIM | -0.137 | 1.000 | -0.846 | 0.573 | HCP-D | QTIM | -0.344 | 0.891 | -1.049 | 0.361 |
| HCP-D | SWU | 0.081 | 1.000 | -0.728 | 0.890 | HCP-D | SWU | 0.164 | 1.000 | -0.644 | 0.973 |
| Narratives | PING | 0.458 | 0.873 | -0.455 | 1.371 | Narratives | PING | 0.320 | 0.988 | -0.585 | 1.224 |
| Narratives | QTIM | -0.075 | 1.000 | -0.946 | 0.797 | Narratives | QTIM | -0.717 | 0.211 | -1.582 | 0.148 |
| Narratives | SWU | 0.143 | 1.000 | -0.811 | 1.097 | Narratives | SWU | -0.209 | 1.000 | -1.160 | 0.743 |
| PING | QTIM | -0.532 | 0.285 | -1.210 | 0.146 | PING | QTIM | -1.037 | 0.000 | -1.711 | -0.362 |
| PING | SWU | -0.315 | 0.968 | -1.097 | 0.467 | PING | SWU | -0.528 | 0.517 | -1.311 | 0.254 |
| QTIM | SWU | 0.217 | 0.997 | -0.515 | 0.950 | QTIM | SWU | 0.509 | 0.482 | -0.228 | 1.245 |

**Supplementary Table 22. Post-hoc pairwise cohort comparisons of GNN logit across the normative cohorts.** Post-hoc pairwise comparisons of the GNN logit between all pairs of the eleven normative cohorts, following the omnibus ANOVA reported in Fig. 6. Rows denote the 55 unique cohort pairs; columns report, for the LH and RH separately, the mean difference between the two cohorts, its 95% CI, and the FDR-corrected q-value. P values are Tukey-adjusted for all 55 pairwise comparisons. Cells shaded in light orange denote p < 0.05.

| **LH** | | | | | | **RH** | | | | | |
| --- | --- | --- | --- | --- | --- | --- | --- | --- | --- | --- | --- |
| **Group1** | **Group2** | **Mean Diff** | **P**  **value** | **95% CI_low** | **95% CI_High** | **Group1** | **Group2** | **Mean Diff** | **P**  **value** | **95% CI_low** | **95% CI_High** |
| ABCD | BGSP | 0.288 | 0.748 | -0.215 | 0.791 | ABCD | BGSP | -0.090 | 1.000 | -0.548 | 0.367 |
| ABCD | BeijingEN | -0.134 | 1.000 | -1.289 | 1.022 | ABCD | BeijingEN | -0.084 | 1.000 | -1.129 | 0.961 |
| ABCD | CHCP | 0.289 | 0.997 | -0.690 | 1.267 | ABCD | CHCP | -0.276 | 0.995 | -1.162 | 0.610 |
| ABCD | CMI | 0.131 | 0.999 | -0.402 | 0.664 | ABCD | CMI | 0.140 | 0.998 | -0.343 | 0.623 |
| ABCD | HCP | -0.050 | 1.000 | -0.609 | 0.508 | ABCD | HCP | 0.050 | 1.000 | -0.454 | 0.553 |
| ABCD | HCP-D | 0.361 | 0.832 | -0.323 | 1.044 | ABCD | HCP-D | 0.259 | 0.959 | -0.360 | 0.877 |
| ABCD | Narratives | -0.099 | 1.000 | -0.971 | 0.772 | ABCD | Narratives | -0.145 | 1.000 | -0.934 | 0.643 |
| ABCD | PING | 0.200 | 0.996 | -0.447 | 0.846 | ABCD | PING | 0.070 | 1.000 | -0.515 | 0.656 |
| ABCD | QTIM | 0.084 | 1.000 | -0.492 | 0.661 | ABCD | QTIM | 0.403 | 0.319 | -0.123 | 0.929 |
| ABCD | SWU | 0.201 | 0.998 | -0.509 | 0.912 | ABCD | SWU | -0.121 | 1.000 | -0.773 | 0.532 |
| BGSP | BeijingEN | -0.422 | 0.986 | -1.589 | 0.746 | BGSP | BeijingEN | 0.007 | 1.000 | -1.050 | 1.063 |
| BGSP | CHCP | 0.001 | 1.000 | -0.993 | 0.994 | BGSP | CHCP | -0.186 | 1.000 | -1.085 | 0.714 |
| BGSP | CMI | -0.157 | 0.998 | -0.717 | 0.402 | BGSP | CMI | 0.230 | 0.930 | -0.277 | 0.737 |
| BGSP | HCP | -0.338 | 0.734 | -0.922 | 0.246 | BGSP | HCP | 0.140 | 0.999 | -0.387 | 0.667 |
| BGSP | HCP-D | 0.073 | 1.000 | -0.632 | 0.777 | BGSP | HCP-D | 0.349 | 0.797 | -0.289 | 0.987 |
| BGSP | Narratives | -0.387 | 0.945 | -1.275 | 0.500 | BGSP | Narratives | -0.055 | 1.000 | -0.858 | 0.749 |
| BGSP | PING | -0.089 | 1.000 | -0.757 | 0.580 | BGSP | PING | 0.161 | 0.999 | -0.445 | 0.766 |
| BGSP | QTIM | -0.204 | 0.991 | -0.805 | 0.397 | BGSP | QTIM | 0.493 | 0.122 | -0.055 | 1.041 |
| BGSP | SWU | -0.087 | 1.000 | -0.817 | 0.644 | BGSP | SWU | -0.030 | 1.000 | -0.701 | 0.640 |
| BeijingEN | CHCP | 0.422 | 0.997 | -1.016 | 1.860 | BeijingEN | CHCP | -0.193 | 1.000 | -1.493 | 1.108 |
| BeijingEN | CMI | 0.264 | 1.000 | -0.916 | 1.445 | BeijingEN | CMI | 0.223 | 1.000 | -0.844 | 1.291 |
| BeijingEN | HCP | 0.083 | 1.000 | -1.109 | 1.276 | BeijingEN | HCP | 0.133 | 1.000 | -0.944 | 1.210 |
| BeijingEN | HCP-D | 0.494 | 0.973 | -0.762 | 1.750 | BeijingEN | HCP-D | 0.342 | 0.997 | -0.793 | 1.478 |
| BeijingEN | Narratives | 0.034 | 1.000 | -1.333 | 1.401 | BeijingEN | Narratives | -0.062 | 1.000 | -1.298 | 1.175 |
| BeijingEN | PING | 0.333 | 0.999 | -0.903 | 1.569 | BeijingEN | PING | 0.154 | 1.000 | -0.964 | 1.272 |
| BeijingEN | QTIM | 0.218 | 1.000 | -0.983 | 1.419 | BeijingEN | QTIM | 0.487 | 0.936 | -0.601 | 1.574 |
| BeijingEN | SWU | 0.335 | 0.999 | -0.936 | 1.606 | BeijingEN | SWU | 0.614 | 0.846 | -0.569 | 1.797 |
| CHCP | CMI | -0.158 | 1.000 | -1.167 | 0.851 | CHCP | CMI | 0.864 | 0.100 | -0.070 | 1.799 |
| CHCP | HCP | -0.339 | 0.993 | -1.362 | 0.684 | CHCP | HCP | 0.459 | 0.895 | -0.487 | 1.405 |
| CHCP | HCP-D | 0.072 | 1.000 | -1.024 | 1.168 | CHCP | HCP-D | 0.699 | 0.487 | -0.316 | 1.714 |
| CHCP | Narratives | -0.388 | 0.995 | -1.610 | 0.834 | CHCP | Narratives | 1.072 | 0.082 | -0.060 | 2.204 |
| CHCP | PING | -0.089 | 1.000 | -1.162 | 0.984 | CHCP | PING | 1.392 | 0.000 | 0.397 | 2.386 |
| CHCP | QTIM | -0.204 | 1.000 | -1.237 | 0.828 | CHCP | QTIM | 0.355 | 0.983 | -0.604 | 1.313 |
| CHCP | SWU | -0.087 | 1.000 | -1.200 | 1.026 | CHCP | SWU | 0.863 | 0.206 | -0.174 | 1.901 |
| CMI | HCP | -0.181 | 0.997 | -0.791 | 0.429 | CMI | HCP | -0.406 | 0.413 | -0.968 | 0.157 |
| CMI | HCP-D | 0.230 | 0.995 | -0.496 | 0.956 | CMI | HCP-D | -0.166 | 0.999 | -0.838 | 0.507 |
| CMI | Narratives | -0.230 | 0.999 | -1.135 | 0.675 | CMI | Narratives | 0.208 | 0.999 | -0.631 | 1.046 |
| CMI | PING | 0.069 | 1.000 | -0.622 | 0.760 | CMI | PING | 0.527 | 0.220 | -0.113 | 1.167 |
| CMI | QTIM | -0.047 | 1.000 | -0.673 | 0.579 | CMI | QTIM | -0.510 | 0.151 | -1.093 | 0.074 |
| CMI | SWU | 0.071 | 1.000 | -0.681 | 0.822 | CMI | SWU | -0.001 | 1.000 | -0.706 | 0.704 |
| HCP | HCP-D | 0.411 | 0.789 | -0.334 | 1.156 | HCP | HCP-D | 0.240 | 0.989 | -0.448 | 0.928 |
| HCP | Narratives | -0.049 | 1.000 | -0.969 | 0.871 | HCP | Narratives | 0.613 | 0.415 | -0.238 | 1.464 |
| HCP | PING | 0.250 | 0.988 | -0.461 | 0.961 | HCP | PING | 0.933 | 0.000 | 0.276 | 1.589 |
| HCP | QTIM | 0.135 | 1.000 | -0.514 | 0.783 | HCP | QTIM | -0.104 | 1.000 | -0.705 | 0.497 |
| HCP | SWU | 0.252 | 0.993 | -0.518 | 1.022 | HCP | SWU | 0.405 | 0.769 | -0.315 | 1.125 |
| HCP-D | Narratives | -0.460 | 0.924 | -1.461 | 0.541 | HCP-D | Narratives | 0.373 | 0.968 | -0.554 | 1.300 |
| HCP-D | PING | -0.161 | 1.000 | -0.974 | 0.652 | HCP-D | PING | 0.693 | 0.104 | -0.060 | 1.446 |
| HCP-D | QTIM | -0.277 | 0.984 | -1.035 | 0.482 | HCP-D | QTIM | -0.344 | 0.891 | -1.049 | 0.361 |
| HCP-D | SWU | -0.159 | 1.000 | -1.024 | 0.705 | HCP-D | SWU | 0.164 | 1.000 | -0.644 | 0.973 |
| Narratives | PING | 0.299 | 0.996 | -0.677 | 1.275 | Narratives | PING | 0.320 | 0.988 | -0.585 | 1.224 |
| Narratives | QTIM | 0.184 | 1.000 | -0.748 | 1.115 | Narratives | QTIM | -0.717 | 0.211 | -1.582 | 0.148 |
| Narratives | SWU | 0.301 | 0.997 | -0.719 | 1.320 | Narratives | SWU | -0.209 | 1.000 | -1.160 | 0.743 |
| PING | QTIM | -0.115 | 1.000 | -0.840 | 0.610 | PING | QTIM | -1.037 | 0.000 | -1.711 | -0.362 |
| PING | SWU | 0.002 | 1.000 | -0.834 | 0.837 | PING | SWU | -0.528 | 0.517 | -1.311 | 0.254 |
| QTIM | SWU | 0.117 | 1.000 | -0.666 | 0.900 | QTIM | SWU | 0.509 | 0.482 | -0.228 | 1.245 |

**Supplementary Table 23. Post-hoc pairwise cohort comparisons of COMPASS across the normative cohorts.** Post-hoc pairwise comparisons of the COMPASS between all pairs of the eleven normative cohorts, following the omnibus ANOVA reported in Fig. 6. Rows denote the 55 unique cohort pairs; columns report, for the LH and RH separately, the mean difference between the two cohorts, its 95% CI, and the FDR-corrected q-value. P values are Tukey-adjusted for all 55 pairwise comparisons. Cells shaded in light orange denote p < 0.05.

| **LH** | | | | | | **RH** | | | | | |
| --- | --- | --- | --- | --- | --- | --- | --- | --- | --- | --- | --- |
| **Group1** | **Group2** | **Mean Diff** | **P**  **value** | **95% CI_low** | **95% CI_High** | **Group1** | **Group2** | **Mean Diff** | **P**  **value** | **95% CI_low** | **95% CI_High** |
| ABCD | BGSP | 8 | 0.971 | -0.545 | 0.235 | ABCD | BGSP | -0.063 | 1.000 | -0.427 | 0.301 |
| ABCD | BeijingEN | -0.641 | 0.425 | -1.536 | 0.254 | ABCD | BeijingEN | -0.401 | 0.897 | -1.232 | 0.430 |
| ABCD | CHCP | -0.162 | 1.000 | -0.921 | 0.596 | ABCD | CHCP | -0.622 | 0.140 | -1.327 | 0.082 |
| ABCD | CMI | -0.002 | 1.000 | -0.415 | 0.411 | ABCD | CMI | 0.018 | 1.000 | -0.366 | 0.402 |
| ABCD | HCP | -0.417 | 0.071 | -0.850 | 0.016 | ABCD | HCP | -0.230 | 0.744 | -0.630 | 0.170 |
| ABCD | HCP-D | -0.081 | 1.000 | -0.610 | 0.449 | ABCD | HCP-D | -0.005 | 1.000 | -0.497 | 0.487 |
| ABCD | Narratives | -0.342 | 0.866 | -1.017 | 0.334 | ABCD | Narratives | -0.021 | 1.000 | -0.648 | 0.606 |
| ABCD | PING | 0.037 | 1.000 | -0.464 | 0.537 | ABCD | PING | 0.247 | 0.827 | -0.219 | 0.712 |
| ABCD | QTIM | -0.287 | 0.592 | -0.734 | 0.159 | ABCD | QTIM | -0.105 | 0.999 | -0.523 | 0.313 |
| ABCD | SWU | -0.120 | 1.000 | -0.671 | 0.431 | ABCD | SWU | -0.113 | 1.000 | -0.632 | 0.406 |
| BGSP | BeijingEN | -0.486 | 0.816 | -1.390 | 0.419 | BGSP | BeijingEN | -0.338 | 0.968 | -1.178 | 0.502 |
| BGSP | CHCP | -0.007 | 1.000 | -0.777 | 0.763 | BGSP | CHCP | -0.559 | 0.290 | -1.274 | 0.156 |
| BGSP | CMI | 0.154 | 0.987 | -0.280 | 0.587 | BGSP | CMI | 0.081 | 1.000 | -0.322 | 0.484 |
| BGSP | HCP | -0.262 | 0.737 | -0.714 | 0.191 | BGSP | HCP | -0.167 | 0.971 | -0.586 | 0.252 |
| BGSP | HCP-D | 0.075 | 1.000 | -0.471 | 0.620 | BGSP | HCP-D | 0.058 | 1.000 | -0.449 | 0.565 |
| BGSP | Narratives | -0.186 | 0.999 | -0.874 | 0.502 | BGSP | Narratives | 0.042 | 1.000 | -0.597 | 0.681 |
| BGSP | PING | 0.192 | 0.983 | -0.326 | 0.710 | BGSP | PING | 0.310 | 0.590 | -0.172 | 0.792 |
| BGSP | QTIM | -0.132 | 0.998 | -0.598 | 0.334 | BGSP | QTIM | -0.042 | 1.000 | -0.478 | 0.394 |
| BGSP | SWU | 0.035 | 1.000 | -0.531 | 0.602 | BGSP | SWU | -0.050 | 1.000 | -0.583 | 0.484 |
| BeijingEN | CHCP | 0.479 | 0.951 | -0.636 | 1.593 | BeijingEN | CHCP | -0.221 | 1.000 | -1.255 | 0.813 |
| BeijingEN | CMI | 0.639 | 0.464 | -0.276 | 1.554 | BeijingEN | CMI | 0.419 | 0.883 | -0.430 | 1.268 |
| BeijingEN | HCP | 0.224 | 1.000 | -0.700 | 1.148 | BeijingEN | HCP | 0.171 | 1.000 | -0.685 | 1.028 |
| BeijingEN | HCP-D | 0.560 | 0.742 | -0.413 | 1.533 | BeijingEN | HCP-D | 0.396 | 0.944 | -0.507 | 1.299 |
| BeijingEN | Narratives | 0.299 | 0.998 | -0.760 | 1.359 | BeijingEN | Narratives | 0.381 | 0.976 | -0.603 | 1.364 |
| BeijingEN | PING | 0.677 | 0.444 | -0.280 | 1.635 | BeijingEN | PING | 0.648 | 0.396 | -0.241 | 1.537 |
| BeijingEN | QTIM | 0.354 | 0.979 | -0.577 | 1.284 | BeijingEN | QTIM | 0.296 | 0.990 | -0.569 | 1.161 |
| BeijingEN | SWU | 0.521 | 0.830 | -0.464 | 1.506 | BeijingEN | SWU | 0.288 | 0.995 | -0.630 | 1.206 |
| CHCP | CMI | 0.161 | 1.000 | -0.621 | 0.942 | CHCP | CMI | 0.640 | 0.141 | -0.085 | 1.366 |
| CHCP | HCP | -0.255 | 0.994 | -1.047 | 0.538 | CHCP | HCP | 0.392 | 0.820 | -0.342 | 1.127 |
| CHCP | HCP-D | 0.081 | 1.000 | -0.768 | 0.931 | CHCP | HCP-D | 0.617 | 0.288 | -0.171 | 1.405 |
| CHCP | Narratives | -0.180 | 1.000 | -1.126 | 0.767 | CHCP | Narratives | 0.602 | 0.496 | -0.277 | 1.480 |
| CHCP | PING | 0.199 | 1.000 | -0.633 | 1.030 | CHCP | PING | 0.869 | 0.013 | 0.098 | 1.641 |
| CHCP | QTIM | -0.125 | 1.000 | -0.925 | 0.675 | CHCP | QTIM | 0.517 | 0.472 | -0.227 | 1.261 |
| CHCP | SWU | 0.042 | 1.000 | -0.820 | 0.905 | CHCP | SWU | 0.509 | 0.616 | -0.296 | 1.314 |
| CMI | HCP | -0.415 | 0.146 | -0.888 | 0.057 | CMI | HCP | -0.248 | 0.757 | -0.684 | 0.189 |
| CMI | HCP-D | -0.079 | 1.000 | -0.641 | 0.483 | CMI | HCP-D | -0.023 | 1.000 | -0.545 | 0.499 |
| CMI | Narratives | -0.340 | 0.895 | -1.041 | 0.361 | CMI | Narratives | -0.039 | 1.000 | -0.689 | 0.612 |
| CMI | PING | 0.038 | 1.000 | -0.497 | 0.574 | CMI | PING | 0.229 | 0.923 | -0.268 | 0.726 |
| CMI | QTIM | -0.286 | 0.714 | -0.771 | 0.200 | CMI | QTIM | -0.123 | 0.999 | -0.576 | 0.329 |
| CMI | SWU | -0.118 | 1.000 | -0.701 | 0.464 | CMI | SWU | -0.131 | 1.000 | -0.678 | 0.416 |
| HCP | HCP-D | 0.336 | 0.728 | -0.241 | 0.913 | HCP | HCP-D | 0.225 | 0.957 | -0.309 | 0.758 |
| HCP | Narratives | 0.075 | 1.000 | -0.638 | 0.788 | HCP | Narratives | 0.209 | 0.995 | -0.451 | 0.870 |
| HCP | PING | 0.453 | 0.221 | -0.098 | 1.004 | HCP | PING | 0.477 | 0.090 | -0.033 | 0.986 |
| HCP | QTIM | 0.130 | 0.999 | -0.373 | 0.632 | HCP | QTIM | 0.125 | 0.999 | -0.342 | 0.591 |
| HCP | SWU | 0.297 | 0.878 | -0.300 | 0.894 | HCP | SWU | 0.117 | 1.000 | -0.442 | 0.676 |
| HCP-D | Narratives | -0.261 | 0.992 | -1.037 | 0.515 | HCP-D | Narratives | -0.015 | 1.000 | -0.735 | 0.704 |
| HCP-D | PING | 0.117 | 1.000 | -0.512 | 0.747 | HCP-D | PING | 0.252 | 0.949 | -0.332 | 0.837 |
| HCP-D | QTIM | -0.207 | 0.988 | -0.794 | 0.381 | HCP-D | QTIM | -0.100 | 1.000 | -0.647 | 0.447 |
| HCP-D | SWU | -0.039 | 1.000 | -0.709 | 0.631 | HCP-D | SWU | -0.108 | 1.000 | -0.735 | 0.520 |
| Narratives | PING | 0.378 | 0.874 | -0.378 | 1.135 | Narratives | PING | 0.268 | 0.978 | -0.434 | 0.969 |
| Narratives | QTIM | 0.055 | 1.000 | -0.667 | 0.776 | Narratives | QTIM | -0.085 | 1.000 | -0.756 | 0.587 |
| Narratives | SWU | 0.222 | 0.998 | -0.568 | 1.012 | Narratives | SWU | -0.092 | 1.000 | -0.830 | 0.646 |
| PING | QTIM | -0.324 | 0.740 | -0.886 | 0.238 | PING | QTIM | -0.352 | 0.523 | -0.876 | 0.171 |
| PING | SWU | -0.157 | 1.000 | -0.804 | 0.491 | PING | SWU | -0.360 | 0.706 | -0.967 | 0.247 |
| QTIM | SWU | 0.167 | 0.998 | -0.439 | 0.774 | QTIM | SWU | -0.008 | 1.000 | -0.579 | 0.564 |

**Supplementary Table 24. Cohort effects on anomaly scores before and after adjustment for demographics and surface quality.** Omnibus tests of between-cohort differences in each anomaly score across the normative cohorts, under three progressively adjusted model specifications. Rows are grouped by hemisphere (left, then right), each listing the three anomaly scores (reconstruction error, GNN logit, COMPASS_1). Columns report, for each specification, the F statistic, its p-value, and the FDR-corrected q-value: the unadjusted one-way ANOVA (ANOVA); an ANCOVA adjusting for age and sex (ANCOVA_1); and an ANCOVA additionally adjusting for the surface-quality proxy (ANCOVA_2, adding the negative-log-transformed FreeSurfer Euler number). Cohort effects were evaluated by type II sums of squares, and FDR correction (Benjamini–Hochberg) was applied across the three anomaly scores within each hemisphere and model specification. A cohort effect that is significant in the unadjusted ANOVA but non-significant after adjustment indicates that the apparent between-cohort variance is attributable to the adjusted covariates rather than to acquisition-related batch effects, except for RH Recon Error.

|  | **ANOVA** | | | **ANCOVA_1** | | | **ANOCVA_2** | | |
| --- | --- | --- | --- | --- | --- | --- | --- | --- | --- |
| **Metric** | **F** | **P value** | **FDR q** | **F** | **P value** | **FDR q** | **F** | **P value** | **FDR q** |
| LH Recon Error | 4.794 | 0.000 | <0.001*** | 0.843 | 0.587 | 0.825 | 1.299 | 0.228 | 0.575 |
| LH GNN Logit | 0.880 | 0.551 | 0.551 | 0.934 | 0.501 | 0.825 | 0.953 | 0.484 | 0.575 |
| LH COMPASS | 2.005 | 0.031 | 0.047* | 0.587 | 0.825 | 0.825 | 0.856 | 0.575 | 0.575 |
| RH Recon Error | 5.272 | 0.000 | <0.001*** | 3.222 | 0.001 | 0.002** | 3.482 | 0.000 | 0.001** |
| RH GNN Logit | 1.699 | 0.078 | 0.078 | 1.690 | 0.080 | 0.120 | 0.780 | 0.649 | 0.649 |
| RH COMPASS | 2.133 | 0.021 | 0.031* | 1.144 | 0.328 | 0.328 | 1.193 | 0.293 | 0.440 |

**Supplementary Table 25. Case-versus-control discrimination performance (AUROC) across methods and cohorts.** Group-separation performance of each method, quantified by the AUROC between each patient group and its healthy controls on global anomaly scores. Rows denote the patient groups and an average among groups, columns denote the six methods compared, reported separately for LH and RH. Each entry is the mean AUROC ± SD over 1,000 bootstrap resamples. To assess reproducibility, we trained four independent models with different random seeds and validation folds. For each patient group, the best-performing method is shown in **bold** and the second-best is underlined; COMPASS_2 is excluded from this ranking because its fusion weight is selected in-sample to maximize case-control separation, making its AUROC an optimistic upper bound rather than a comparable operating point. SGM likewise carries an in-sample reference advantage and should be read as a favorable-case reference.

| **Seed 1**  **(Fig. 6)** | **LH** | **SGM** | **OC-SVM** | **Recon Error** | **GNN Logit** | **COMPASS_1** | ***COMPASS_2*** |
| --- | --- | --- | --- | --- | --- | --- | --- |
|  | SV | 0.623 ± 0.040 | 0.503 ± 0.042 | **0.735 ± 0.035** | 0.589 ± 0.039 | 0.708 ± 0.036 | *0.739 ± 0.035* |
|  | TGA | 0.509 ± 0.044 | 0.447 ± 0.043 | 0.642 ± 0.042 | 0.673 ± 0.041 | **0.681 ± 0.040** | *0.686 ± 0.040* |
|  | ToF | 0.600 ± 0.054 | 0.478 ± 0.058 | 0.622 ± 0.060 | 0.659 ± 0.052 | **0.670 ± 0.057** | *0.674 ± 0.057* |
|  | PMG | **0.947 ± 0.035** | 0.190 ± 0.072 | 0.729 ± 0.085 | 0.741 ± 0.083 | 0.784 ± 0.081 | *0.795 ± 0.078* |
|  | **Average** | 0.672 ± 0.057 | 0.420 ± 0.068 | 0.659 ± 0.069 | 0.669 ± 0.065 | **0.712 ± 0.065** | *0.727 ± 0.063* |
|  | **RH** | **SGM** | **OC-SVM** | **Recon Error** | **GNN Logit** | **COMPASS_1** | ***COMPASS_2*** |
|  | SV | **0.620 ± 0.038** | 0.505 ± 0.041 | 0.579 ± 0.041 | 0.561 ± 0.039 | 0.591 ± 0.040 | *0.592 ± 0.041* |
|  | TGA | 0.555 ± 0.044 | 0.464 ± 0.042 | 0.604 ± 0.040 | 0.585 ± 0.043 | **0.629 ± 0.042** | *0.630 ± 0.042* |
|  | ToF | 0.654 ± 0.054 | 0.457 ± 0.055 | 0.589 ± 0.050 | **0.678 ± 0.048** | 0.667 ± 0.049 | *0.687 ± 0.048* |
|  | PMG | **0.904 ± 0.050** | 0.686 ± 0.084 | 0.763 ± 0.075 | 0.748 ± 0.077 | 0.786 ± 0.079 | *0.798 ± 0.075* |
|  | **Average** | **0.686 ± 0.059** | 0.560 ± 0.067 | 0.638 ± 0.065 | 0.637 ± 0.064 | 0.677 ± 0.065 | *0.688 ± 0.063* |
| **Seed 2** | **LH** | **SGM** | **OC-SVM** | **Recon Error** | **GNN Logit** | **COMPASS_1** | ***COMPASS_2*** |
|  | SV | 0.615 ± 0.038 | 0.488 ± 0.040 | **0.662 ± 0.036** | 0.535 ± 0.040 | 0.628 ± 0.038 | 0.662 ± 0.036 |
|  | TGA | 0.502 ± 0.043 | 0.482 ± 0.044 | 0.650 ± 0.042 | 0.616 ± 0.041 | **0.660 ± 0.040** | 0.661 ± 0.040 |
|  | ToF | 0.595 ± 0.053 | 0.512 ± 0.055 | 0.592 ± 0.060 | 0.588 ± 0.053 | **0.623 ± 0.055** | 0.623 ± 0.055 |
|  | PMG | **0.947 ± 0.035** | 0.288 ± 0.089 | 0.757 ± 0.083 | 0.760 ± 0.087 | 0.796 ± 0.076 | 0.804 ± 0.076 |
|  | **Average** | 0.663 ± 0.057 | 0.490 ± 0.067 | 0.645 ± 0.068 | 0.647 ± 0.064 | **0.684 ± 0.063** | 0.700 ± 0.062 |
|  | **RH** | **SGM** | **OC-SVM** | **Recon Error** | **GNN Logit** | **COMPASS_1** | ***COMPASS_2*** |
|  | SV | **0.623 ± 0.039** | 0.559 ± 0.043 | 0.591 ± 0.041 | 0.581 ± 0.040 | 0.606 ± 0.041 | 0.610 ± 0.041 |
|  | TGA | 0.556 ± 0.043 | 0.451 ± 0.044 | 0.599 ± 0.042 | 0.598 ± 0.042 | **0.621 ± 0.042** | 0.622 ± 0.042 |
|  | ToF | 0.654 ± 0.054 | 0.476 ± 0.055 | 0.622 ± 0.048 | 0.645 ± 0.051 | **0.673 ± 0.049** | 0.676 ± 0.049 |
|  | PMG | **0.901 ± 0.050** | 0.271 ± 0.081 | 0.739 ± 0.083 | 0.677 ± 0.090 | 0.752 ± 0.082 | 0.791 ± 0.079 |
|  | **Average** | **0.687 ± 0.059** | 0.429 ± 0.067 | 0.634 ± 0.066 | 0.610 ± 0.068 | 0.649 ± 0.066 | 0.672 ± 0.064 |
| **Seed 3** | **LH** | **SGM** | **OC-SVM** | **Recon Error** | **GNN Logit** | **COMPASS_1** | ***COMPASS_2*** |
|  | SV | 0.615 ± 0.038 | 0.449 ± 0.041 | **0.650 ± 0.037** | 0.533 ± 0.039 | 0.626 ± 0.037 | 0.652 ± 0.037 |
|  | TGA | 0.502 ± 0.043 | 0.387 ± 0.043 | 0.613 ± 0.042 | 0.622 ± 0.043 | **0.648 ± 0.042** | 0.650 ± 0.042 |
|  | ToF | 0.595 ± 0.053 | 0.442 ± 0.056 | 0.609 ± 0.056 | 0.648 ± 0.051 | **0.654 ± 0.054** | 0.663 ± 0.053 |
|  | PMG | **0.947 ± 0.035** | 0.229 ± 0.077 | 0.758 ± 0.082 | 0.694 ± 0.088 | 0.751 ± 0.080 | 0.769 ± 0.082 |
|  | **Average** | 0.667 ± 0.056 | 0.384 ± 0.069 | 0.650 ± 0.067 | 0.639 ± 0.065 | **0.681 ± 0.064** | 0.692 ± 0.064 |
|  | **RH** | **SGM** | **OC-SVM** | **Recon Error** | **GNN Logit** | **COMPASS_1** | ***COMPASS_2*** |
|  | SV | **0.623 ± 0.039** | 0.473 ± 0.039 | 0.608 ± 0.040 | 0.583 ± 0.039 | 0.617 ± 0.039 | 0.621 ± 0.039 |
|  | TGA | 0.556 ± 0.043 | 0.498 ± 0.043 | 0.589 ± 0.042 | 0.625 ± 0.041 | **0.631 ± 0.042** | 0.644 ± 0.041 |
|  | ToF | 0.654 ± 0.054 | 0.501 ± 0.057 | 0.616 ± 0.051 | 0.666 ± 0.050 | **0.681 ± 0.047** | 0.685 ± 0.048 |
|  | PMG | **0.890 ± 0.052** | 0.257 ± 0.081 | 0.712 ± 0.087 | 0.691 ± 0.087 | 0.726 ± 0.084 | 0.733 ± 0.084 |
|  | **Average** | **0.684 ± 0.060** | 0.428 ± 0.067 | 0.627 ± 0.068 | 0.603 ± 0.067 | 0.638 ± 0.066 | 0.662 ± 0.066 |
| **Seed 4** | **LH** | **SGM** | **OC-SVM** | **Recon Error** | **GNN Logit** | **COMPASS_1** | ***COMPASS_2*** |
|  | SV | 0.615 ± 0.038 | 0.489 ± 0.040 | **0.680 ± 0.036** | 0.533 ± 0.039 | 0.643 ± 0.036 | 0.681 ± 0.035 |
|  | TGA | 0.502 ± 0.043 | 0.437 ± 0.041 | 0.645 ± 0.041 | 0.633 ± 0.043 | **0.659 ± 0.041** | 0.659 ± 0.041 |
|  | ToF | 0.595 ± 0.053 | 0.453 ± 0.055 | 0.619 ± 0.056 | 0.672 ± 0.048 | **0.683 ± 0.053** | 0.689 ± 0.051 |
|  | PMG | **0.947 ± 0.035** | 0.141 ± 0.060 | 0.643 ± 0.091 | 0.683 ± 0.087 | 0.703 ± 0.085 | 0.703 ± 0.085 |
|  | **Average** | 0.663 ± 0.057 | 0.434 ± 0.062 | 0.609 ± 0.068 | 0.650 ± 0.066 | **0.665 ± 0.066** | 0.692 ± 0.064 |
|  | **RH** | **SGM** | **OC-SVM** | **Recon Error** | **GNN Logit** | **COMPASS_1** | ***COMPASS_2*** |
|  | SV | **0.623 ± 0.039** | 0.473 ± 0.039 | 0.609 ± 0.040 | 0.580 ± 0.039 | 0.617 ± 0.039 | 0.619 ± 0.039 |
|  | TGA | 0.556 ± 0.043 | 0.498 ± 0.043 | 0.589 ± 0.042 | 0.627 ± 0.041 | **0.631 ± 0.042** | 0.645 ± 0.041 |
|  | ToF | 0.654 ± 0.054 | 0.501 ± 0.057 | 0.617 ± 0.051 | 0.667 ± 0.050 | **0.676 ± 0.047** | 0.686 ± 0.047 |
|  | PMG | **0.890 ± 0.052** | 0.257 ± 0.081 | 0.712 ± 0.087 | 0.716 ± 0.084 | 0.720 ± 0.085 | 0.744 ± 0.083 |
|  | **Average** | **0.684 ± 0.060** | 0.428 ± 0.067 | 0.628 ± 0.068 | 0.608 ± 0.067 | 0.639 ± 0.066 | 0.666 ± 0.066 |

**Supplementary Table 26. Baseline comparison of associations between anomaly scores and clinical outcomes.** Associations between each anomaly score and clinical outcomes, compared across methods for the validation cohorts using the simplest unadjusted statistic appropriate to each: Pearson correlation for the continuous CHD outcomes, and the KW test for the ordinal PMG language outcome. Rows denote the region-outcome pairs; only pairs reaching significance for at least one method are shown. Columns denote the six methods compared. Each cell reports the association statistic (Pearson $r$ for CHD, the H statistic for PMG) and its FDR-corrected q-value. For each row, the strongest association is shown in **bold** and the second-strongest underlined, ranked by effect-size magnitude ($|r|$, or $H$). These unadjusted statistics are intended for method-to-method comparison and differ by design from the covariate-adjusted estimates in the main analyses; SGM carries an in-sample reference advantage and should be read as a favorable-case reference.

| **Cohort** | **Region-Outcome Pair** | **SGM** | **OC-SVM** | **Recon Error** | **GNN Logit** | **COMPASS_1** | **COMPASS_2** |
| --- | --- | --- | --- | --- | --- | --- | --- |
| CHD | LF ×  General Memory | -0.075 (0.169) | 0.040 (0.469) | **-0.172** (0.002)** | -0.081 (0.141) | -0.160** (0.003) | -0.145** (0.008) |
|  | LF ×  Processing Speed | -0.015 (0.790) | 0.038 (0.501) | -0.204*** (0.000) | -0.135* (0.015) | **-0.216*** (0.000)** | -0.203*** (0.000) |
|  | LH ×  Processing Speed | -0.080 (0.151) | 0.056 (0.313) | **-0.214*** (0.000)** | -0.089 (0.110) | -0.188** (0.001) | -0.212*** (0.000) |
|  | LP ×  Verbal Comprehension | **-0.134* (0.017)** | 0.130* (0.020) | -0.037 (0.511) | -0.061 (0.277) | -0.063 (0.263) | -0.050 (0.369) |
|  | RF ×  Full-Scale IQ | -0.138* (0.013) | 0.086 (0.122) | -0.104 (0.063) | -0.13* (0.02) | **-0.160** (0.004)** | **-0.160** (0.004)** |
|  | RF ×  Processing Speed | -0.102 (0.066) | 0.107 (0.054) | -0.085 (0.128) | -0.171** (0.002) | -0.178** (0.001) | **-0.182** (0.001)** |
|  | RT ×  Executive Function | -0.147** (0.007) | 0.000 (0.994) | -0.103 (0.059) | -0.135* (0.013) | **-0.156** (0.004)** | -0.145** (0.008) |
| PMG | LF ×  Language | 1.545 (0.462) | 0.645 (0.724) | 5.526 (0.063) | 9.730** (0.008) | **9.905** (0.007)** | **9.905** (0.007)** |
|  | LH ×  Language | 1.830 (0.400) | 4.632 (0.099) | 4.080 (0.130) | 7.164* (0.028) | **8.155* (0.017)** | **8.155* (0.017)** |
|  | LP ×  Language | 4.716 (0.095) | 0.289 (0.865) | 5.651 (0.059) | 1.344 (0.511) | 5.506 (0.064) | **7.098* (0.029)** |
|  | RF ×  Language | 5.997 (0.050) | 0.645 (0.724) | 6.013* (0.049) | 9.730** (0.008) | **9.905** (0.007)** | **9.905** (0.007)** |

**Supplementary Table 27. Summary of the normative training cohorts.** Composition of the normative training dataset by cohort and hemisphere. For each hemisphere, rows list the cohorts followed by a total row. Columns report the original number of subjects with available T1-weighted scans (Original N), the number removed during MRI and surface quality control (QC Excluded N), the resulting analyzed sample (Final N), the mean ± SD of age, the age range, the numbers of male and female subjects, and the mean ± SD of surface quality proxy (Euler number). Quality control comprised visual inspection of reconstructed surfaces and exclusion of failed or low-quality reconstructions (Supplementary Fig. 3). Subjects with missing MRI scans or failed FreeSurfer processing (total N=51; see Supplementary Fig. 3 top row, second column) were excluded from this table. Age is given in years. To prevent overfitting to specific age range and to balance the age distribution across the total training dataset, we employed a sex-balanced, random subset of ABCD (1,000 males and 1,000 females; 8 out of them were excluded due to the low-quality reconstructions).

| **LH** | **Cohort** | **Original N** | **QC Excluded N** | **Final N** | **Age (Mean ± SD)** | **Age Range** | **Males** | **Females** | **Euler number** |
| --- | --- | --- | --- | --- | --- | --- | --- | --- | --- |
|  | HCP | 1102 | 3 | 1099 | 28.8 ± 3.7 | 22.0 - 37.0 | 502 | 597 | -27.4 ± 10.8 |
|  | HCPD | 646 | 41 | 605 | 14.0 ± 3.8 | 8.0 - 21.9 | 298 | 307 | -33.5 ± 17.9 |
|  | CHCP | 260 | 0 | 260 | 23.2 ± 4.1 | 18.0 - 39.0 | 126 | 134 | -21.5 ± 9.6 |
|  | CMI | 1692 | 792 | 900 | 11.1 ± 3.7 | 5.0 - 21.9 | 512 | 388 | -38.8 ± 32.4 |
|  | BeijingEN | 179 | 0 | 179 | 21.2 ± 1.9 | 17.0 - 28.0 | 72 | 107 | -21.1 ± 10.2 |
|  | BGSP | 1570 | 11 | 1559 | 21.5 ± 2.9 | 19.0 - 35.0 | 661 | 898 | -20.7 ± 10.4 |
|  | PING | 755 | 33 | 722 | 12.5 ± 5.1 | 3.0 - 21.0 | 379 | 343 | -35.0 ± 17.1 |
|  | QTIM | 1198 | 196 | 1002 | 21.6 ± 3.7 | 12.0 - 30.0 | 383 | 619 | -79.3 ± 24.1 |
|  | SWU | 565 | 12 | 553 | 20.1 ± 1.3 | 17.0 - 27.0 | 241 | 312 | -49.0 ± 22.7 |
|  | Narratives | 339 | 2 | 337 | 21.5 ± 3.5 | 18.0 - 38.0 | 135 | 202 | -23.6 ± 11.2 |
|  | ABCD* | 1992 | 26 | 1966 | 9.9 ± 0.6 | 8.9 - 11.1 | 984 | 982 | -21.4 ± 11.5 |
|  | TOTAL | 10298 | 1116 | 9182 | - | - | 4293 | 4889 | -33.7 ± 25.3 |
| **RH** | **Cohort** | **Original N** | **QC Excluded N** | **Final N** | **Age (Mean ± SD)** | **Age Range** | **Males** | **Females** | **Euler number** |
|  | HCP | 1102 | 0 | 1102 | 28.8 ± 3.7 | 22.0 - 37.0 | 502 | 600 | -28.3 ± 11.9 |
|  | HCPD | 646 | 27 | 619 | 13.9 ± 3.8 | 8.0 - 21.9 | 308 | 311 | -28.4 ± 15.0 |
|  | CHCP | 260 | 0 | 260 | 23.2 ± 4.1 | 18.0 - 39.0 | 126 | 134 | -21.9 ± 9.9 |
|  | CMI | 1692 | 816 | 876 | 11.1 ± 3.7 | 5.0 - 21.9 | 502 | 374 | -34.7 ± 28.0 |
|  | BeijingEN | 179 | 0 | 179 | 21.2 ± 1.9 | 17.0 - 28.0 | 72 | 107 | -19.1 ± 8.7 |
|  | BGSP | 1570 | 11 | 1559 | 21.5 ± 2.9 | 19.0 - 35.0 | 659 | 900 | -18.3 ± 9.3 |
|  | PING | 755 | 33 | 722 | 12.5 ± 5.0 | 3.0 - 21.0 | 374 | 348 | -33.0 ± 15.1 |
|  | QTIM | 1198 | 230 | 968 | 21.7 ± 3.6 | 12.0 - 30.0 | 360 | 608 | -69.5 ± 22.3 |
|  | SWU | 565 | 30 | 535 | 20.1 ± 1.3 | 17.0 - 27.0 | 225 | 310 | -47.6 ± 22.0 |
|  | Narratives | 339 | 2 | 337 | 21.5 ± 3.5 | 18.0 - 38.0 | 136 | 201 | -20.9 ± 9.8 |
|  | ABCD* | 1992 | 43 | 1949 | 9.9 ± 0.6 | 8.9 - 11.1 | 973 | 976 | -19.2 ± 11.0 |
|  | TOTAL | 10298 | 1192 | 9106 | - | - | 4237 | 4869 | -30.7 ± 22.6 |

**Supplementary Table 29. Demographic and quality-control summary of the CHD cohorts.** Composition of the CHD cohort by hemisphere. For each hemisphere, rows list the subgroups followed by a total row. Columns report the original number of subjects with available T1-weighted scans (Original N), the number removed during MRI and surface quality control (QC Excluded N), the resulting analyzed sample (Final N), the mean ± SD of age, the age range, the numbers of male and female subjects and the mean ± SD of SES and Euler number. Quality control comprised visual inspection of reconstructed surfaces and exclusion of failed or low-quality reconstructions (Supplementary Fig. 3). Age is given in years.

| **LH** | **Group** | **Original N** | **QC Excluded N** | **Final N** | **Age**  **(Mean ± SD)** | **Age Range** | **Males** | **Females** | **SES**  **(Mean ± SD)** | **Euler number** |
| --- | --- | --- | --- | --- | --- | --- | --- | --- | --- | --- |
|  | Control | 94 | 2 | 92 | 15.4 ± 1.9 | 10.1 - 19.7 | 45 | 47 | 53.3 ± 9.9 | -38.5 ± 17.1 |
|  | SV | 115 | 0 | 115 | 14.8 ± 3.0 | 10.2 - 20.0 | 67 | 48 | 48.9 ± 13.8 | -43.1 ± 20.5 |
|  | TGA | 91 | 0 | 91 | 16.2 ± 0.7 | 14.9 - 19.9 | 70 | 21 | 47.1 ± 12.5 | -42.3 ± 17.7 |
|  | ToF | 41 | 0 | 41 | 14.8 ± 1.0 | 13.3 - 17.0 | 27 | 14 | 50.4 ± 10.9 | -34.6 ± 18.6 |
|  | TOTAL | 341 | 2 | 339 | 15.3 ± 2.1 | 10.1 - 20.0 | 209 | 130 | 49.8 ± 12.3 | -40.6 ± 18.8 |
| **RH** | **Group** | **Original N** | **QC Excluded N** | **Final N** | **Age (Mean ± SD)** | **Age Range** | **Males** | **Females** | **SES**  **(Mean ± SD)** | **Euler number** |
|  | Control | 94 | 0 | 94 | 15.4 ± 1.9 | 10.1 - 19.7 | 46 | 48 | 53.4 ± 9.9 | -39.7 ± 21.5 |
|  | SV | 115 | 0 | 115 | 14.8 ± 3.0 | 10.2 - 20.0 | 67 | 48 | 48.9 ± 13.8 | -40.0 ± 21.9 |
|  | TGA | 91 | 0 | 91 | 16.2 ± 0.7 | 14.9 - 19.9 | 70 | 21 | 47.1 ± 12.5 | -41.5 ± 16.5 |
|  | ToF | 41 | 0 | 41 | 14.8 ± 1.0 | 13.3 - 17.0 | 27 | 14 | 50.4 ± 10.9 | -36.7 ± 15.2 |
|  | TOTAL | 341 | 0 | 341 | 15.3 ± 2.1 | 10.1 - 20.0 | 210 | 131 | 49.8 ± 12.3 | -39.9 ± 19.7 |

**Supplementary Table 30. Neurodevelopmental outcome scores by group in the CHD cohort.** Distribution of neurodevelopmental outcome scores across the CHD subgroups. Rows denote the eight assessed outcomes; columns are grouped by subgroup, each reporting the number of subjects with an available score (N), the score range, and the mean ± SD. Cognitive indices (full-scale IQ, verbal comprehension, perceptual reasoning, general memory, processing speed) were derived from the age-appropriate Wechsler scale, executive function from the average of DKEFS subtests, and math and reading composites from WIAT.

| **Outcome** | **Control** | | | **SV** | | | **TGA** | | | **ToF** | | |
| --- | --- | --- | --- | --- | --- | --- | --- | --- | --- | --- | --- | --- |
|  | **N** | **Range** | **Mean ± SD** | **N** | **Range** | **Mean ± SD** | **N** | **Range** | **Mean ± SD** | **N** | **Range** | **Mean ± SD** |
| Executive Function | 91 | 5.7 - 14.5 | 10.8 ± 1.6 | 113 | 1.5 - 13.8 | 8.6 ± 2.3 | 92 | 3.5 - 14.0 | 9.4 ± 2.0 | 39 | 3.4 - 13.5 | 9.0 ± 2.5 |
| Full-Scale IQ | 90 | 76.0 - 141.0 | 108.0 ± 11.4 | 114 | 55.0 - 132.0 | 92.0 ± 16.6 | 79 | 64.0 - 138.0 | 99.5 ± 15.5 | 40 | 48.0 - 126.0 | 96.4 ± 19.8 |
| Verbal Comprehension | 90 | 83.0 - 138.0 | 110.1 ± 11.7 | 114 | 55.0 - 130.0 | 98.1 ± 15.4 | 79 | 68.0 - 150.0 | 102.9 ± 16.4 | 40 | 55.0 - 134.0 | 102.2 ± 19.6 |
| Perceptual Reasoning | 90 | 69.0 - 137.0 | 105.5 ± 12.2 | 114 | 58.0 - 127.0 | 93.1 ± 15.8 | 79 | 62.0 - 136.0 | 96.7 ± 14.5 | 41 | 49.0 - 135.0 | 94.6 ± 17.2 |
| General Memory | 90 | 63.0 - 136.0 | 103.7 ± 15.1 | 114 | 54.0 - 129.0 | 91.9 ± 15.9 | 92 | 50.0 - 136.0 | 91.9 ± 18.0 | 40 | 50.0 - 130.0 | 93.8 ± 21.4 |
| Processing Speed | 90 | 73.0 - 140.0 | 102.5 ± 13.4 | 114 | 50.0 - 133.0 | 88.2 ± 16.5 | 79 | 67.0 - 129.0 | 100.1 ± 14.2 | 41 | 56.0 - 136.0 | 94.1 ± 17.7 |
| Math Composite | 92 | 82.0 - 128.0 | 106.4 ± 10.4 | 113 | 40.0 - 122.0 | 91.3 ± 17.1 | 92 | 56.0 - 130.0 | 98.4 ± 15.3 | 41 | 42.0 - 127.0 | 99.2 ± 16.8 |
| Reading Composite | 92 | 70.0 - 138.0 | 111.8 ± 13.9 | 113 | 40.0 - 141.0 | 92.4 ± 23.4 | 92 | 40.0 - 127.0 | 99.1 ± 18.8 | 41 | 40.0 - 129.0 | 96.5 ± 24.2 |

**Supplementary Table 31. Demographic and quality-control summary of the PMG cohorts.** Composition of the PMG cohort by hemisphere. For each hemisphere, rows list the subgroups followed by a total row. Columns report the original number of subjects with available T1-weighted scans (Original N), the number removed during MRI and surface quality control (QC Excluded N), the resulting analyzed sample (Final N), the mean ± SD of age, the age range, the numbers of male and female subjects and the mean ± SD of Euler number. Quality control comprised visual inspection of reconstructed surfaces and exclusion of failed or low-quality reconstructions (Supplementary Fig. 3). Age is given in years.

| **LH** | **Group** | **Original N** | **QC Excluded N** | **Final N** | **Age (Mean ± SD)** | **Age Range** | **Males** | **Females** | **Euler (Mean ± SD)** |
| --- | --- | --- | --- | --- | --- | --- | --- | --- | --- |
|  | CON | 26 | 5 | 21 | 9.7 ± 5.0 | 2.0 - 18.0 | 8 | 13 | -47.0 ± 26.2 |
|  | PMG | 18 | 0 | 18 | 8.8 ± 5.3 | 2.0 - 18.0 | 12 | 6 | -54.7 ± 26.6 |
|  | TOTAL | 44 | 5 | 39 | 9.3 ± 5.1 | 2.0 - 18.0 | 20 | 19 | -50.5 ± 26.3 |
| **RH** | **Group** | **Original N** | **QC Excluded N** | **Final N** | **Age (Mean ± SD)** | **Age Range** | **Males** | **Females** | **Euler (Mean ± SD)** |
|  | CON | 26 | 4 | 22 | 9.4 ± 5.1 | 2.0 - 18.0 | 8 | 14 | -43.7 ± 24.7 |
|  | PMG | 18 | 0 | 18 | 8.8 ± 5.3 | 2.0 - 18.0 | 12 | 6 | -45.8 ± 17.4 |
|  | TOTAL | 44 | 4 | 40 | 9.1 ± 5.1 | 2.0 - 18.0 | 20 | 20 | -44.6 ± 21.5 |
